# Parallel neuronal ensembles control behavior across sensorimotor levels in *Drosophila*

**DOI:** 10.64898/2025.12.13.693955

**Authors:** Sander Liessem, Samuel K. Asinof, Aljoscha Nern, Marissa Sumathipala, Edward Rogers, Mert Erginkaya, Chris J. Dallmann, Gwyneth M. Card, Jan M. Ache

**Author notes:** Correspondence: Gwyneth M. Card, Jan Marek Ache. Equal first authors (Sander Liessem, Samuel K. Asinof).

## Abstract

Nervous systems can process information in serial or in parallel, trading off efficiency for flexibility and speed. How these network architectures are implemented across sensorimotor pathways to control behavior is unclear. We investigate this tradeoff directly in *Drosophila* by comparing neuronal circuits underlying landing and takeoff, behaviors transforming similar visual cues to whole-body motor output. Using a whole-CNS connectome, electrophysiology, and behavioral analysis, we reconstruct the complete feedforward pathway for landing, including visual feature detectors, a dedicated ensemble of descending neurons (DNs), and a core premotor circuit in the nerve cord. Comparison to the takeoff pathway reveals that, despite encoding the same sensory feature and engaging similar muscle groups, neuronal circuits controlling the two behaviors are separated at every sensorimotor level. Extending this analysis to the complete DN population reveals a blueprint for descending motor control: DNs across the behavioral space utilized by the fly are organized as a set of parallel, loosely-overlapping ensembles that form a continuum from command-like control, with individual DNs determining behavioral output, to population coding, with multiple DNs controlling behavior synergistically. Distinct combinations of sensory feature detectors differentially recruit DN ensembles to enable flexible, context-dependent behavioral control.

## Introduction

Behavior arises from the coordinated activity of interconnected neuronal networks processing input from sensory neurons, integrating it with internal states, and providing output to motor neurons (MNs) controlling the body’s muscles. These networks may be organized serially, such that information for multiple behaviors is processed within a single pathway sequentially through the sensorimotor stages of sensory detection, feature extraction, decision making, and motor planning (Arber & Costa, 2022; Eaton et al., 2001; Felleman & Van Essen, 1991; Korn & Faber, 2005; Riesenhuber & Poggio, 1999). Alternatively, the networks may be divided into separate circuits for each behavior that process information simultaneously in parallel (Arber & Costa, 2022; Bullmore & Sporns, 2012; Chen et al., 2017; Doudlah et al., 2022; Griffa et al., 2023; Kaiser & Hilgetag, 2006; Slangewal et al., 2025). Depending on the task and context, it may be advantageous for the central nervous system (CNS) to use one or the other of these architectures. Serial architectures minimize wiring and neuron number but are slower because they suffer from bottlenecks in processing, whereas parallel architectures allow rapid, scalable processing at the cost of duplicating computations and require more connections and coordination across circuits. Different sensorimotor processing levels may also employ different architectures. For example, parallel architectures often occur in peripheral sensory circuits, such as the early visual system, that process different features of an object such as shape, motion, and color separately (Klapoetke et al., 2022; Longden et al., 2023; Städele et al., 2020; Turner et al., 2022). Serial architectures are primarily associated with cognitive tasks, such as decision-making, and evidence from response times indicates that there are structural bottlenecks along the process of choosing a response to a sensory cue (Kang et al., 2021; Sigman & Dehaene, 2005).

Animals achieve sensorimotor integration with brains of different sizes and complexity. For instance, the nematode *C. elegans* relies on 302-385 multifunctional neurons to control its entire behavioral output (Huang et al., 2023), the CNS of the fruit fly *Drosophila* features ∼165,000 neurons (Bates et al., 2025; Berg et al., 2025), and the mouse brain contains more than 70,000,000 neurons (Herculano-Houzel et al., 2006). Despite their potentially lower computational power, animals with small nervous systems integrate complex, multimodal sensory stimuli and display an impressive range of behaviors (Büschges & Ache, 2025). Animals with numerically smaller brains might need to rely on serial processing or use individual neurons for multiple tasks in sensorimotor integration (multiplexing), since they have limited room for redundancy (Atanas et al., 2023). Vertebrates, on the other hand, are equipped with larger numbers of neurons, which could allow them to more extensively distribute computations across parallel networks.

In *Drosophila*, we can now directly address questions of network architecture at scale, as sensorimotor pathways can be mapped at synaptic resolution in connectomes of the entire CNS (Bates et al., 2025; Berg et al., 2025). Because the fly must coordinate complex whole-body movements with a compact CNS, one might expect efficiency to favor serial architectures that minimize redundant circuitry, particularly for behaviors that share a sensory stimulus space and target the same muscle groups. Landing and escape takeoff are two behaviors triggered by the same looming stimulus, which diverge into distinct motor patterns requiring movements of the same body parts (Ache et al., 2019a; Ache et al., 2019b). Thus, they provide an excellent starting point to determine whether sensorimotor pathways for related behaviors are routed through shared serial circuits or distinct parallel pathways. Understanding whether these behaviors share circuitry requires defining the neuronal circuit elements of each pathway. For the takeoff pathway, neuronal cell types detecting looming (Ache et al., 2019b; Klapoetke et al., 2017; Klapoetke et al., 2022; von Reyn et al., 2017; Wu et al., 2016), DNs driving takeoff and associated preparatory actions (Dombrovski et al., 2023; von Reyn et al., 2014), and pre-motor neurons in the VNC (Azevedo et al., 2024; Cheong et al., 2025) have been identified. In contrast, the landing pathway is far less resolved with only two DN types (DNp07 and DNp10) known to drive the landing motor pattern when activated (Ache et al., 2019a). Neither of the identified DN types overlap with known takeoff DNs, suggesting that the pathways for takeoff and landing are arranged in parallel at the level of the DNs. However, it is likely that flies use more than two DNs to control landing. Furthermore, it remains unclear whether landing and takeoff circuits overlap at other sensorimotor processing levels. They might, for example, use the same sensory circuits for looming detection or converge onto shared motor pathways. Previously, we showed that visual stimuli evoke activity in landing DNs only during flight (Ache et al., 2019a), suggesting a shared sensory feature could be routed into one or the other of the two motor pathways depending on the behavioral state to enable wiring economy through serial organization (Maimon, 2011; McGinley et al., 2015).

Here we first apply connectomic and experimental tools available in *Drosophila* to determine and characterize the neuronal elements, from sensory networks in the visual periphery to motor networks in the VNC, that control the fly’s landing and takeoff behaviors. We then analyze the organization of the landing and takeoff pathways at three key layers across the CNS: sensory feature detection, descending control, and pre-motor networks. This yields the surprising finding that the circuits underlying the two behaviors are virtually non-overlapping. By comparing circuit architectures in the novel whole-CNS connectome, maleCNS (Berg et al., 2025), to the same circuits in previously published connectomes of the brain (Dorkenwald et al., 2024; Schlegel et al., 2024; Zheng et al., 2018) and VNC (Cheong et al., 2025; Marin et al., 2024; Stürner et al., 2024; Takemura et al., 2024), we confirm the robustness of our connectome-based predictions and conservation of circuit structure across sex and individuals.

Finally we extend the analysis strategy developed for the landing and escape pathways to all 1314 fly DNs and their upstream sensory inputs and downstream motor networks in the VNC. This reveals that the brain controls distinct VNC subnetworks via parallel ensembles of DNs. Several of these ensembles are organized around DNs tightly linked to specific behaviors, such as walking, grooming, takeoff, and landing. This organization suggests that descending control is achieved in a behavior-dependent continuum from command-like control to population coding.

## Results

### An ensemble of DNs controls landing via a core premotor circuit

To investigate how the sensorimotor pathways involved in landing and takeoff behaviors employ serial or parallel organization, we first set out to identify neurons that actuate landing (Fig. 1A). DNp07 and DNp10 are the only neuron types identified in landing control to date (Ache et al., 2019a). Both are DNs, which connect the brain and nerve cord, and feature similar axonal arborizations in the VNC (projections along the leg neuropil oblique track). When activated, each DN drives a classic “landing response” phenotype, which consists of a strong front leg extension with simultaneous extension and lowering of the middle and hind legs (Balebail et al., 2019; Borst, 1986; Goodman, 1960; Tammero & Dickinson, 2002). We reasoned that other DNs that have similar axon morphology might drive the same landing behavior via synapses onto the same motor or premotor neurons in the VNC. To pinpoint additional neurons involved in the fly’s landing response, we first used the connectomes to identify DNs whose output connectivity is similar to DNp07 and DNp10. We used data from two different connectomes: the complete male adult central nervous system (maleCNS, (Berg et al., 2025)) and the male ventral nerve cord (MANC, (Marin et al., 2024; Stürner et al., 2024; Takemura et al., 2024). In each connectome, individual neurons are hierarchically categorized into “groups” (neurons indistinguishable in connectivity and morphology, e.g. bilateral pairs) and then “types” (sets of similar groups which match across connectomes and the literature, see Methods, (Marin et al., 2024)). We treated each DN’s typewise synapse count onto all other neurons as a vector, and computed the cosine similarity of this vector between each of the fly’s 482 DN types and DNp07 or DNp10, respectively (Fig. 1B, Fig. S1-1A).

**Figure 1:**
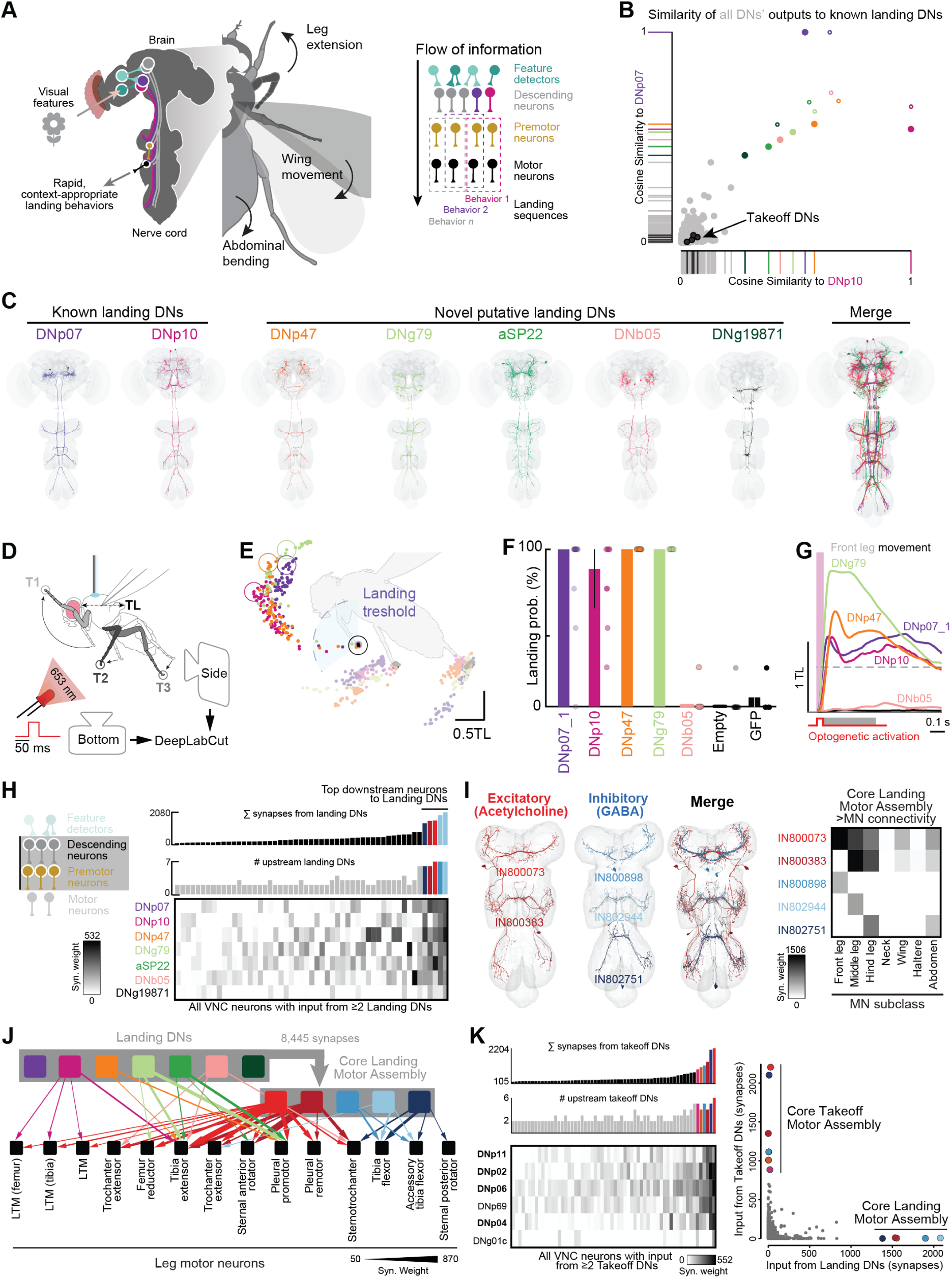
Identification of functional DN ensembles and core motor circuits in the connectome. **A**, Flow of behaviorally-relevant information through the CNS leading to whole-body movements during landing. **B**, Cosine similarity of postsynaptic connectivity of all DNs to DNp07 and DNp10. Filled circles, maleCNS. Open circles, MANC. **C**, Members of the putative Landing DN Ensemble (maleCNS). **D**, Schematic of experimental setup. **E**, Mean leg trajectories of single flies after optogenetic activation of DNp07, DNp10, DNp47, and DNg79 (color code as in (C)). Black, empty control. Dashed line, landing threshold based on 50% of the mean maximum Euclidean distance to rest from all flies in which DNp07 was activated. Center of open circles, maximum Euclidean distance to rest in that particular fly. **F**, Percent of flies that performed landing responses (based on front leg extensions greater than dashed line from (E)). Landing responses for an additional DNp07 driver line are shown in Fig. S1-1C. **G**, Overlay of T1 movement (Euclidean distance) with landing threshold (dashed line) and time window for quantification (gray patch) during optogenetic activation (red). **H**, Left, diagram depicting synaptic connectivity from landing DNs to MNs in the VNC. Bottom right, heatmap depicting connectivity from DNs onto each neuron that receives at least 50 synapses (groupwise) from two landing DNs, sorted by total synaptic input from all landing DNs. Middle right, total number of landing DNs providing at least 50 synapses onto a downstream neuron. Top right, sum of all synapses from landing DNs onto neurons in heatmap columns. **I**, Left, morphology of the Core Landing Motor Assembly, the five neurons that receive the most input from the landing DNs. Right, direct synaptic connectivity between the Core Landing Motor Assembly and MNs, grouped according to the body part innervated by each MN. **J**, Diagram depicting connectivity between landing DNs, Core Landing Motor Assembly, and leg MNs. **K**, Left, heatmap and bar plots depicting connectivity from takeoff DNs onto downstream neurons (Fig. S1-2H, as in (H)). Right, scatter plot of synaptic outputs from takeoff DNs (y-axis) and landing DNs (x-axis) onto downstream neurons identified in (H) and (I).

As expected, DNp07 and DNp10 have substantially similar output connectivity according to this cosine similarity metric. Our approach identified five additional DN types (DNp47, DNg79, aSP22, DNb05, and DNg19871) that share a large fraction of their output with DNp07 and DNp10. Of these, DNp47, aSP22, and DNb05 are all “unique” DN types consisting of one left-right pair, whereas DNg79 includes two left-right pairs, and DNg19871, a group within the DNg06 type (see Fig. S1-2), comprises three DNs on the left and two on the right. In addition to their similar output connectivity, five of the seven DN types exhibited substantially similar morphologies in their VNC branching patterns (Fig. 1C, Fig. S1-1B). DNg19871 is the DN type whose output connectivity is least similar to DNp07 and DNp10 (Fig. 1B), and whose morphology in the brain and VNC is the most different from the other DNs we identified (Fig. 1C, S1-2A). All seven DN types are predicted to be cholinergic, indicating they all have excitatory effects on downstream neurons. We refer to these seven DNs as a putative “Landing DN Ensemble”, which refers to a group of neurons that share similar output connectivity and promote the same behavior (landing). In contrast, previously identified DN types that promote takeoff (DNp02, DNp04, DNp06, DNp11, and the Giant Fiber/DNp01 (Dombrovski et al., 2023; von Reyn et al., 2017)), have low output similarity to DNp07 and DNp10 (Fig. 1B, black points). This suggests descending ensembles controlling takeoff and landing are parallel and non-overlapping.

We next investigated whether the novel putative landing DNs we identified indeed drive landing behavior (Fig. 1D-G). One putative landing DN type, aSP22, is a sexually dimorphic neuron that was shown to drive sequential courtship actions when optogenetically activated in non-flying flies (McKellar et al., 2019). Consistent with a role in landing, it was also shown to induce landing-like leg extensions when activated during flight (McKellar et al., 2019). To achieve specific activation of the other DNs, we used available split-GAL4 driver lines (Namiki et al., 2018; Zung et al., 2025) to drive restricted expression of the red-shifted channelrhodopsin CsChrimson. Genetic driver lines were available for all landing DN types (Supplemental Table 1) except for DNg19871 (Fig. S1-2). Using these tools, we activated each landing DN type optogenetically in pin-tethered flies, video-recorded the flies’ behavior (Fig. 1D), and used DeepLabCut (Mathis et al., 2018) to track movements of the front-(T1), middle-(T2), and hind-leg tarsal tips (T3) (Fig. 1D, Video S1-8). Although landing in both unrestrained and optogenetically-activated animals commonly involves extension of all six legs, the strong extension of both front legs is the most reliable component of the response and has typically been used to define occurrences of the landing behavior. We thus initially classified a movement as landing if front leg extension exceeded 50% of the amplitude elicited by DNp07 activation within the 50 ms activation window (Fig. 1E).

Based on this binary definition, all but one DN in the putative landing ensemble reliably elicited landing responses, whereas control flies exhibited none (Fig. 1F-G, S1-1C). DNb05 was the one putative landing DN that did not elicit landing responses (Fig. 1F-G, Video S6). We confirmed this result (data not shown) using multiple genetic driver lines (see Supplemental Table 1) targeting DNb05 (Zung et al., 2025). The lack of an activation phenotype does not rule DNb05 out as a functional member of the Landing DN Ensemble, as it could contribute to landing in concert with the rest of the DN ensemble even if it does not drive landing in isolation. A similar form of population control has been suggested for takeoff DN types, some of which only elicit behavioral effects when co-activated with other DNs (Dombrovski et al., 2023).

To understand how the Landing DN Ensemble controls motor output, we identified the VNC interneurons onto which the DNs converge by examining their postsynaptic partners in the maleCNS and MANC connectomes. Of the 273 groups that are strongly connected (≥50 groupwise synapses) to any of the landing DNs, only five groups receive more than 1000 combined landing DN input synapses (Fig. 1H). These five groups are dedicated downstream partners of the Landing DN Ensemble. They each receive input from at least six of the seven landing DNs which, together, provide between 21% and 34% of the total synaptic input to each of these downstream neurons (Fig. S1-2D). Mirroring this, between 5% and 15% of each landing DN’s total output is dedicated to these five interneuron groups (Fig. S1-2E). We therefore defined these interneurons as the “Core Landing Motor Assembly”, because they represent the VNC circuit that executes the motor program driven by the Landing DN Ensemble.

The Core Landing Motor Assembly can be divided into two serially homologous sets of VNC interneurons (Fig. 1I, Fig. S1-1D-E, Fig. S1-2F). The first consists of two excitatory cholinergic neuron groups (IN800073 and IN800383) that ramify bilaterally in front and middle leg neuropils, respectively, and additionally output to all ipsilateral leg neuropils, as well as the wing and abdominal neuropils. The second set consists of three inhibitory GABAergic interneuron groups (IN800898, IN802944, and IN802751), each of which project bilaterally within a different thoracic segment (Fig. 1I). Hence, unlike the DN ensemble, the VNC core motor assembly consists of neurons controlling behavioral output via different mechanisms, such as inhibition and excitation, leading to a synergistic motor pattern. The excitatory and inhibitory interneuron sets of the Core Landing Motor Assembly directly synapse onto MNs, but differ in their targets. The excitatory set primarily targets tibia extensor and long tendon muscle MNs in each leg, as well as a number of MNs that are dedicated to moving the coxa anteriorly (i.e., the pleural promotor and sternal anterior rotators, Fig. 1J, Fig. S1-1F). A subset of these MNs also receive direct input from the Landing DN Ensemble, reinforcing this excitation. In contrast, the inhibitory set outputs heavily onto tibia flexor and accessory tibia flexor MNs for each leg pair (Fig. 1J). This connectivity and the fact that the Landing DN Ensemble bilaterally excites both sets suggests that the net output of the Core Landing Motor Assembly would drive tibia extension and rotate the coxa forward while suppressing tibia flexion in all six legs (Fig. 1J, Fig. S1-1G). Thus, we identified an ensemble of DNs that converges onto a dedicated core premotor assembly in the VNC. This network is set up to drive the movements characteristic of landing — smooth extension and anterior rotation of all legs.

Several premotor neurons downstream of takeoff promoting DNs have previously been identified (Cheong et al., 2025). To enable comparison of the VNC circuits controlling takeoff and landing, we performed the same DN similarity analysis with DNp02 and DNp11, two DNs involved in the activation of “long-mode” takeoff (distinct from “short-mode” takeoffs driven by the giant fiber, DNp01) in response to a visual looming stimulus (Dombrovski et al., 2023) (Fig. S1-2G). This analysis revealed a “Takeoff DN Ensemble” comprised of four DN types previously shown to be involved in control of takeoff (DNp02, DNp04, DNp06, DNp11) (Dombrovski et al., 2023) and two DN types not previously associated with takeoff (DNp69, DNg01c). Employing similar criteria as those used to identify landing premotor neurons, we identified six neurons that receive a combined input of 800 synapses from the entire Takeoff DN Ensemble and at least 50 synapses from at least four takeoff DN types (the “Core Takeoff Motor Assembly”, Fig. 1K, Fig. S1-2H). Compared to the Core Landing Motor Assembly, the network downstream of the takeoff DNs was somewhat more diffuse: the Takeoff DN Ensemble members each dedicate between 9 and 25% of their output to the Core Takeoff Motor Assembly (Fig. S1-2L).

Strikingly, there is no overlap in membership or in input connectivity between the landing and takeoff core assemblies, and only a handful of neurons receive more than 100 total synapses from both DN ensembles (Fig. 1J-K, Fig. S1-2H). This suggests that landing and takeoff are controlled by parallel DN ensembles targeting separate VNC networks. The Core Takeoff and Landing Motor Assemblies drive behavior via distinct premotor motifs: whereas the Core Motor Assembly for landing consists of neurons intrinsic to the VNC, that for takeoff includes ascending and descending neurons that span the neck connective (Fig. S1-2I) but also drive leg and wing MNs through both direct and feedforward connectivity (Fig. S1-2J, (Cheong et al., 2025)). The different makeup of landing and takeoff premotor networks suggests different requirements for the motor control of these two behaviors. In particular the integration of ascending neurons into the takeoff premotor architecture suggests that rapid feedback about imminent takeoff is important to communicate directly to the brain, potentially to coordinate other behavior modules. These feedback pathways might signal the initiation of flight (Cheong et al., 2024), which requires a variety of physiological changes, including modulation of sensory processing (Longden & Krapp, 2010; Maimon et al., 2010; Suver et al., 2012), neuroendocrine pathways, and metabolism (Liessem et al., 2023).

### DNs within an ensemble promote behavioral variants

Outside of their shared outputs onto the Core Landing Motor Assembly, the landing DN types differ substantially in their output connectivity (Fig. 1H, J). This divergence could enable finer control of the movements produced by the ensemble through differential recruitment of individual DNs. In line with this hypothesis, optogenetic activation of each landing DN type drove unique leg kinematics, resulting in varying extension amplitudes and directions for different legs (Fig. 2A-B, F, Fig. S2-1A-C). For example, activation of DNg79 produced greater front and middle leg extension than any other landing DN, and DNp10 elicited less forward rotation of the middle and hind legs than DNp07 (Ache et al., 2019a) and the other landing DNs (Fig. 2A-B, Video S5).

**Figure 2:**
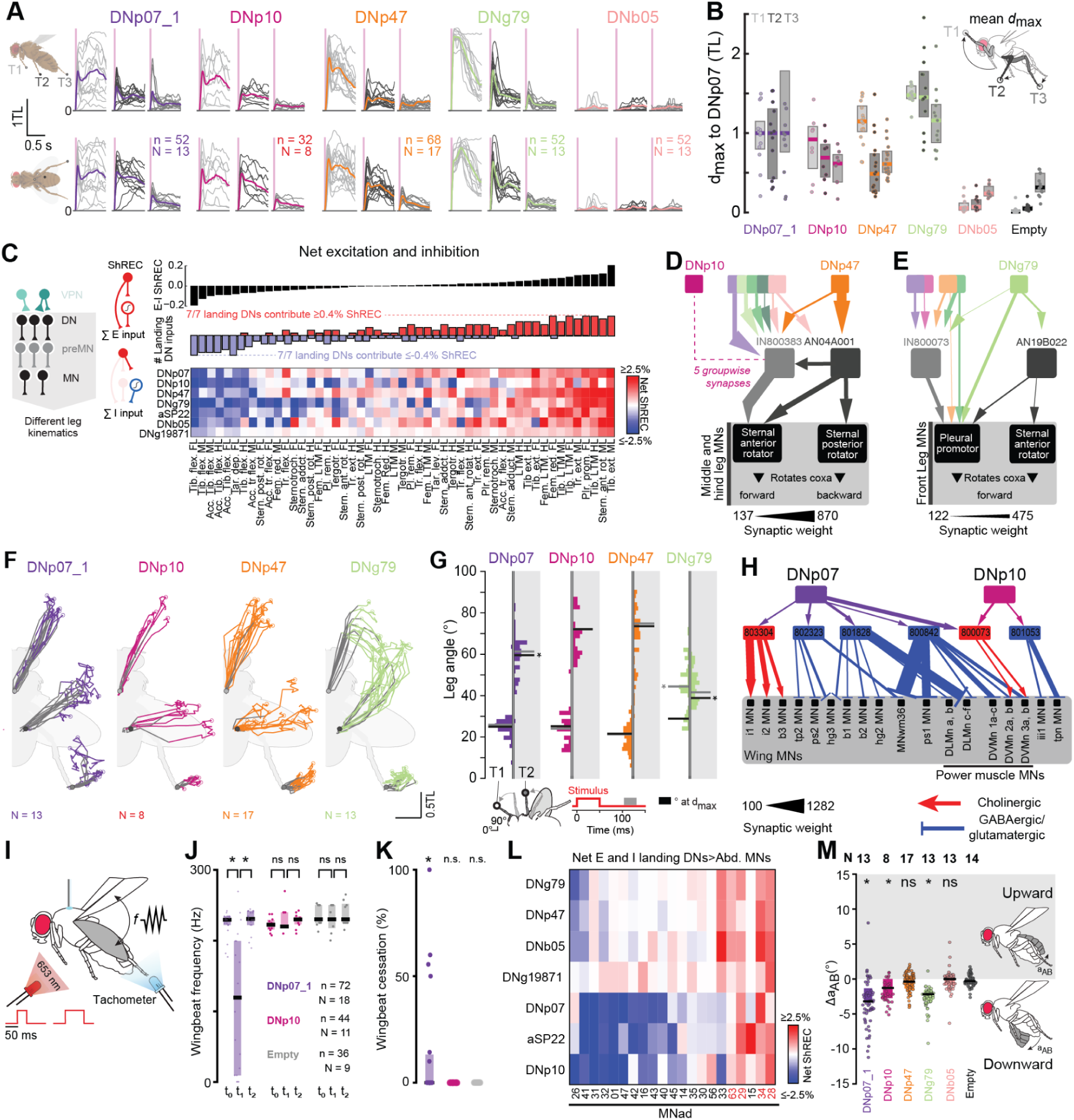
Connectivity of DNs in the nerve cord predicts commonalities and differences in leg kinematics. **A**, Mean movement amplitude (Euclidean distance from resting position) of front, middle, and hind leg during DN activation of all flies (grey). Colored line, grand mean of all flies. **B**, Quantification of the maximum Euclidean distance of T1-T3 normalized to DNp07. **C**, Left, diagram depicting synaptic connectivity from landing DNs to MNs in the VNC. Right, net Short-Range Effective Connectivity (ShREC) from landing DNs onto MNs driving different leg muscles, grouped by neuron type and separated by leg segment. Bottom, net ShREC (excitatory ShREC-inhibitory ShREC) from each landing DN type onto each leg MN type. FL, front leg; ML, middle leg; HL, hind leg; tib., tibia; flex., flexor; tar., tarsal; dep., depressor; accs., accessory; stern., sternal; rot., rotator; lev., levator; tr., trochanter; red., reductor; pl., pleural, LTM, long tendon muscle. Middle, total number of landing DNs that contribute net ShREC of at least ±0.4% to each leg MN type. Blue bars, number of DNs contributing −0.4% net ShREC. Red bars, number of DNs contributing 0.4% net ShREC. Top, net ShREC from all landing DNs to each leg MN. All columns are sorted by net ShREC. Heatmap displays values between −2.5% and 2.5%, actual minimum −7.6% (DNb05:tr.ext. FL) and actual maximum 4.3% (DNb05:tr.ext. ML). **D-E**, Typewise connectivity diagram of landing DNs and additional premotor and motor neurons outside the Core Landing Motor Assembly. IN800383 drives MNs of the middle and hind leg sternal anterior rotator. AN04A001 drives MNs of the middle and hind leg sternal anterior and posterior rotators. AN190B022 drives MNs of the pleural promotor and sternal anterior rotator muscles of the front leg. Minimum threshold for displaying connection: 135 (D) or 120 (E) synapses. **F**, Optogenetic activation of DNs drives leg extensions with kinematic differences. Ventral view of the fly with superimposed mean trajectories of the tarsal tips (T1-T3) during (grey) and after optogenetic activation (color coded) of the respective neurons. **G**, Probability distribution of T1 and T2 leg angles (bandwidth 2°) from rest and along the anterior-posterior axis/midline within the last third of our main analysis window (80 ms). Gray line, median leg angle from each fly within that analysis window. Black line, median leg angle at maximum Euclidean distance from rest from each fly in the full analysis window. Grey and black asterisks, significantly different (one-way ANOVA with Tukey–Kramer post hoc correction for multiple pairwise comparisons) from DNp10 in the respective time windows. **H**, Connectivity from DNp07 and DNp10 onto MNs controlling wing musculature. “Power Muscle MNs” indicate MNs that innervate power muscles that deform the thorax to beat the wings upwards or downwards. Arrowhead, cholinergic connection. T-bar, GABAergic/glutamatergic connection. Minimum threshold for displaying a connection is 100 synapses. **I**, Schematic of experimental setup for optogenetic activation of DNs to measure effects on wing movement. **J**, Optogenetic activation of DNp07 reduces wingbeat frequency during flight. Quantification of wingbeat frequencies for each fly in the time windows (t_0_-t_2_) depicted in Fig. S2-2C. Only the first four trials in which flies did not cease flight after DN activation were considered. All p-values calculated via two-sided Wilcoxon rank-sum tests. **K**, Number of optogenetic activation trials (out of 10) in which the fly ceased to fly. DNp07, N = 19; DNp10, N = 11; Empty, N = 9. All p-values calculated via Wilcoxon signed rank tests. **L**, Net ShREC from all landing DNs (rows) to strongly connected abdominal MNs (columns). Columns are sorted to only include those MNs with total excitatory or inhibitory ShREC >1% and sorted from lowest to highest net ShREC. Heatmap displays values between −2.5% and 2.5%, actual minimum −7.2% (DNp10:MNad26) and actual maximum 7.4% (aSP22:MNad15). Red text highlights abdominal MNs that receive strong convergent excitation from landing DNs (≥0.4% excitatory ShREC from each DN and net ShREC summed across all landing DNs>0). **M**, Quantification of abdominal deflection within 80 ms after stimulus onset quantified as deviation from the abdominal resting position (mean during 100 ms before activation).

To analyze how these kinematic differences might arise from differences in connectivity, we developed a metric, <u>Sh</u>ort-<u>R</u>ange <u>E</u>ffective <u>C</u>onnectivity (ShREC, see Methods), to estimate the synaptic influence of all landing DNs onto downstream MNs. Because it is difficult to determine the net effects of multiple sign (excitation and inhibition) changes across more than two synaptic hops, we confined our analysis to quantifying direct excitation, disynaptic excitation, and disynaptic inhibition for each landing DN (Fig. 2C, Fig. S2-1D). To this end, each VNC neuron group was classified as excitatory (cholinergic neurons) or inhibitory (glutamatergic or GABAergic neurons), according to the neurotransmitter predictions available in maleCNS (Eckstein et al., 2024). We weighted the outputs of neurons directly downstream of the landing DNs, using a logistic function to scale their synaptic influence according to the excitatory input they receive from each landing DN type (see Methods). We then summed all sources of excitation and inhibition onto every leg MN (Fig. 2C). Consistent with their strong connectivity onto the Core Landing Motor Assembly, each landing DN provides common patterns of excitation and inhibition to synergistic muscles that drive the characteristic landing response — extension of all six legs (Fig. 2C, Fig. S1-1G). But we also discovered differences in effective connectivity and underlying wiring patterns that provide compelling explanations for the differences in leg movements elicited by each DN (Fig. 2A-C). For instance, DNp10 only provides five synapses onto the excitatory Core Landing Motor Assembly neuron IN800383 (Fig. 2D), which rotates the middle and hind legs forward via the sternal anterior rotator MNs. This is a much lower synapse count than all other landing DN types exhibit. Similarly, DNp47 provides substantial synaptic input onto IN800383, but also outputs extensively onto an excitatory premotor neuron type outside the core set, AN04A001, which putatively rotates the same segment backwards through simultaneous excitation of the sternal posterior rotator MNs. Hence, unlike the other landing DN types, DNp10 and DNp47 are not set up to drive strong forward rotation of the middle and hind legs. In concordance with this, DNp10 and DNp47 activation drives significantly less anteriorly directed middle leg extension than other landing DNs (Fig. 2F-G).

Another example of connectivity that supports differential control of leg movement is that of DNg79. DNg79 provides strong synaptic input to AN19B022, a cholinergic ascending neuron, which in turn synapses onto MNs driving pleural promotor muscles that rotate the front leg forward at the coxa joint (Fig. 2E, Fig. S2-1F). In parallel, DNg79 is connected to GABAergic interneurons such as IN13A051, IN13A011, IN16B014, IN19A024, and IN19A016 (Fig. S2-1F), which strongly inhibit the MNs targeting the front leg sternotrochanter, the trochanter extensor, the sternal posterior rotator, and the pleural remotor muscles (the latter two of which rotate the coxa backwards). This connectivity suggests that stimulation of DNg79 should evoke unique movements of the first two leg segments, the coxa and trochanter, which would affect the trajectories of all distal segments.

Analyzing leg movements following the optogenetic activation of different landing DN types confirmed these predictions. DNp10 and DNp47 consistently drove more laterally-directed middle and hind leg movements compared to all other DNs (Fig. 2F-G, Fig. S2-1G-H, Video S1-6). During DNg79 activation, the front leg coxae rotated forwards and underneath the head, followed by a dorsally directed, lateral extension of the distal segments (Fig. S2-1I, Video S5). In parallel, the middle legs reached a very anterior position (Fig. 2F-G, Fig. S2-1I). Thus, the tips of the front and middle legs follow a trajectory that is distinct from all other landing DN types - matching the unique downstream connectivity of DNg79. We suspect that the movement sequence driven by DNg79 could support inverted landing on surfaces above the fly, during which the front legs are rotated to reach above the head (Balebail et al., 2019; Hyzer, 1962; Liu et al., 2019).

So far we have considered the role of landing DNs and their downstream targets in the context of leg motor control during the prominent and rapid leg extensions flies display during landing. However, observations from both tethered and free-flight landing approaches indicate that *Drosophila* coordinate leg movements with other body parts, including the wings and abdomen (Dickinson & Tu, 1997; Götz et al., 1979; Zanker, 1988), and eventually must cease flight. Given their complex outputs in the VNC, landing DNs could contribute to all or several of these movements, and thus control the entire landing sequence by actuating the legs, wings, and abdomen. To investigate potential landing DN control of these other body parts, we calculated excitatory and inhibitory ShREC from the landing DNs onto all MNs controlling the wings (Fig. 2H, Fig. S2-2A-B) and abdomen (Fig. 2L, Fig. S2-2H). In the wing motor system, large thoracic power muscles (DLM, dorsolateral muscles; DVM, dorsoventral muscles) drive wing beating, while smaller steering and indirect control muscles adjust wing orientation, stroke amplitude, and thoracic tension (Dickinson & Tu, 1997; Lindsay et al., 2017). Some members of the landing ensemble have strong mono- and di-synaptic connectivity onto MNs controlling wing muscles of all three classes (power, steering, and indirect control). DNp07, for instance, recruits GABAergic interneurons that synapse onto MNs targeting the steering and power muscles and are thus predicted to suppress flight (Fig. 2H, Fig. S2-2A-B). It also synapses onto a cholinergic interneuron that provides strong output onto MNs controlling steering muscles i1 and i2 (IN803304, Fig. 2H), which likely decrease wing stroke amplitude (Lindsay et al., 2017). Other members of the Landing DN Ensemble provide highly selective output onto a sparse subset of wing MNs. For instance, DNp10 exerts limited influence on the activity of most wing MNs, but does provide feedforward inhibition onto the iii1 and tpn indirect control MNs via IN801053 (Fig. 2H, Fig. S2-2A-B). One excitatory member of the Core Landing Assembly (IN800073), which receives input from the entire Landing DN Ensemble, excites DVMn, the MN that drives the power muscle raising the wings. Taken together, these connectivity patterns suggest that DNp07 is wired to actuate wing movements related to active descent and potentially the cessation of flight, whereas DNp10 might induce more selective wing movements via its excitation of DVMn and sparse connections to steering MNs.

To test these predictions, we optogenetically activated DNp07 and DNp10 while measuring wing beat frequencies during flight (Fig. 2I-J). Activation of DNp07, but not DNp10, markedly reduced the wingbeat frequency during flight (Fig. 2J, Fig. S2-2C-D, Video S1-3), and led to the cessation of flight in 6/19 flies (Fig. 2K). This effect was transient and graded, such that high stimulus intensities drove a stronger decrease in wingbeat frequency (Fig. S2-2E-F). Interestingly, these effects were reversed in non-flying flies, in which activation of DNp10, but not DNp07, caused flies to raise their wings (Fig. S2-2G, Video S9), suggesting that the effects of landing DNs onto the wing motor system depend on the fly’s behavioral state.

We observed strong net excitation (Excitatory ShREC ≥0.4% and positive net ShREC) from each of the landing DNs onto a select number of abdominal MNs (MNad28, MNad29, MNad63, and MNad34, Fig. 2L, red text). We hypothesized that this convergent excitatory input underlies abdominal movements common to many landing sequences. To determine what these movements might be, we measured abdominal bending during optogenetic activation of the landing DNs (Fig. 2M, Fig. S2-2I-J). Activation of DNp07, and to a lesser degree DNp10 and DNg79, produced significant downward deflection of the abdomen. In contrast, activation of DNp47 and control flies did not result in downward deflection of the abdomen (Fig. 2M, Fig. S2-2J). Downward deflection could contribute to a pitch-up maneuver, which flies employ to decelerate when landing on vertical substrates (Balebail et al., 2019; David, 1978; Wagner & Land, 1997). In particular, the abdominal bending induced by DNg79 might benefit overhead landings by changing the body pitch angle, matching the leg movements driven by this DN.

In summary, DNp07, DNp10, DNp47, and DNg79 drive extensions of all six legs as part of the landing response, via their shared connections to the Core Landing Motor Assembly. Each landing DN type additionally targets a unique set of VNC neurons that enable variations of leg movements during landing as well as different wing and abdomen movements. Hence the Landing DN Ensemble differentially controls movements of all body parts associated with landing so that the recruitment of subsets of landing DNs could fine-tune body movements for different situations. Analogously, individual members of the Takeoff DN Ensemble enable takeoff in different directions by shifting the body’s center of mass before leg extension (Dombrovski et al., 2023). This suggests a similar organization of both ensembles into DN subsets that drive context-dependent variations of a core motor pattern.

### DN Ensembles for landing and takeoff are recruited by parallel visual pathways

Thus far we have shown that the landing and takeoff motor programs are coordinated by non-overlapping ensembles of DNs and premotor neurons, but it remains unclear if these distinct assemblies are recruited by common sensory circuits. Previously, we have shown that the exact same frontal looming stimulus can induce flies to either land or take off, depending on whether or not they are in flight, and that both DNp07 and DNp10 are only responsive to visual stimuli during flight (Ache et al., 2019a). Based on these findings, an efficient circuit organization would be for a singular set of looming detecting neurons to activate both landing and takeoff DNs, with the visual input gated out during non-flight periods for landing DNs and gated out during flight for takeoff DNs in order to couple the looming information to the appropriate motor pathway in each behavioral state.

To examine visual recruitment of Landing vs. Takeoff DN Ensembles, we focused on Visual Projection Neurons (VPNs), which convey information from the optic lobes to the central brain (Klapoetke et al., 2022; Mathew et al., 2023; Panser et al., 2016; Wu et al., 2016) (Fig. 3A-D). One class of VPN, the columnar VPNs, receive synaptic input within layers of retinotopically arrayed optic lobe neuropils, the lobula and/or lobula plate (Fig. 3E). This arrangement endows these neurons with receptive fields tuned selectively to particular visual features, such as looming, localized to specific locations within the fly’s visual field. Takeoff DNs were demonstrated to be directly activated by four columnar VPN types: LC4, LPLC2, LPLC1, and LC6 (Ache et al., 2019b; Dombrovski et al., 2023; von Reyn et al., 2017).

**Figure 3:**
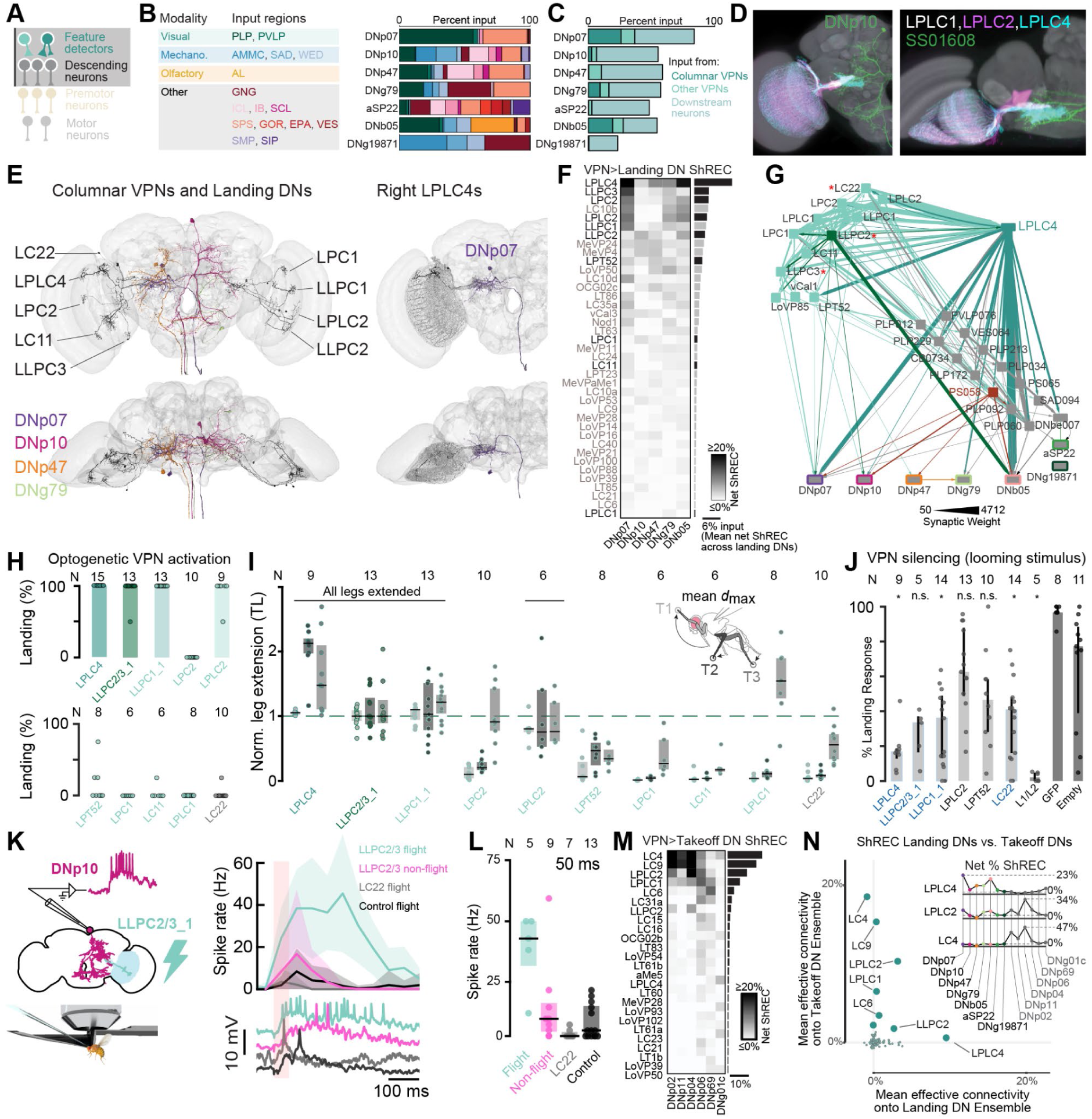
A subset of parallel feature detecting VPNs converge onto the Landing DN Ensemble and are necessary and sufficient for landing. **A**, Diagram depicting synaptic connectivity from sensory feature detectors to landing DNs in the brain. **B**, Proportion of inputs to each landing DN type according to brain region, colored according to region (key on left). Regions that are bilaterally symmetric are summed into a single category. PLP, posterior lateral protocerebrum; PVLP, posterior ventrolateral protocerebrum; AMMC, antennal mechanosensory and motor center; SAD, saddle; WED, wedge; AL, antennal lobe; GNG, gnathal ganglia; ICL, inferior clamp; IB, inferior bridge; SCL, superior clamp; SPS, superior posterior slope; GOR, gorget; EPA, epaulette; VES, vest; SMP, superior medial protocerebrum; SIP, superior intermediate protocerebrum **C**, Proportion of inputs to each landing DN type that stem directly from a columnar Visual Projection Neuron (“columnar VPN,” dark green), directly from another VPN type (“other VPNs,” teal), or from any non-VPN neuron that receives at least 10 synaptic inputs from any type of VPN (“downstream neurons,” pale green). **D**, expression pattern of several VPN driver lines and their overlap with DNp10. The illustrations show two views of an overlay of registered confocal images of split-GAL4 driver lines with the template brain used for registration. **E**, Presynaptic sites of VPNs in (D) exhibit substantial overlap with landing DN dendrites. **F**, Weighted mono- and di-synaptic connectivity from cholinergic VPNs (rows) onto landing DNs (columns). Rows are filtered to include only the top 40 VPNs in terms of mean effective connectivity onto landing DNs. VPNs tested in (H-I) are labeled in black. For unfiltered VPN-DN connectivity, see Figure S3-3A. Right, net ShREC averaged across all landing DNs. Heatmap displays values between 0 and 20%, actual minimum −5.3% (LPLC2:DNp47), actual maximum 23.4% (LPLC4:DNp07) **G**, Selected VPNs that connect directly and indirectly to the Landing DN Ensemble mainly via LPLC4. Red asterisks, VPNs that are optogenetically stimulated in (K-L). **H**, Percent of flies that perform landing responses when indicated VPN is optogenetically stimulated (based on front leg extensions, see also Fig. S3-2). **I**, Mean maximum Euclidean distance of the front, middle, and hindleg, normalized to LLPC2/3_1 responses. Dots, single fly averages. Detailed leg kinematics are depicted in Fig. S3-2F-G. Leg movements were captured in side-view at 1000 Hz. **J**, Landing probability of VPN silenced flies using Kir2.1 in response to a frontal looming stimulus. VPNs highlighted in blue, significant reduction of landing in at least 2 out of 3 visual stimulation protocols (loom, approaching bar, moving bar, Fig. S3-2I-J, all p-values from Wilcoxon rank sum tests). For p-values, see Supplemental Table 2. **K**, Left, Experimental design for *in vivo* functional connectivity experiments. Right, Spike rate of DNp10 (bin size 50 ms) during 50 ms long optogenetic stimulation of LLPC2/3_1, LC22 and empty control flies during flight. Bottom, example traces. Top, average spike rate across flies (N, shown in (L)). Magenta trace shows response to LLPC2/3 stimulation during non-flight. **L**, Quantification of DNp10 spike rate within 100 ms after LED offset. **M**, Weighted effective connectivity from cholinergic VPNs onto takeoff DNs, as in (F). Right, net ShREC averaged across takeoff DNs as in (F). Heatmap displays values between 0 and 20%, actual minimum −2.3% (LT82b:DNp11), actual maximum 47.1% (LC4:DNp04). **N**, Effective connectivity of each VPN onto landing DNs (averaged across DNs) plotted against effective connectivity of the same VPN onto takeoff DNs (averaged across DNs). Inset, effective connectivity of three example VPNs onto each member of the Landing and Takeoff DN Ensemble.

To determine whether landing DNs are recruited by these same VPNs, we analyzed landing DN upstream connectivity (Fig. 3A). Each Landing DN except DNg19871 has dendritic arborizations in visual neuropils, such as the posterior lateral protocerebrum and posterior ventrolateral protocerebrum (Fig. 3B), and receives mono- and di-synaptic inputs from columnar VPN types (Fig 3C-D). We used ShREC to estimate the influence of each of the 248 cholinergic VPN types onto the five landing DNs (DNp07, DNp10, DNp47, DNg79, and DNb05) that receive the strongest input from VPNs (Fig. 3F, Fig. S3-3A). A subset of VPNs, especially LPLC4, have strong effective connectivity onto each of these landing DNs. LPLC4 synapses directly onto DNp07, DNg79, and DNb05 while also exciting PS058, one of the highest-weighted cholinergic inputs of both DNp47 and DNp10 (Fig. 3G). Many of the other VPNs that connect indirectly to the Landing DN Ensemble, such as LPLC2, LLPC1, and LC22, do so primarily via their output to LPLC4 (Fig. 3G). Hence, LPLC4 serves as a critical hub that links the Landing DN Ensemble to input from a variety of anatomically distinct visual feature detectors spanning the entire visual field. While every member of the Landing DN Ensemble receives strong input from LPLC4, the landing DN ensemble receives differential input from the other VPN types. For instance, LLPC2 displays stronger synaptic connectivity to DNp10 compared to DNp07 and DNg79, which in turn receive more input from LLPC3. (Fig. 3F-G).

While it is expected that VPNs with strong effective connectivity to the Landing DN Ensemble, like LPLC4, are critical for landing, there is no clear way to predict whether VPNs with weaker connectivity to the DN layer are similarly indispensable. In principle, each neuron that connects to the DN layer could affect behavior, with the number of synapses determining the magnitude of the contribution. VPN types such as LC22, for example, are linked to the Landing DN Ensemble by lower-weight synaptic pathways, and may still contribute to the motor drive as part of a larger population (Fig. 3G). To functionally assess the degree to which the VPNs implicated by our connectome analysis are involved in landing behavior, we tested whether optogenetic activation of individual VPNs drove landing or other movements in flight, and whether silencing them impaired visually-induced landing responses. We selected eleven VPN types for which cell-type specific split-GAL4 lines were available (Fig. S3-1), including VPNs with strong direct and disynaptic connections to the landing DNs (LPLC4, LLPC1, LLPC2, LLPC3, LPLC2, LPC2, LPT52), and those with weaker, indirect connections (LC22, LC11, LPC1, LPLC1).

Individual VPN types were activated optogenetically in tethered, flying flies while we monitored the flies’ behavioral responses (Fig. 3H-I, Fig. S3-2A-G). Activation of VPN types with very strong effective connectivity to the Landing DN Ensemble (LPLC4, LLPC1, LPLC2, and one split-GAL4 line that expressed in both LLPC2 and LLPC3, referred to as LLPC2/3), drove robust landing responses, as defined by the bilateral front leg extensions typical for landing that occurred simultaneously with middle and hind-leg movements (Video S11-21). This is consistent with the idea that landing involves movements of all legs. Interestingly, different VPN types drove different leg movement amplitudes (Fig. 3I). For instance, LPLC4 activation consistently elicited stronger middle leg extensions than other VPN types (Fig. 3I, Fig. S3-2F-G). Other VPNs, including LPLC1 and LPC2, failed to elicit the strong front leg extensions typical for landing, but reliably drove more restricted body movements, such as shifting the hind legs (Fig. 3I, Fig. S3-2F-G, Video S13 and S15) or bending the abdomen (Fig. S3-2H). These VPNs might contribute to landing responses, but do not drive the entire motor program. For those VPNs that did drive front leg extension, stronger activation generally produced leg extensions with higher amplitudes, higher velocities, and longer duration (Fig. S3-2D). This suggests that VPN activity does not simply trigger landing, but finely controls landing movements, presumably via increased drive to individual DNs and potentially the recruitment of additional DNs at higher activity levels. In contrast, optogenetic activation of L1 and L2 neurons, which are core inputs to the motion vision pathway and thus upstream of most VPNs (Rister et al., 2007), did not drive landing-like leg extensions (Fig. 3I, Fig. S3-2B, F, Videos S21). Taken together, our results show that multiple VPN types can drive landing responses and differentially recruit DNs to drive different combinations of leg and body movements.

We next silenced VPN types via cell type-specific overexpression of the inwardly rectifying potassium channel Kir2.1. Flies were exposed to a series of visual stimuli that ranged in their effectiveness at eliciting landing responses (approaching bar > looming > moving bar), and silencing effects were assessed by scoring the likelihood of each stimulus to elicit the front leg extensions typical for landing. Silencing LPLC4, LLPC2/3, or LLPC1 diminished the likelihood of landing responses to visual stimulation for at least two of the three stimuli compared to controls (Fig. 3J, Fig. S3-2I-J). As expected, these effects were weaker than the impact of silencing L1 and L2 neurons, which renders flies motion-blind and abolished landing (Rister et al., 2007). Silencing other VPNs, including LPLC2 and LPT52, did not significantly affect landing probability as defined by front leg extension (Fig. 3J, Fig. S3-2I-J). However, these neurons might still contribute to other aspects of the landing response, or other flight-related behaviors.

Given the complex nature of sensorimotor pathways in general and the VPN-DN connectivity in particular, it is difficult to predict how activity dynamically spreads through the network in behaving flies. To directly assess this, we performed patch-clamp recordings from the DN layer while activating VPNs optogenetically (Fig. 3K). Since its functional role in the Landing DN Ensemble is well-established, we generated genetic reagents to use DNp10 as a readout to test VPN-DN connectivity. First, we tested LLPC2 VPNs (using the LLPC2/3 line), which we expected to be a core component of the landing circuit given their strong connections to the DN ensemble and their robust activation and silencing phenotypes. Second, we tested LC22, which provides weaker, indirect connections to the Landing DN Ensemble, and does not evoke landing when activated. Thus, we selected one VPN type which is predicted to drive strong activity changes in the DN, and one which is expected to have weaker effects.

As suggested by the connectome analysis, optogenetic activation of LLPC2/3 drove strong depolarizations and increased the spike rate of DNp10, confirming that this is a strong and functional excitatory connection (Fig. 3K-L, Fig. S3-2K). This response was significantly stronger than that of two genetic controls, which revealed only a weak visual response to the light pulse used for optogenetic activation. LC22 activation, on the other hand, resulted in a negligible increase in the DNp10 membrane potential and spike rate (Fig. 3K-L, Fig. S3-2K). Together, these results confirm that our EM-based connectivity predictions are functionally relevant and accurate.

DNp10 and DNp07 exhibited much weaker responses to visual stimuli during non-flight periods in an earlier analysis (Ache et al., 2019a). We leveraged our connectivity analysis and genetic reagents to directly test whether this state-dependence is an inherent feature of the landing circuit or inherited from upstream connections by recording DNp10 responses to optogenetic excitation of LLPC2/3 during flight and non-flight periods. The strong excitatory influence of LLPC2/3 on DNp10 was diminished during non-flight periods and comparable to that of negative controls during flight (Fig. 3K-L). Hence, the functional connectivity of the LLPC2/3 VPNs and DNp10 depends on the fly’s behavioral state. Our findings demonstrate this state-dependent modulation occurs at the VPN-DN interface rather than via an increase in upstream visual neuron responsiveness. Such behavioral state gating ensures that VPNs only recruit landing DNs in the appropriate behavioral context.

To our surprise, the Landing DN Ensemble received inputs almost entirely distinct from known takeoff DN inputs (Fig. 3M-N, Fig. S3-2L). Indeed, only LPLC2 was among the stronger upstream inputs to both the Landing and the Takeoff DN Ensembles (Fig. 3N, S3-3B). To systematically compare VPN inputs to the Landing and Takeoff DN Ensembles, we performed the same ShREC analysis for direct and indirect VPN input to takeoff DNs as we had to landing DNs. The five strongest VPN inputs to the Takeoff DN Ensemble are previously identified looming-responsive VPNs that make direct synapses onto takeoff DNs (LC4, LC9, LPLC2, LPLC1, and LC6). Our analysis also showed that, in addition to LPLC2, LLPC2 had moderately strong input to both Landing and Takeoff DN Ensembles, but this did not apply to any other VPN type (Fig. S3-3B-C). Hence, landing and takeoff are mediated by distinct, largely non-overlapping VPN populations. For one of the two VPNs the pathways share (LLPC2), we have shown that its connectivity onto the landing DN layer (DNp10) is gated by flight state. Extending our analysis beyond VPNs to all upstream inputs, we discovered that landing and takeoff DN types do not make direct connections to the same neurons, and are hence recruited by parallel sensory streams with little overlap (Fig. S3-3D).

We hypothesize that this parallel architecture permits independent modulation of sensorimotor ensembles controlling different behaviors to enable flexible and context-dependent sensorimotor coupling. Moreover, this parallel organization could allow takeoff and landing DNs to encode different parameters of the same looming stimuli to account for the different temporal requirements of takeoff and landing relative to a perceived approach.

### DN visual receptive fields match movement directions

The behavioral variants elicited by different landing DNs could represent specific responses to visual stimuli, for instance motion in different locations of the visual space. This raised the possibility that subsets of DNs can be differentially recruited by the layer of visual feature detectors. Each VPN type is composed of many neurons tiling the visual space, so that each VPN possesses a unique spatial receptive field. For example, LPLC4 comprises a set of about 50 neurons on each side of the brain. The dendrites of individual LPLC4 neurons are arrayed retinotopically throughout the lobula complex in the optic lobe and each receives strong spatially tuned net excitation spanning ∼22° of the fly’s ipsilateral visual field horizontally and vertically (S4-1A). We therefore analyzed if individual neurons within a VPN type preferentially target specific members of the Landing DN Ensemble to tune them towards different locations of visual space.

We first looked at the direct connectivity from LPLC4 neurons onto postsynaptic cells and found that LPLC4 neurons with receptive fields in different parts of the fly’s visual field tended to connect to distinct downstream populations, including landing DNs (Fig. 4A-B). For example, DNp07 tends to receive input from LPLC4s with ventrally-directed receptive fields, while DNg79 receives input from dorsally-directed LPLC4s. PS058, a strong excitatory input to both DNp10 and DNp47, receives its strongest input from medial LPLC4s. We validated our predictions of DN-specific retinotopic gradients with a dynamical method that relied upon neuronal connectivity from a separate connectomic resource (FAFB/FlyWire). Using a connectome-constrained leaky integrate-and-fire model (Shiu et al., 2023) we predicted the activity of each Landing DN upon activation of spatially segregated LPLC4 clusters (Fig. S4-1B). We found that all landing DNs receive strong excitation from at least one of these LPLC4 clusters. However, all DNs receive most excitation from a preferred zone of the visual field. These results validate our observations of effective connectivity in the maleCNS connectome: In the simulation, DNg79 neurons respond most to dorsal clusters, DNp07 is mainly driven by anterior-ventral clusters, and DNp10 and DNp47 mainly respond to more posterior-ventral clusters (Fig. S4-1B).

**Figure 4:**
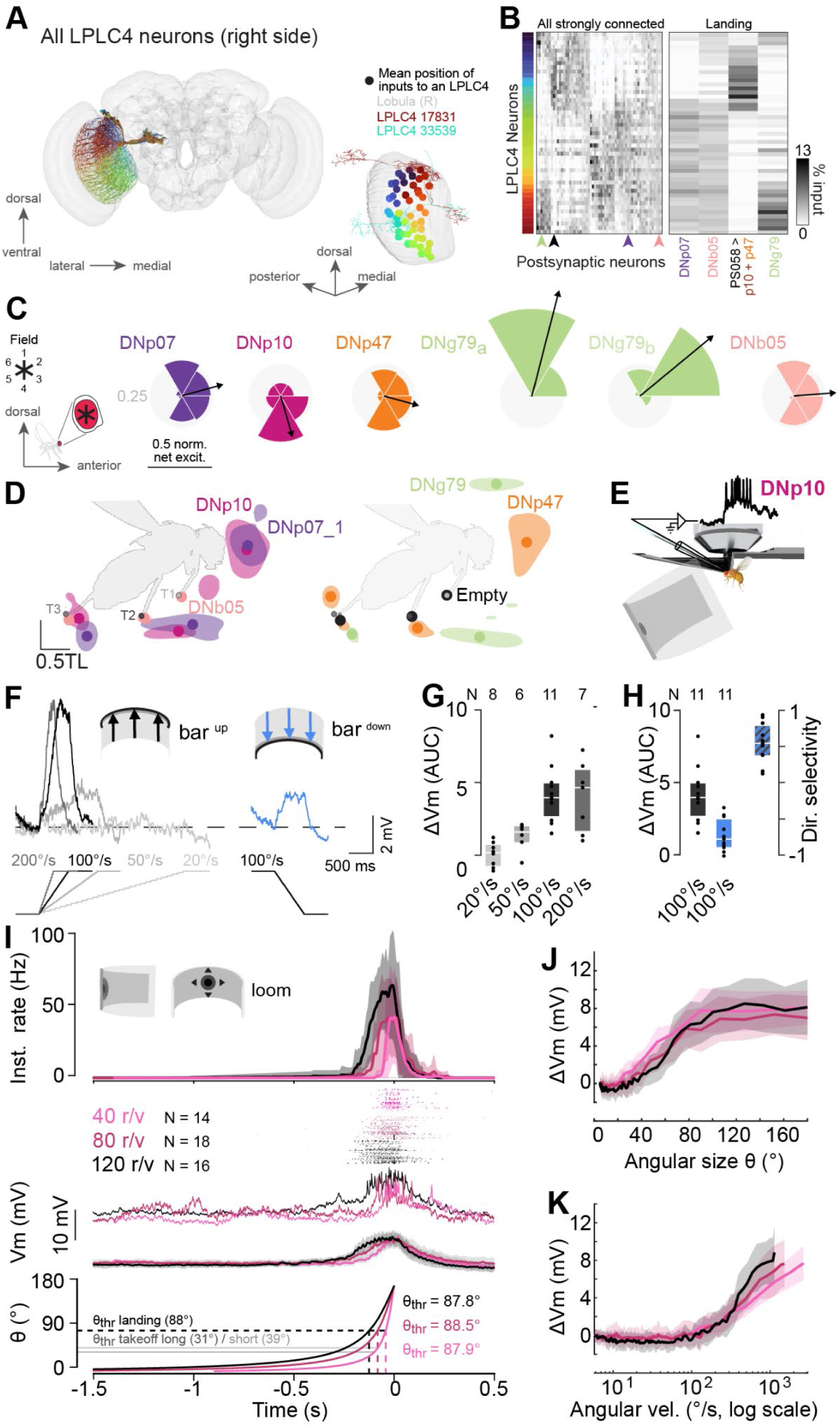
Landing DNs receive input from a subset of parallel feature detecting VPNs. **A**, LPLC4s have retinotopically-tuned dendritic fields in the lobula that collectively span the fly’s visual field (front view). Top, Each LPLC4 colored according to the mean location of its dendritic field. Bottom, Mean locations of LPLC4 dendritic fields in the right lobula alongside two exemplary LPLC4 neurons. **B**, LPLC4 neurons (rows, sorted according to position of their dendritic fields in lobula) output onto downstream neurons (columns, neurons sorted typewise) according to the location of their retinotopic inputs (left bar). Arrowheads indicate columns expanded in right panel. Right, LPLC4s sensitive to distinct parts of the visual field make differential outputs onto landing DNs and PS058, one of the highest-weighted cholinergic inputs to both DNp47 and DNp10. **C**, Normalized net ShREC from visual fields onto each Landing DN. Bin size, 60°. Grey circle, 0.25 normalized net ShREC. Arrow, directional vector. **D**, Probability density distribution (50% cutoff) of the mean Euclidean distance from resting position of all files within 400 ms after optogenetic activation. Solid circle, grand mean of the maximum Euclidean distance. **E**, Schematic of experimental design to record DNp10 activity during visual stimulation. **F**, Visual responses of DNp10 to horizontal upward (grayscale) and downward (magenta) bar stimuli with different speeds **G**, Change in membrane potential during visual bar stimulation (plus another 100 ms after offset of stimulation) as area under the curve and normalized to the stimulus duration. **H**, Responses to upward- and downward-directed bar stimuli at 100°/s (see Methods). **I**, DNp10 response to visual looming stimuli at different *r/v*. From top to bottom, change in instantaneous spike frequency, spike events in each trial per fly, three example recordings of DNp10 during different looms, mean membrane potential with standard deviation (shaded areas) of all flies during different looming speeds, looming stimulus (color coded based on speed). For clarity, only the last 1.5 s of the looming responses are shown. Dashed vertical lines indicate the time at which instantaneous firing rate reached 50% of its maximum, with corresponding θ values denoting the angular size of the looming stimulus at that response time (mean as horizontal black line). Horizontal grey lines, corresponding θ at 40 r/v for short- and long-mode takeoff from (von Reyn et al., 2014). **J**, Mean DNp10 response to looming stimuli with different *r*/*v* values plotted against instantaneous looming angular size. **K**, As in (J), but plotting DNp10 membrane potential against instantaneous angular velocity.

We extended this analysis beyond LPLC4 to include all VPNs, and identified similar synaptic gradients that drive sharp differences in retinotopic tuning between the landing DNs (Fig. 4C). Briefly, using the maleCNS connectome, we computed ShREC from all VPNs one or two synaptic hops upstream of each landing DN, and multiplied these values by the visual tuning of each VPN (computed as the net excitatory input to each VPN within each lobular column). DNg79 received strong net mono- and di-synaptic excitation from VPNs tuned to the dorsal visual field (Fig. 4C). In contrast, DNp10 is tuned towards the ventral and anterior visual field, while the remaining landing DNs receive input from VPNs sensitive to visual features in front of the fly (Fig. 4C).

This arrangement suggests an organizing principle that links retinotopic features to specific DN subpopulations, which, via their distinct VNC connectivity, differentially control motor output. Indeed, the activation of DNg79, endogenously driven by dorsal visual features, causes the fly to lift its front legs above its head (Fig. 4D). Activation of DNp10, which receives most of its input from VPNs tuned to the ventro-lateral and ventro-frontal visual fields, evokes less anterior rotation of the middle legs than DNp07 does (Fig. 2F-G, Fig. 4D). This suggests that the motor circuits targeted by each DN bias reaching movements in the direction of the visual feature that they are most sensitive to. Therefore, these input gradients provide a mechanism to bias the DN ensemble activity towards features detected in particular locations of the visual field and facilitate different landing strategies.

To test how this spatially-biased connectivity could guide context-appropriate landing movements, we recorded the response of DNp10 towards a horizontal bar moving up- or downwards using *in vivo* patch-clamp recordings in flying flies (Fig. 4E-H). We found that DNp10 has a strong, speed dependent preference for bars moving upwards in the ventral visual field (Fig. 4F-G, Fig. S4-1D). Stimulation with the same speed but opposite movement direction resulted in a much smaller depolarization (Fig. 4H) and much lower spike rates (Fig. S4-1D). Hence DNp10 is tuned to objects moving upward in the ventral visual field in front of the fly, adding a directional component matching the spatial tuning of both the sensory input and the motor output of this DN.

Behavioral experiments and decades of research suggest that the sensorimotor pathways controlling landing, including the DN ensemble, should be responsive to frontal looming stimuli. To test whether landing DNs are sensitive to frontal looming stimuli despite their complex spatial tuning and non-overlapping inputs with looming-sensitive takeoff DNs, we recorded from DNp10 during the presentation of frontal looming stimuli mimicking different approach dynamics (Fig. 4I-K). Looming stimuli evoked a rapid depolarization of DNp10 during the approach phase, followed by a repolarization while the stimulus remained at its maximum size. The spike rate increased during the approach, with the majority of spikes preceding the time of collision (Fig. 4I). This spike pattern is suited to drive leg extensions before touchdown. Using three different looming speeds (parameterized by their size-to-speed ratios, (Ache et al., 2019b)), we found that slower looming (higher *r/v* ratios) lead to slightly higher DNp10 spike rates since spikes occur over a longer time window during slower approaches of larger objects (Fig. 4I). Plotting the membrane potential against the angular size (Fig. 4J) or angular speed (Fig. 4K), however, revealed the underlying depolarization is largely independent of the *r/v* ratio. DNp10 encodes a combination of the angular size and speed of approaching objects exceeding 100°/s.

In principle, the parallel organization of sensory streams reaching the takeoff and landing DN Ensembles could enable different response thresholds for the two behaviors, which could be tuned to the requirements of each behavior. To assess how landing DNs might encode looming and how this differs from takeoff DNs, we estimated the angular size threshold of DNp10 to looming stimuli with different *r/v* values using analyses typically conducted for takeoff circuits (Fig. S4-1E). We found that DNp10 exhibits a size threshold of 88° independent of the approach dynamics, which is about twice as large as the size threshold for slow and fast takeoff (von Reyn et al., 2014), suggesting that landing pathways respond later during looming, but still early enough to deploy leg extensions during comparatively slow, voluntary approaches (Fig. 4I). In line with this, landing responses occur about 300 ms later than takeoff responses during the same looming stimuli (*r/v* = 80ms, (Ache et al., 2019a)). The requirements for takeoff DNs is to enable flies to escape rapidly from approaching predatory threats, which benefits from smaller size thresholds leading to earlier responses during looming. Hence, the parallel VPN input streams to landing and takeoff DNs make it possible that these two sensorimotor pathways are tuned to different aspects of the same visual stimulus, matching behavioral requirements.

Taken together, our analysis of the sensory input to takeoff and landing DNs reveal that retinotopic inputs from visual feature detectors tune individual landing DNs to distinct regions of visual space, and their divergent premotor connectivity converts these signals into movements matching the sensory stimulus. This system enables flies to flexibly deploy appropriate landing maneuvers depending on approach angle and environmental context. Despite these specializations, landing DNs strongly respond to frontal looming stimuli, but are tuned to different parameters than takeoff DNs.

### The connectome predicts parallel sensorimotor ensembles for diverse behaviors

The sensorimotor circuits underlying landing and takeoff behaviors appear to have few overlapping elements but surprisingly similar structural organization (Fig. 5A). Both comprise a set of 5-6 DNs that can be differentially recruited by select types of feature-tuned VPNs. In both cases, individual DNs within the ensemble drive variations of the main action in different directions. In the takeoff stream, DNp02 activation drives backward leaning and DNp11 activation drives forward leaning and takeoff (Dombrovski et al., 2023). In these cases, the correct movement direction for escape is hypothesized to be achieved by differential activation of the two DNs having oppositely oriented visual receptive fields along the anterior-posterior axis. As in the landing circuit, the takeoff DN visual receptive fields are established via biased connectivity between the DN and individual members of the presynaptic VPN with differently oriented local receptive fields (Dombrovski et al., 2023; Dombrovski et al., 2025). Given this similarity in organization between the landing and takeoff circuits, we next asked whether all DNs could be grouped into distinct behavioral modules based on similarity of their outputs, defining distinct ensembles across the behavioral space displayed by the fly.

**Figure 5:**
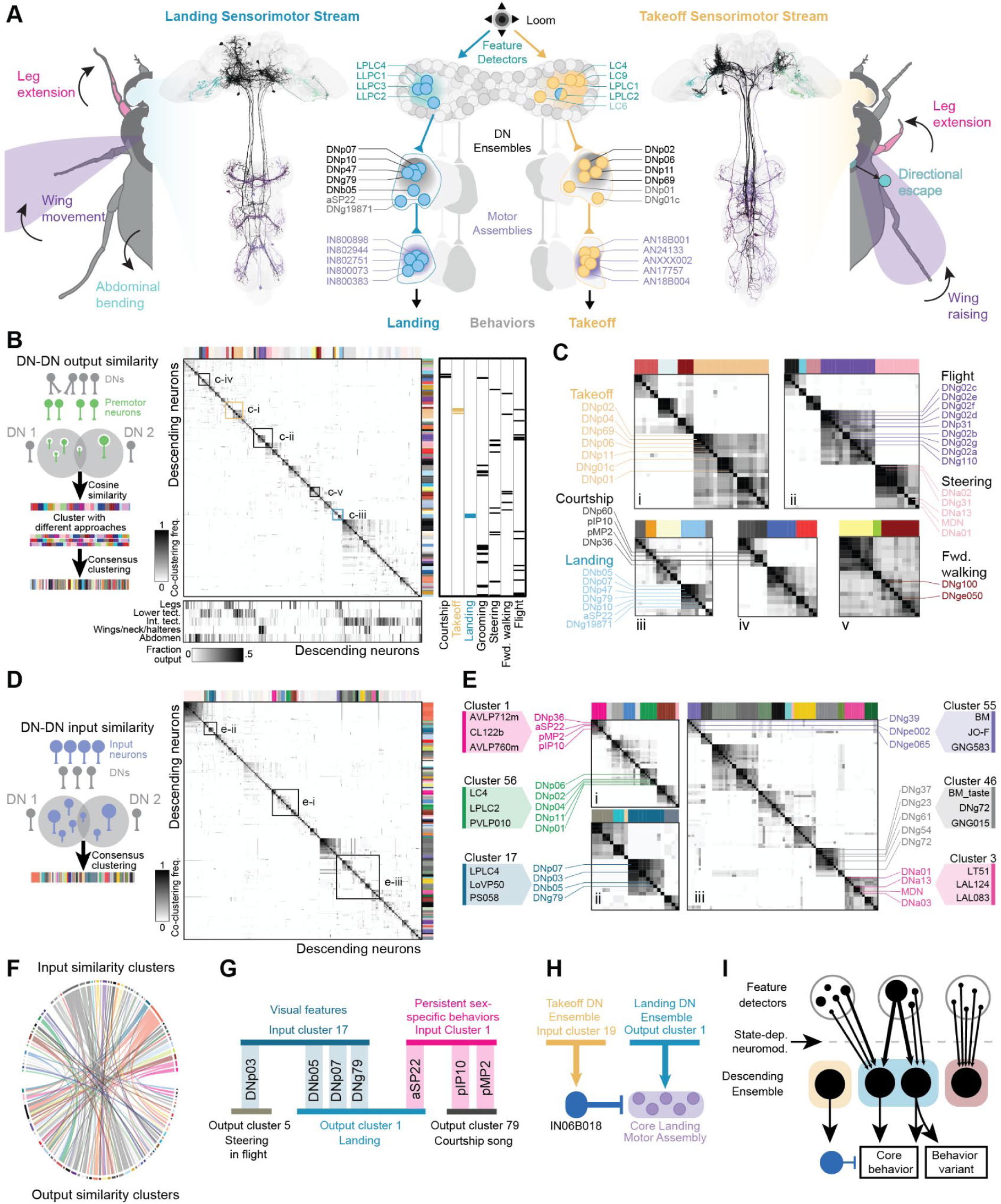
VNC connectivity predicts functional DN ensembles. **A**, Reconstruction of the Landing and Takeoff Sensorimotor Streams, organized as a set of parallel, loosely-overlapping feature detectors, non-overlapping sets of DN Ensembles, and Core Motor Assemblies in the VNC. **B**, Left, Schematic of consensus clustering approach. Middle, All DNs in maleCNS dataset clustered according to their output similarity. Heatmap side panels depict the result of consensus clustering. Bottom, Fraction of each DN’s output in five major VNC regions. Outer right, Annotations for known behavioral functions of clustered DNs. Insets in heatmap are shown in (C) at higher magnification. **C**, DN clusters correspond to groups with known behavioral roles. Heatmaps contain higher-magnification reproduction of insets in (B). Selected DNs that have identified behavioral contributions are labeled. **D**, Cosine similarity for all DNs based on typewise inputs, presented as in (B). **E**, Higher magnification reproduction of insets in (D). Outer colored boxes, top three neurons that provide the most total synaptic input to the DNs in each cluster and also synapse onto each DN in the respective cluster (min. 5 synapses). **F**, Diagram showing how input-defined DN clusters (top) map onto output-defined DN clusters (bottom). Colors reflect input clusters. No input cluster maps uniquely onto a single output cluster. Singleton connections are omitted for clarity (full diagram in Fig. S5-1E). **G**, Landing DNs (output cluster 1) are assigned to multiple input clusters, confirming that different sensory pathways can trigger landing (also see Fig. 3B, S3-3D). aSP22, known for its role in courtship, clusters with the inputs to other courtship neurons including pIP10 and pMP2. **H**, A GABAergic interneuron downstream of the takeoff DNs inhibits members of the Core Landing Motor Assembly. **I**, Summary diagram.

We discovered the Landing DN Ensemble because each of its members had similar output connectivity as other landing DNs. To explore whether all of the fly’s DN types are arrayed into modules comparable to the Landing and Takeoff DN Ensembles, we applied the approach we developed for the landing DNs to the entire DN population. Briefly, we compared output connectivity between all DN pairs using cosine similarity, then grouped DNs using a consensus clustering approach validated by iterative resampling (see Methods). This approach reduces the effect of biases inherent to any particular clustering algorithm by integrating multiple clustering methods and different parameter values (Fig. 5B). Our analysis revealed 81 ensembles of DNs, each of which contained neurons with downstream connectivity highly similar to one another, but distinct from DNs in other clusters (Fig. 5B, Fig. S5-1A-B).

Consistent with these ensembles corresponding to functional behavioral units, DNs that target the same anatomical regions and that are known to actuate similar behaviors each sorted into the same clusters (Fig. 5B, 5C). The Landing DN Ensemble, for example, formed a single cluster (Fig. 5C, left). The DNs that actuate “long-mode” takeoff also grouped together as members of a cluster that additionally contained the Giant Fiber (DNp01), a DN responsible for “short-mode” takeoff (Fig. 5C, left). DNs that are associated with non-ballistic movement sequences such as walking, flight, and courtship also assorted into behavior-specific clusters (Fig. 5C, right) but often formed multiple “cores”, likely due to the complexity of these behaviors. We conclude that the descending premotor circuits of *Drosophila* are organized in parallel, into modular functional ensembles that individually actuate specific motor patterns and collectively span the fly’s behavior space.

To uncover the extent to which the parallel organization of the sensory inputs to Landing and Takeoff DN Ensembles reflects a general organizational principle, we repeated our clustering approach using DN inputs. Again, DNs sorted into clusters (k=90) with high intra-cluster but low inter-cluster similarity (Fig. 5D, Fig. S5-1C-D). We observed clusters of input-sorted DNs marked by convergent input from neurons sensitive to the same sensory modality (e.g., the members of cluster 46, which are all driven by presumed taste-sensitive inputs or cluster 55, driven by mechanosensory neurons). Other input-defined clusters are distinguished by their common functions. For example, DNs involved in locomotion and steering, such as DNa13, MDN, and DNa01 (Bidaye et al., 2014; Chen et al., 2018; Cheong et al., 2025), are all members of cluster 3, and they each receive unilateral input both from visual neurons such as LT51 and neurons from olfactory areas such as LAL124 (Fig. 5E). Takeoff DNs such as DNp01 and DNp11 were driven by the same looming-sensitive VPN types, including LC4 and LPLC2 (Fig. 5E) (Ache et al., 2019b; Dombrovski et al., 2023).

Strikingly, there were no instances of any input-defined DN ensemble that was identical with an output-defined DN ensemble (Fig. 5F, Fig. S5-1E). For instance, aSP22 is engaged by inputs distinct from the other members of the Landing DN Ensemble: when clustered according to its outputs, it joins the other landing DNs, but when clustered according to its inputs, it joins with other sexually dimorphic DNs that control courtship, such as pMP2 and pIP10 (Fig. 5E, G). The input neurons that connect to most DNs in this latter set (input cluster 1) include subtypes of P1, which are sexually dimorphic cells linked to the long-lasting brain states that facilitate persistent sex-specific social behaviors (Fig. 5G) (Anderson, 2016; Clowney et al., 2015; Kimura et al., 2008; Rubin et al., 2025), and other male-specific neurons like AVLP712m and AVLP760m. Other members of the Landing DN Ensemble, including DNp07, DNg79, and DNb05, are driven by the same typewise inputs that also synapse onto DNp03 – a neuron that, when activated, causes flies to avoid oncoming objects in flight (Buchsbaum & Schnell, 2025). This parallel descending architecture enables widespread integration of multimodal sensory inputs, and links them to different motor networks to produce flexible and context-specific motor output. We hypothesize that state-dependent neuromodulation and synaptic gradients provide additional mechanisms to flexibly link sensory features to movement patterns within appropriate contexts.

Consistent with a previous study (Braun et al., 2024), we found that direct DN-DN connectivity tended to be stronger within output-defined clusters than between them, which might be a mechanism to reinforce activity within a cluster and thus promote a specific behavioral output (Fig. S5-1F-G). However, this positive feedback motif was not a prominent feature in most clusters, and many DN-DN interactions crossed ensemble boundaries, suggesting that DN-DN connectivity is not a major driver of functional ensembles (Fig. S5-1F). Unlike serial architectures, parallelized networks can implement inter-ensemble inhibition to refine action selection. While we did not find extensive inhibitory interactions between ensembles on the DN level, we did observe examples of DN output clusters that inhibit the output of other DN clusters. For instance, all takeoff DNs except DNg01c output onto the GABAergic interneuron IN06B018, which directly targets members of the Core Landing Motor Assembly (Fig. 5H). Thus, the initiation of takeoff could instantaneously suppress landing-related leg extensions – which are triggered by similar visual features.

Taken together, our analysis indicates that many descending circuits share important architectural features with the takeoff and landing pathways (Fig. 5I). These DN ensembles are arranged into clusters that provide shared output onto a series of core VNC neurons. These “largely parallel sensorimotor streams” can be efficiently toggled on and off by state-based modulation or inhibitory circuits to facilitate action selection across different time scales. The constituents of each descending ensemble also receive unique input from multimodal feature detectors, allowing the DN population to flexibly initiate and control context-appropriate motor output.

## Discussion

Using a complete adult *Drosophila* CNS connectome together with functional experiments, we investigated the neuronal circuit logic underlying visuomotor transformations. We focused on landing and takeoff behaviors because they are similar in input (frontal looming stimuli) and output (rapid leg extension), but are employed in different contexts. We first identified neurons in the sensorimotor pathway controlling landing, then compared them to the takeoff pathway at each sensorimotor processing layer. We found circuits underlying these behaviors are organized as largely parallel sensorimotor streams, extending from feature detectors to premotor circuits. Each stream comprised a feedforward chain of strongly connected core neurons embedded in a broader network of weaker, but more specific, connectivity enabling finer control (Fig. 5A). Our subsequent analysis of all of the fly’s descending pathways revealed that this organization into largely parallel sensorimotor streams represents a broadly-utilized architecture for behavioral control.

We find that separate ensembles of six to seven excitatory DN types drive landing and takeoff. Previously, these and other rapid actions were attributed to individual “command-like” DNs (Ache et al., 2019a; Bidaye et al., 2014; Braun et al., 2024; Flood et al., 2013; Gao et al., 2013; Hampel et al., 2015; Koto et al., 1981; Kupfermann & Weiss, 2010; Schnell et al., 2017; von Philipsborn et al., 2011; von Reyn et al., 2014). What is the functional advantage of behavioral control by multiple DNs vs. a single command neuron? As a group, DNs within the landing or takeoff ensembles synapse onto premotor assemblies of five to eight neurons in the nerve cord that coordinate the basic movement pattern for each behavior. This function could arguably be served by an individual command-like neuron. However, we found that individual DNs additionally contribute movement variants through distinct connections with motor neurons. For example, DNg79 promotes further dorsal front leg extension in landing, and DNp02 drives postural adjustments that promote backward-directed takeoffs (Dombrovski et al., 2023). Ensemble DNs also differentially connect to abdomen and wing motor neurons, enabling coordinated whole-body adjustments. This is similar to the arrangement of the Mauthner system in fish, where a single DN can activate a rapid escape turn in response to looming or tactile stimulation. Yet, that DN is embedded in an ensemble of other reticulospinal DNs that fine-tune escape direction and together can initiate escape in less urgent circumstances (Eaton et al., 2001; Korn & Faber, 2005).

One possibility for an efficient, serialized architecture could be that the same looming feature detectors drive either landing or takeoff, with the behavioral state determining which DN ensemble is recruited. Consistent with this model, looming stimuli only drive strong responses in landing DNs during flight (Ache et al., 2019a) and state-dependent modulation of sensory processing is widespread in the vertebrate and invertebrate CNS, suggesting this could be a general motif (Maimon, 2011; McGinley et al., 2015). Unexpectedly, however, we found that landing and takeoff DNs receive their primary input from distinct VPN types, indicating that the two pathways diverge upstream of the DN level. Hence, instead of a serial organization that relies heavily on state-dependent circuit modulation for action-selection, the CNS implements a parallel organization from the earliest stages of sensorimotor processing. This parallel architecture enables actions to be differentially tuned to similar sensory features in that downstream DN Ensembles can have different looming size activation thresholds and response latencies to the same approaching object, as appropriate to the requirements of each behavior. Parallel units also provide substrate for behaviors to inhibit each other to enforce a winner-take-all action selection mechanism. It has previously been suggested that DNs promoting one behavior preferentially inhibit DNs promoting another (Braun et al., 2024). Interestingly, we found that the Takeoff DN Ensemble activates a VNC interneuron which inhibits neurons within the Core Landing Motor Ensemble, providing an additional layer of lateral inhibition in the VNC as an action-selection mechanism.

Having identified two parallel sensorimotor streams that control two different behaviors, we investigated if other descending pathways fit this blueprint. Several recent publications sorted DNs into functional groups, sometimes including other sensorimotor layers (Bates et al., 2025; Berg et al., 2025; Braun et al., 2024; S.-y. Takemura et al., 2024; Winding et al., 2023), resulting in broad behavioral assignations. Our approach expands on this, using a consensus clustering strategy to compare pairs of DN types according to the likelihood that they are grouped together using a range of clustering algorithms. Our methodology reveals over 80 DN clusters that vary in size and structure. Some, like the Landing DN Ensemble, include a tight “core” of DNs that frequently co-cluster, surrounded by a looser periphery. Others, such as grooming-related or walking-steering groups, are larger and assemble into a hierarchy with multiple cores, consistent with behavioral requirements for complex movement sequences. Takeoff-related DNs also form a cluster containing several groupings. While most of the neurons in the Takeoff DN Ensemble cluster in a single core, the Giant Fiber (DNp01), a DN associated with a distinct kind of takeoff (von Reyn et al., 2014), locates to a different within-cluster grouping. Hence, the structure of the similarity matrix provides a predictive framework for future functional studies: it describes an organizational logic underlying each behavioral module and also highlights uncharacterized DNs putatively involved in behaviors of interest.

The relationship between these clusters and movement primitives remains to be fully resolved. Broad functional categories such as “flight control” span multiple clusters (Fig. 5B, Fig. S5-1H). But closer examination reveals potential specializations of different clusters. For example, DN types in output cluster 7 drive wing power MNs underlying the basic wing stroke (Fig. 5B, Fig. S5-1H), whereas those in output cluster 5 execute turns via asymmetric excitation of steering wing MNs (Cheong et al., 2025; Schnell et al., 2017), and those in output cluster 9 are proposed to bilaterally drive steering wing MNs to control thrust, lift, or pitch maneuvers (Cheong et al., 2025). Future work should continue detailed quantifications of movement patterns so that the level of behavioral description can be accurately matched to the level of connectome analyses.

Our analysis shows that *Drosophila*, despite its small nervous system comprising ∼165,000 neurons, relies heavily on parallel sensorimotor control streams to integrate sensory stimuli and control motor output across its behavioral space. This differs from the organization in *C. elegans*, which relies heavily on multiplexing (Huang et al., 2023), but instead reflects the sensorimotor organization of vertebrates, whose CNS provides more leeway for redundancies thanks to the large numbers of neurons in each processing layer. Indeed, parallel, distributed processing is widespread in the vertebrate sensory and motor systems (International Brain et al., 2025; Steinmetz et al., 2019; Stringer et al., 2019). Large neuronal ensembles spread across diverse brain areas respond to auditory, visual, chemosensory, and somatic sensory stimuli, hierarchically refining coarsely tuned information from peripheral detectors according to expectations, saliency, brain state, and behavioral context (Fiser et al., 2016; Kato et al., 2012; Kato et al., 2015; Lin et al., 2019; McGinley et al., 2015; Munoz et al., 2017; Niell & Stryker, 2010; Schneider et al., 2018; Smith et al., 2017; Yu et al., 2024). Information from both lower-order and higher-order sensory areas is routed, with exquisite specificity, to “behavioral switchboards”, brain areas containing neural populations distinguished both by their projection patterns and their parallelized control of specific innate behaviors (Gonzalez-Rueda et al., 2024; Zingg et al., 2017). This includes pathways for self-grooming, pup grooming, rearing, different parameters of locomotion (such as turning, stopping, initiation, and speed control), and various aggressive or defensive responses (Capelli et al., 2017; Cregg et al., 2024; Ferreira-Pinto et al., 2021; Goñi-Erro et al., 2023; Kiehn, 2016; Kohl et al., 2018; Stempel, 2024). These premotor regions instruct action selection according to integrated input from sensory and motor planning regions, as well as modulatory projections (Tovote et al., 2016). With the new connectomic and precise genetic tools available in *Drosophila,* we are now able to map complete neuronal pathways from sensory feature detectors to the premotor circuits coordinating motor output across the wide behavioral range displayed by the fly - and make powerful predictions about their functional organization.

## Resource availability

### Lead Contact

Further information and requests for resources and reagents should be directed to and will be fulfilled by the lead contact, Jan M. Ache and Gwyneth M. Card.

### Material availability

Newly generated fly lines are listed in Supplemental Table 1 and are available from the lead contact upon request.

### Data and code availability

The datasets generated during and/or analysed during the current study have been deposited and are publicly available as of the date of publication. Original code for analysis is publicly available as of the date of publication. DOIs are listed in the key resource table. Microscopy data reported in this paper will be shared by the lead contact upon request. Any additional information required to reanalyze the data reported in this paper is available from the lead contact upon request.

## Acknowledgments

Much of this work was done as part of The FlyEM Project Team at Janelia Research Campus in collaboration with the Drosophila Connectomics Group, University of Cambridge, and the associated labs, which were supported by funding from the Wellcome Trust and the HHMI/Janelia FlyEM Project Team. We thank these groups for early access to the maleCNS connectome data.

We thank M. Dickinson and T. Lindsay (California Institute of Technology) for building the Kinefly setup; and Konrad Oechsner and Thomas Walter (both University of Wuerzburg, UoW) for technical support. We thank Greg Jefferis, Alexander Bates (both (MRC Cambridge), Romain Franconville (Janelia Research Campus) for neuprintr package. We thank the FlyLight Project Team and Fly Facility at Janelia Research Campus (Howard Hughes Medical Institute, Ashburn, VA) for split-GAL4 screening and imaging and fly husbandry, respectively. We thank Arthur Zhao for advice and code examples for visualizing columnar input in the optic lobe. We also thank Tess Oram and Han Cheong for contributions to initial connectomic analyses and Katharina Eichler for her help identifying descending neurons in maleCNS and FlyWire.

This work was supported by grants from the Deutsche Forschungsgemeinschaft (DFG, German Research Foundation) to J.M.A. via the Emmy Noether program (DFG AC 371/1-1) and the Next Generation Networks for Neuroscience (NeuroNex) research program, grant C^3^NS (DFG AC 371/2-1). G.M.C. is an investigator of the Howard Hughes Medical Institute. This project was supported by a Janelia Visiting Scientist Project to G.M.C. and J.M.A.

## Author Contributions

Conceptualization: SL, SKA, GMC, JMA Data Curation: SL, SKA, AN, MS, JMA

Formal Analysis: SL, SKA, AN, ME, CJD, JMA Funding Acquisition: GMC, JMA

Investigation: SL, SKA, MS, ME, CJD Methodology: AN, ER

Project Administration: GMC, JMA Resources: GMC, JMA

Software: SL, SKA, CJD Supervision: GMC, JMA Visualization: SL, SKA, AN

Writing – Original Draft: SL, SKA, GMC, JMA

Writing – Review & Editing: SL, SKA, ME, CJD, GMC, JMA

## Inclusion and diversity

We support inclusive, diverse, and equitable conduct of research.

## Ethical compliance

Studies involving *Drosophila melanogaster* are exempt from ethical approval.

## Methods

### Experimental model and subject details

#### Fly husbandry

All *Drosophila melanogaster* stocks were maintained on standard fly food (cornmeal-agar-molasses medium) at 22–25 °C and 50% relative humidity under a 16 h/8 h light/dark cycle. All experiments were conducted on mated females that were used 3-5 days after eclosion. For optogenetic activation of VPNs, larval flies were raised on standard cornmeal food plus 0.2 mM retinal (R2500, Sigma-Aldrich, St. Louis, MO, USA) and switched to standard food plus 0.4 mM retinal on eclosion. For optogenetic activation of landing DNs, one-day-old flies were transferred onto standard fly food containing 300 µM retinal. In both cases, all fly vials were kept in the dark until flies were prepared for experiments.

#### Fly stocks and genotypes

Fly stocks and genotypes used are shown in Supplemental Table 1 and Supplemental Table 3. For split-GAL4 lines, also see section “Split-GAL4 driver identification and characterization”. For optogenetic activation of VPNs and simultaneous expression of GFP in DNp10 neurons a new line was created in this study. Here, we injected VT031084-LexA into the JK22C landing site and made the following recombinant (“I2B”) for DNp10 (on Chr. 2): w; VT031084(JK22C)-LexA, pJFRC 57-13XLexAOp-GFP (su(Hw)attP5);+. We then crossed in UAS-Chrimson (3. Chr.) which result in the final genotype that was crossed with individual VPNs in our patch-clamp experiments: w+; VT031084(JK22C)-LexA, pJFRC 57-13XLexAOp-GFP (su(Hw)attP5); 20XUAS-CsChrimson-mCherry(su(Hw)attP1).

#### Fly tethering

All experiments were performed under dim light at room temperature. Flies were cold anesthetized at 4°C and either mounted with their head and thorax fixed to a pyramid-shaped fly holder (Ache et al., 2019a; Maimon et al., 2010), or with the dorsal side of the thorax fixed to a tungsten wire (Feng et al., 2020) using UV glue. In both cases, the legs and wings were left free of glue allowing for flight and landing responses (see Fig. 1h Fig. 4l). In pin-tethered flies the head could move freely as well. In patch-clamp experiments, front legs were removed to eliminate interference with recordings and visual stimulation (Ache et al., 2019a).

#### Connectomic analyses

Unless indicated, all analyses in the main manuscript were performed on the Janelia maleCNS connectome (v0.9) using the R programming language with the neuprintr and malecns packages. Connectivity diagrams were produced using the igraph and RCy3 libraries for R and Cytoscape (3.10.2). Heatmaps were generated using the gplots package (Warnes et al., 2005). Meshes of neurons and brain regions were downloaded using the malecns package and displayed using the nat toolbox and nat.ggplot.

All analyses were confirmed with parallel analyses in the MANC connectome (v1.2.3, using the malevnc R package) and in FlyWire (v783, using the fafbseg R package). Neurons in maleCNS and MANC are classified according to a hierarchy of annotations based on connectivity, cell body location, and morphology. Individual neurons (bodyIDs) are clustered into “groups” (sets of highly similar bodyIDs, especially bilaterally symmetric pairs, with numeric labels), which are sorted into “types” based on criteria such as common connectivity and serial homology. Type labels are mostly consistent across connectomic resources and across the literature, and reflect an amalgamation of both systematic and non-systematic alphanumeric nomenclature schemes. Group labels can overlap completely with type labels (i.e., a type consists of a single group), especially for paired neurons without serial homology, such as most DNs.

All neurons in this manuscript are referred to according to their type, unless there are compelling differences between neurons of different within-type groups (i.e., only one of the neurons receives substantial input from a presynaptic neuron, or the groups exhibit obvious morphological differences). In this case, we refer to the neuron with the same class-specific prefix used in the type name (e.g., DNg for a gnathal DN, IN for a VNC interneuron, or AN for an ascending neuron) but then substitute the maleCNS group ID for the remainder. We use this alternative nomenclature for several neurons that are mentioned frequently throughout the text, including DNg19871 and IN800073. All bodyIDs for each neuron referred to by a unique name in this paper are described in Supplemental Table 5.

#### Short-range effective connectivity (ShREC)

We sought to measure the signed synaptic influence of a set of presynaptic neurons (source neurons) onto a set of postsynaptic neurons (target neurons). Since we did not feel that we could confidently predict the effects of two or more sign changes (e.g. inhibition of inhibition), we confined our analysis to measurements of monosynaptic excitation, disynaptic excitation, and disynaptic inhibition. Multi-hop synaptic influence is often estimated by multiplying input fractions across synaptic hops. While we have previously utilized this methodology, we believe that it undervalues the impact of multi-hop connections relative to direct connectivity. We instead measured multi-hop synaptic influence by setting a threshold for the recruitment of downstream neurons, then estimated the magnitude of that recruitment. In a sense, this simulates the function of neurons that fire action potentials: they require a certain threshold synaptic input to fire a single spike, and then often increase their firing rate with increasing input, up to a certain maximum firing rate. Briefly, we devised a simple metric that represents this influence of presynaptic input on firing rate as the sum of a weighted vector of synaptic inputs onto the target neurons. Neurons that are more than two hops downstream of the source neurons have a weight of zero, and direct connections from the source neurons have a weight of 1. Neurons that do not use acetylcholine, GABA, or glutamate as a neurotransmitter also have a weight of zero. Neurons downstream of the source neurons and upstream of the target neurons have a weight determined by a logistic function such that neurons receiving a minimum number of inputs (Thresh_Min_) from the start neurons have a weight of .01 and neurons receiving an asymptotic number of inputs (Thresh_Max_) have a weight of 1. Weights for GABAergic and glutamatergic neurons were multiplied by −1, reflecting their inhibitory effect in *Drosophila*. Finally, we then scaled the synaptic inputs to the end neurons by the weight vector and summed the result. These scores were then normalized according to total upstream inputs to each end neuron. For simplicity we employed a common Thresh_Min_ and Thresh_Max_ for all neurons, set to the 90th and 99th percentile of all non-zero edge weights in the Male CNS connectome (10 and 50 synapses, respectively).

For our analysis of VPN output onto descending neurons, “source neurons” were defined as all cholinergic visual projection neurons on the right side of the brain with at least 50 lobular inputs that made at least ten synapses onto any neuron that made at least ten synapses onto any right-side landing DN. ShREC was computed for each of these neurons onto downstream landing DNs (see above). Maps of lobular input were computed for each source neuron based on ROI-wise cholinergic input in each lobula column, with GABAergic and glutamatergic input in each lobular column subtracted. Negative values in each column were set to zero, and lobular maps were normalized by each source neuron’s total lobular input. Finally, to calculate the short-range effective visual receptive field of each landing DN, we took the sum of all lobular maps onto the downstream landing DN, weighted according to effective connectivity of each source neuron.

#### Consensus clustering of descending neurons

Type-wise adjacency matrices of the downstream connectivity of all landing DNs were generated, either un-normalized or normalized according to total output. Cosine similarities between each DN’s output vectors were computed, and these vectors were clustered according to five different methods: spectral clustering, k-medoids clustering, and hierarchical clustering using three linkage methods: Ward’s method, average linkage, and complete linkage. For each method, five k values were specified ranging from 50 to 125. In total, DNs were sorted into 40 distinct sets of clusters.

A consensus clustering approach was used to group DNs according to how frequently they co-clustered with each other. Briefly, each DN’s set of cluster labels was treated as a vector and DNs were compared according to Hamming distance (1 - the fraction of their cluster assignments that overlapped). This DN-DN distance matrix was grouped into 81 clusters using hierarchical clustering (average linkage) since that method produced clusters that were most stable when randomly resampled (see below and Fig. S5-1). This entire procedure was repeated to cluster DNs according to similarity in typewise inputs, resulting in 90 input clusters.

Clustering was evaluated based on how stable assignments were upon random resampling of 80% of the distance matrices. Each clustering specification (clustering method and k value) was resampled 50 times, and stability was computed as the average Rand Index across iterations. We used this technique to compare clustering of the hamming distance matrix using four clustering methods (K-medoids clustering and hierarchical clustering with Ward’s method, average linkage, and complete linkage) and a range of k values from 50 to 100. Our final k values were selected because they represented the lowest values of k within plateaus of mean Rand values (Fig. S5-1).

### Fly behavior experiments

#### Optogenetic activation of VPNs and DNs

The behavioral experiments shown in Fig. 3H-J and Fig. S3-2 were performed with the experimenter blind to the genotype of the flies. Flies were mounted on a pyramid-shaped fly holder illuminated by two infrared (850 nm) LEDs and filmed from the side at 1,000 Hz using an SA4 high-speed video camera (Photron, San Diego, CA, USA) with a resolution of 256 by 256 pixels. For optogenetic activation of VPNs, we used a 625-nm FiberCoupled LED with 1-mm fiber patch cable (Thorlabs, Newton, NJ, USA) pointed directly at the fly from below. The LED was controlled by a T-cube LED driver (Thorlabs), on which the power level was manually selected. The LED was triggered via a data acquisition board (NI-DAQ, National Instruments, Austin, TX, USA), controlled by custom-written MATLAB code (MATLAB 2020b, MathWorks, Natick, MA, USA). We used three different LED power settings (50%, 75%, and 100%) that resulted in different light intensities measured at the position of the fly’s head (59.0, 163.2, and 210.4 µW/mm^2^ respectively). Light pulses were 50 ms in duration with a 10 s interstimulus interval. For each light intensity, at least 5 stimulations per fly were recorded.

For the behavioral experiments shown in Fig. 2, flies were pin tethered and illuminated by three infrared (850 nm) LEDs (SFH4550, OSRAM, Munich, BY, Germany). Videos were recorded from the side and bottom at 150 Hz using two cameras (acA1300-200um, Basler, Ahrensburg, SH, Germany) with a resolution of 1024 by 1152 pixels. For optogenetic activation during flight, 50 ms light pulses with a 10 s interstimulus interval were generated using a 625 nm LED adjusted to output 150 µW/mm^2^. For the alignment of the analog recorded optogenetic activation signal (via a NI-DAQ data acquisition board) and the videos, a 5 ms light pulse was generated at the end of each video using another infrared LED. Wingbeat signals were recorded using a tachometer (University of Cologne, Animal Physiology Electronics Workshop, model 969) (Liessem et al., 2023). Wingbeat signals were detected in MATLAB using the peakfinder function.

#### VPN silencing using Kir 2.1

The behavioral experiments shown in Fig. 3J and Fig. S3-2I-J, were again performed with the experimenter blind to the genotype of the flies (Supplemental Table 3). VPNs were silenced by expression of the inwardly rectifying potassium channel Kir2.1. Flies were pin tethered and mounted in the center of a cylindrical blue LED display (Lindsay et al., 2017; Marie et al., 2016; Reiser & Dickinson, 2008) (470-nm peak wavelength), spanning 216° of the visual field in the horizontal, and 72° in the vertical dimension. Each pixel covered 2.25° of the fly’s visual field in the center of the display. Visual stimulations were constructed as described in Ache et al., 2019b. In brief, exponential looming stimuli with radius to velocity (*r/v*) of 80 ms with smooth edges (using four different brightness values to interpolate the edge pixel intensity) were centered in the fly’s frontal visual field, and bars moving front-to-back unilaterally at 500° s–1. Flies were filmed at 100 Hz from below and both sides. The ventral view was used to analyze landing responses which was defined as the front leg moving into a defined region of interest (ROI)(Ache et al., 2019a), which was adjusted for each fly. Mean ROI intensities were calculated in real time, fed into a digitizer (Digidata 1440A), and recorded at 20,000 Hz. The mean ROI intensity values were lowpass filtered (50-ms time window) and thresholded to acquire timestamps for leg extensions. The threshold was defined as three times the standard deviation for any given trace.

#### Electrophysiology

For *in vivo* whole-cell patch-clamp experiments, flies were handled and prepared as described before (Liessem et al., 2023). In brief, visually guided patch-clamp recordings were obtained from DNp10 neurons by expression of GFP. Experiments were conducted during daylight conditions and at room temperature. Flies were cold anesthetized at 4°C for immobilization, and the head and thorax were fixed to a pyramid-shaped fly holder. This leaves the legs and wings free, allowing flight. The proboscis was fixed with UV glue to increase recording stability. On the posterior side of the head, a small window was cut into the cuticle. Trachea, fat tissue, and the ocellar ganglion were removed to gain access to DNp10, which was visualized under a customized fluorescence microscope based on the SliceScope (Scientifica, Uckfield, UK). The neural sheath covering the location of the neurons was ruptured by local application of collagenase (0.025 % in extracellular saline) using a blunt patch-pipette. During preparation and experiments the fly brain was continuously perfused with carbonated extracellular saline (95% O_2_ and 5% CO_2_) containing: 103 mM NaCl, 3 mM KCl, 5 mM N-Tris (hydroxymethyl)methyl-2-aminoethane-sulfonic acid, 8 mM trehalose, 10 mM glucose, 26 mM NaHCO_3_, 1 mM NaH2PO_4_, 1.5 mM CaCl_2_ and 4 mM MgCl_2_ (adjusted to 273–275 mOsm, pH 7.3) (Gouwens & Wilson, 2009). Whole cell patch-clamp experiments were performed using patch-clamp electrodes (7-12 MΩ) filled with intracellular saline (40 mM potassium aspartate, 10 mM HEPES, 1 mM EGTA, 4 mM MgATP, 0.5 mM Na3 GTP, 1 mM KCl, adjusted to 260–275 mOsm, pH 7.3). Intracellular recordings were recorded in current clamp mode with a MultiClamp 700B amplifier (Molecular Devices, San Jose, CA, USA) and digitized (Digidata 1440A, Molecular Devices) with a 10 kHz low-pass filter and sampled at 20 kHz controlled by pCLAMP 11 Software Suite (Molecular Devices) and corrected for a 13 mV liquid junction potential (Gouwens & Wilson, 2009). Spikes were detected using MATLAB using a custom written spike detector GUI. In brief, spikes were initially identified in a short time window using the peakfinder function and manually inspected. We then used Pearson correlations or Cosine similarity analysis on the full recording to identify events with similar shapes. Subsequently, accuracy of spike detection was confirmed throughout the entire recording, and if necessary corrected within the GUI. Whole-cell patch-clamp recordings had to fulfill the following quality standards to be acceptable for analysis: spike amplitudes needed to be larger than 10 mV, the resting membrane potential had to be below −48 mV, and the seal resistance before breaking into the cell needed to be larger than 6 GΩ.

All optogenetics experiments were carried out using a 625 nm LED for CsChrimson activation adjusted to an intensity of 150 µW/mm^2^. The LED was operated using TTL triggers controlled by preset protocols in pCLAMP. Each recording consisted of 10 activations with a stimulation length of either 50 ms, 100 ms, or 1s and a 10 s inter-stimulus-intervals. The TTL triggers were recorded simultaneously with the membrane potential and wingbeat and stored in abf format (Axon Binary File). For optogenetic stimulations during flight, only flies were considered for further analysis that flew throughout the complete trial. For analysis of non-flying flies, the opposite was true. In both cases, we only took the first 4 trials that met these criteria for further analysis. Spikes were detected as described above and changes in spike frequency were analyzed in different time windows. Spike rate was binned using a bin size of 50 ms from 500 ms before to 500 ms after optogenetic stimulation. Changes in spike rate were determined within 100 ms after optogenetic activation.

For visual stimulations during patch-clamp recordings, flies were centered to a cylindrical green LED display. Exponential looming stimuli with different radius to velocity ratios (r/v, 120, 80, 40) with sharp edges were centered in the fly’s frontal visual field. Before each trial, the pattern was paused at the first frame (4 black dots) for 1 s. After each trial, the last frame (nearly all LEDs black) was shown for 5 s, with an additional 5 s of all LEDs off. Horizontal bars moving from either ventral to dorsal or dorsal to ventral were shown with different speeds (20°/s, 50°/s, 100°/s, and 200°/s). In each trial, the first frame (2 rows of LEDs off) was presented for 1 s until the start of the pattern. At the end of each trial, the last frame (2 rows of LEDs off at the opposite side of the arena) was shown for 5 s with another 5 s of all LEDs off. In both visual stimulation protocols, we only considered experiments for further analysis in which flies were flying throughout the whole stimulation. Here, only the first 4 trials that met these criteria were considered. Membrane potential, wingbeat, and the visual arena controller signals were recorded simultaneously and stored in abf format. Spikes were detected as described above and changes in spike frequency and membrane potentials were analyzed in different time windows. Using a bin size of 50 ms, we evaluated changes in spike rate from 1 s before to 2 s after visual stimulation. For visualization of changes in membrane potential, traces were smoothed (using a 10 ms running window). We calculated the mean membrane potential 100 ms before onset of the stimulus and subtracted this value for baseline subtraction. For quantification, we took the mean membrane potential throughout the stimulus with an additional 100 ms after stimulus offset. The contrast response between two stimuli of the same speed but with opposite direction was calculated as follows: (AUC_up_-AUC_down_)/(AUC_up_+AUC_down_). For quantification of the spike rate, we calculated the mean spikes within the two bins with the highest spike rate.

The instantaneous firing rate (ISF) was calculated for each trial as the reciprocal of the inter-spike intervals (1/ISI) and interpolated onto a continuous time axis, followed by averaging across repetitions per animal. The time point at which the grand mean ISF first reached 50% of its maximum value was defined as the 50% ISF time, and the corresponding angular size (θ) was obtained by interpolating the loom’s time-dependent θ function at that time. The neuronal delay was estimated from the y-axis intercept of the linear fit between time-of-contact (TOC) and r/v, representing the latency between visual stimulus processing and neuronal response onset. This delay was subtracted from the 50% ISF time to estimate the angular size (θ) at the neuronal response onset corrected for processing delay.

### Split-Gal4 driver identification and characterization

Split-GAL4 driver genotypes are listed in Supplemental Table 1. Most lines used in this study have been previously described (Mathew et al., 2023; Nern et al., 2025; Tuthill et al., 2013; Wu et al., 2016). To generate SS02019, we screened the expression patterns of candidate hemidriver combinations (selected based on published GAL4 driver images (Jenett et al., 2012; Pfeiffer et al., 2008; Tirian & Dickson, 2017), as previously described for other split-GAL4 lines, and combined the identified AD and DBD into a stable fly stock (SS02019). We further characterized this driver by imaging the overall expression pattern (using reporter fly line w;;5XUAS-IVS-myr::smFLAG in VK00005, pJFRC51-3XUAS-IVS-Syt::smHA in su(Hw)attP1) and individual cells (using MCFO-1, (Nern et al., 2015)). Images for split-GAL4 screening and characterization were acquired by the Janelia FlyLight Project team using published protocols (Meissner et al., 2023; Zung et al., 2025); step-by-step protocols are available at https://www.janelia.org/project-team/flylight/protocols). Montages of maximum intensity projections of registered (JRC2018U, (Bogovic et al., 2020)) confocal stacks (Fig S4-1B-C) were generated using Fiji (https://fiji.sc/). Overlays of registered images (Fig. 4D and Fig. S4-1A) were displayed using VVDviewer (Kawase, 2025).

### DeepLabCut Tracking

We used the DeepLabCut software (Mathis et al., 2018) to train artificial neural networks that automatically detect the position of different body parts. For each tethering method (head fixed and pin-tethered) and camera perspective used, we trained an independent network. From the side view, antennal basis (AN), pronotum on the thorax (TH), abdominal tip (AB), and the camera facing tarsal tips of the front-(T1), middle-(T2), and hindlegs (T3) were labeled. From the bottom view, AN, AB, and T1-T3 from both sides of the fly were labeled. For the initial training dataset of head-fixed flies, we manually labeled 50 randomly selected frames in 183 videos (9150 frames) from different flies, genotypes and LED intensities. The initial network was trained by this training dataset from the default resnet50 weights for 1030000 iterations. We used this trained model to label all ∼1800 video files and refined the network by extracting and correcting frames using the DLC ‘jump’ method (∼5850 frames in 540 videos) followed by retraining of the neuronal network. The final network consisted of a total of ∼15000 manually labeled frames. Since videos acquired in pin-tethered flies had a higher spatial but a lower temporal resolution, less frames were necessary for DLC training. The final network after outlier correction to track the fly from the side consisted of ∼2500, the one tracking the fly from below of 3000 manually labeled frames. Moreover, we manually inspected and refined labels in flies tracked from below and the side at 150 Hz in 100 ms before and after optogenetic activation (∼55.500 frames) since determination of leg angles with a lower frame rate is more prone to outliers. Labels were exported as csv files and further processed in MATLAB. For simplicity, we mirrored the labels of the left legs along the body axis and used the mean position of both leg pairs for further analysis. For quantifications, such as determination of the maximum Euclidean distance or the landing probability, leg trajectories were smoothed either by 2 frames (in videos with 150 Hz) or 5 frames (in videos with 1000 Hz).

In all behavioral experiments in which we tracked the flies using DLC, trials were only considered for analysis if flies were flying throughout the whole video. Flies were only considered for analysis if at least 4 trials per condition (e.g., LED intensity) were acquired. Trials in which flight cessation or flailing occurred prior to optogenetic activation were excluded.

For further analysis, videos from both cameras were set to the same scaling factor (0.9 for sideview camera). For this, we used the distance of the DLC labels in the same frame in both videos (random frame prior to stimulation) for the front and hind legs. Flies were furthermore normalized to the size of their thorax (TL) for several reasons, one being that the thorax is the only body part of the fly that is immobile in both head- and pin-tethered flies throughout the complete video. The TL was acquired from a randomly selected frame in all analyzed videos from the side view and was determined by the position of the neck and pronotum (Fig. 1D). Additionally, the TL was used to compensate for minor differences in mounting angles of the flies by converting DLC labels from global coordinates into a thorax centric coordinate system. This is very reasonable because all tracked legs are attached to and move in relation to the immobilized thorax.

Leg trajectories and Euclidean distances (*d*) were determined in relation to the “resting position” during flight, which is the mean position of the leg over 100 ms prior to stimulation. If not otherwise indicated, leg trajectory and *d* are plotted as mean responses across all trials of a fly. The maximum Euclidean distance (*d*_max_) was determined for each leg within 400 ms after optogenetic activation, if not otherwise indicated. We analyzed the trajectory of T1-T3 during (50 ms) and after (400 ms) optogenetic activation relative to their resting position, but also showed longer time windows (for example in Fig. 1G).

One criterion for landing in insects is the simultaneous extension of all legs, with the front legs reaching beyond the head (Borst, 1986; Goodman, 1960; Tammero & Dickinson, 2002). In the first set of experiments (DNs screen), leg extensions were normalized to DNp07_1, which is one of the landing DNs described previously (Ache et al., 2019a). In the second set of experiments (VPN screen), responses were normalized to LLPC2/3_1. LLPC2/3 reliably drove landing responses when activated, and the neurons were shown to be part of the landing circuit, i.e., they receive input from looming sensitive neurons and are monosynaptically connected to two landing DNs (Ache et al., 2019a). We therefore scored a landing response if T1d_max_ was higher than 50% of the mean T1d_max_ of all flies tested, in which we were stimulating either DNp07_1 or LLPC2/3_1 neurons with 100% LED intensity.

Changes in body curvature/posture were determined with the tracked TH and AB segments. We therefore determined the mean angle of AB in relation to TH, which is a fixed point of the fly that can not move, within 100 ms before stimulation. We then determined the changes in abdomen angle over 800 ms after stimulation (bin size 50 ms). For quantification, we took the mean differences of all trials per fly within the first 80 ms after optogenetic stimulation.

Leg angles were determined perpendicular to the body axis starting from the mean resting position. For the probability distribution shown in Fig. 2F-G (bin size, 2°), changes in leg angles were plotted for the stimulus duration (50 ms) and 80 ms after optogenetic stimulation, since we only wanted to capture the initial leg responses. Differences in leg angles for DNp10 for the last third of the analysis window and the maximum Euclidean distance to rest within the complete analysis window were analyzed using one-way ANOVA with Tukey–Kramer post hoc correction for multiple pairwise comparisons.

### Quantification and statistical analysis

Data analysis was performed in MATLAB. Statistical tests used and p-values are described in Supplemental Table 2 and the main text, the figures, or the figure captions. If not otherwise noted, values are presented as medians and IQR throughout the text and figure legends. Details about the time windows used for quantification are given within the detailed description of the individual experiments.

### Leaky integrate-and-fire computational model

To predict the effects of VPNs on the activity of the landing DNs, we ran a leaky integrate-and-fire model (Shiu et al., 2023) on the FlyWire connectome (Dorkenwald et al., 2024; Schlegel et al., 2024). Model details are described in Shiu et al. (Shiu et al., 2023). In short, the model uses the entire connectivity of the brain and assigns positive or negative weight to each connection based on the number of synapses between neurons and the predicted neurotransmitter (Eckstein et al., 2024) of the presynaptic neuron. We adopted the simulation parameters from (Shiu et al., 2023), including the assumption that acetylcholine has an excitatory effect and GABA and glutamate have an inhibitory effect. The intrinsic spike rate of the neurons in the model is zero. The VPNs of interest on the left side of the brain were stimulated by adding a Poisson spike train of defined frequency. We stimulated the neurons of interest at 200 Hz for 1 s per trial. We simulated 30 trials per condition. The resulting spike rate of all landing DNs was averaged across the 30 trials per condition.

## Supplemental Information

### Supplemental Figures

**Fig. S1-1:**
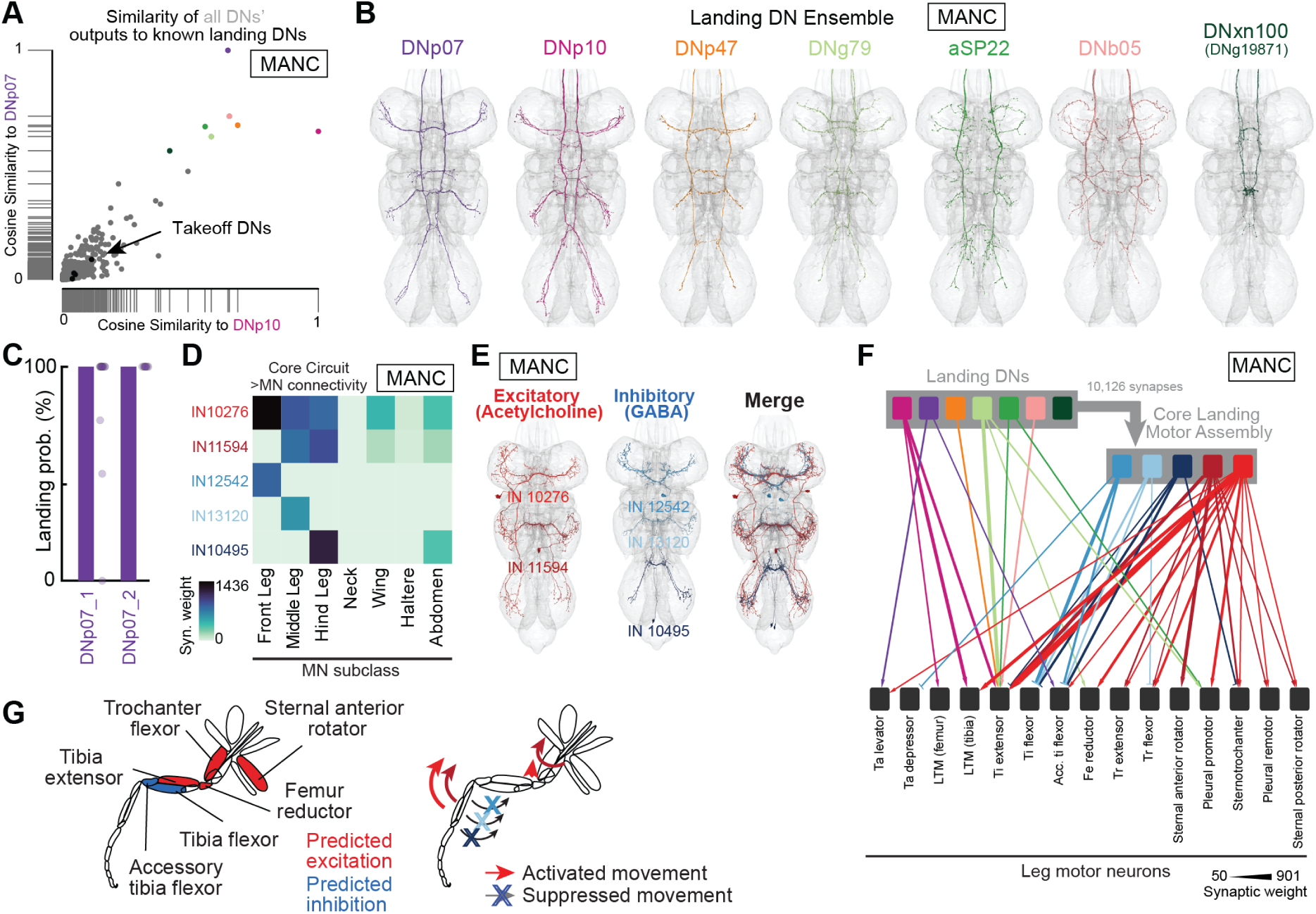
Landing DNs and Core Landing Motor Assembly in MANC. **A**, Cosine similarity of synaptic connectivity of all DNs to DNp07 and DNp10. **B**, Members of the putative Landing DN Ensemble. **C**, Percent of flies that perform landing responses (based on front leg extensions greater than dashed line from Fig. 1E) using two different driver lines for DNp07. **D**, Synaptic connectivity between the Core Landing Motor Assembly and leg, upper tectulum, and abdominal MNs. **E**, Morphology of the Core Landing Motor Assembly, the five neurons that receive substantial input from 5 of the 6 landing DNs, in MANC. **F**, Diagram depicting connectivity between landing DNs, Core Landing Motor Assembly, and leg MNs based on data from MANC. **G**, Predicted activation (red) or inhibition (blue) of leg muscle groups by the activity of the Core Landing Motor Assembly. All connectome analysis in this figure is based on MANC.

**Fig. S1-2.**
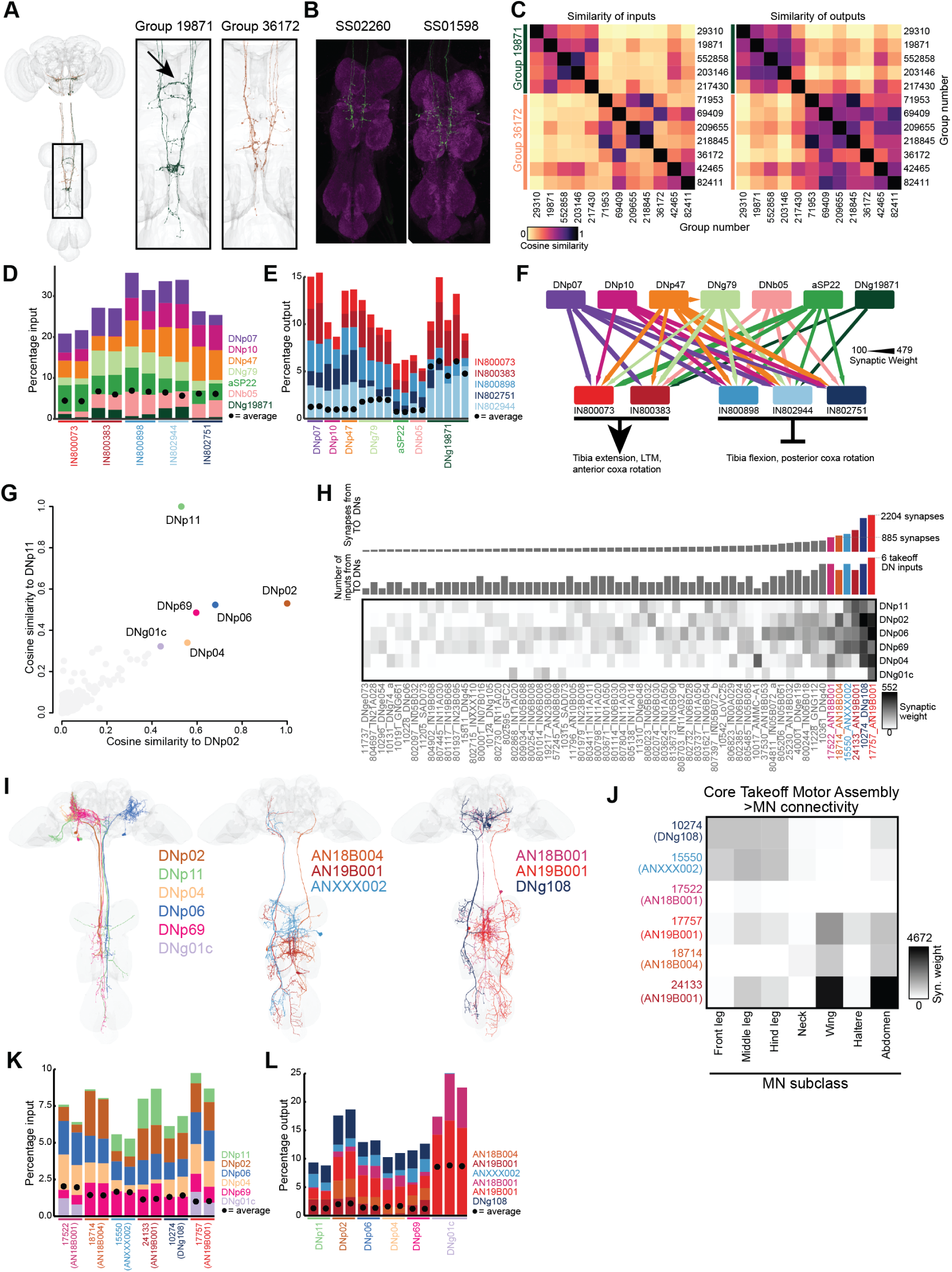
DN ensembles and core motor assemblies controlling takeoff and landing. **A-C**, Neurons belonging to type DNg06 comprise two groups. **A**, Morphology of all bodyIDs with type “DNg06.” Neurons are colored according to group. Arrowhead indicates a trans-midline tract unique to DNg19871. **B**, Images of split-GAL4 lines known to express in DNg06, (from Flylight Project, Janelia Research Campus). All known lines exhibit morphological similarities to group 36172, not group 19871. **C**, DNg06 groups 19871 vs. 36172 have distinct input (left) and output connectivity (right). Heatmaps depict cosine similarity of bodyID-wise connectivity. **D**-**F**, The Landing DN Ensemble converges onto a five-neuron “Core Landing Motor Assembly”. **D**, Percentage of synaptic input to each member of the Core Landing Motor Assembly that arrives from each landing DN. Circles reflect the mean input to an Assembly neuron from an average set of neurons, that is the average number of synapses each neuron receives across all individual inputs, multiplied by 19 (the number of landing DNs). Here, total input is the sum of all synaptic input from other neurons. **E**, Percentage of synaptic output from each landing DN onto each member of the Core Landing Motor Assembly. Circles depict mean weight of all outputs from each landing DN, multiplied by 10 (the number of landing DNs). Total output is the sum of all synaptic output to other neurons. **F**, Groupwise connectivity of landing DNs onto Core Landing Motor Assembly. Arrowheads, cholinergic synaptic connections. T-bars, glutamatergic/GABAergic synaptic connections. Minimum syn. weight threshold, 100 synapses. **G-J**, Neurons belonging to the Takeoff DN Ensemble and the Core Takeoff Motor Assembly. **G**, cosine similarity of all DNs to DNp02 (x axis) and DNp11 (y axis) as in Fig. 1B. Highlighted neurons, DNs similar to both DNp11 and DNp02. These comprise the Takeoff DN Ensemble. **H**, Connectivity from Takeoff DN Ensemble onto all downstream neurons that receive 50 or more inputs from two of those DNs. Highlighted bars, neurons that receive >800 synapses from all takeoff DNs and ≥50 synapses from ≥4 of the takeoff DNs. These comprise the Core Takeoff Motor Assembly. Rows are labeled by [neuron group]_[neuron_type]. **I**, Morphology of the neurons in the Takeoff DN Ensemble (left) and the Core Takeoff Motor Assembly (middle, right). **J**, Direct output of the Core Takeoff Motor Assembly onto MNs binned according to subclass, as in Fig. 1I. **K**, Percentage of synaptic input to each member of the Core Takeoff Motor Assembly, as in D. **L**, Percentage of synaptic output from each takeoff DN onto each member of the Core Takeoff Motor Assembly, as in E. All connectome analysis in this figure is based on maleCNS.

**Fig. S2-1:**
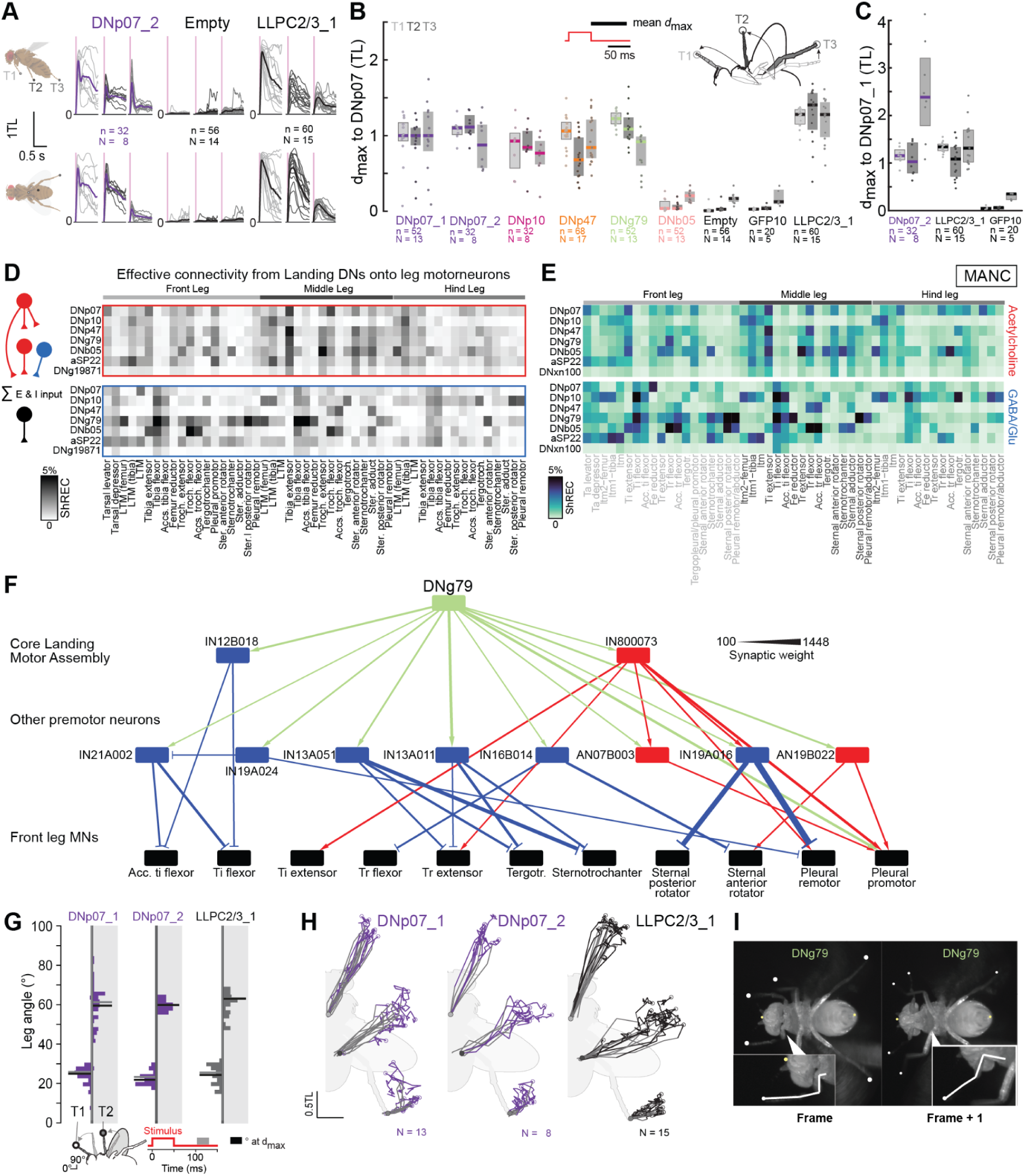
Further differences in landing DN-driven leg kinematics and DN-MN connectivity in MANC. **A**, Movement amplitude (Euclidean distance from resting position) of front, middle, and hind legs during DN activation. Colored line, grand mean of all flies. Gray lines, averages per fly. Activation of DNs and VPNs was conducted in different setups, hence one VPN line (LLPC2/3_1) was included with the DNs for comparison. **B-C**, Quantification of the maximum Euclidean distance of T1-T3 normalized to DNp07_1. **B**, quantification of leg movements driven by activation of all neuron types (based on bottom view). **C**, quantification of leg movements driven by additional neuron types (based on side view). **D-E**, Heatmaps showing effective connectivity (top, direct and disynaptic excitation; bottom, disynaptic inhibition) between each landing DN and MNs driving different leg muscles in the maleCNS (D) and the MANC connectome (E). MNs sorted and colored according to their target leg. Heatmaps display values between 0 and 5%: for (D) maximum inhibitory value 8.4% (DNb05:FL Tr. Flex.), maximum excitatory value 4.8% (DNb05:ML Tr. Ext.); for (E) maximum inhibitory value 9.1% (DNg79:FL Sternal posterior rotator), maximum excitatory value 4.5% (DNb05:HL Tr. Flex.). **F**, Connectivity of DNg79 onto Core Landing Motor Assembly, MNs innervating the front leg, and other INs and ANs upstream of those MNs. All neurons grouped typewise. Red, cholinergic neurons. Blue, GABAergic or glutamatergic neurons. **G**, Probability distribution of T1 and T2 leg angles (bin size 2°) relative to resting position, measured along the anterior-posterior axis/midline within the last third (grey bar) of the analysis window (80 ms, below). Gray line, median leg angle from each fly within that analysis window. Black line, median leg angle at maximum Euclidean distance from rest from each fly within the full analysis window. Grey and black asterisks, significantly different (one-way ANOVA) from DNp10 in the respective time windows. **H**, Ventral view of the fly with superimposed trajectories of the tarsal tips during (grey) and after (black) optogenetic activation of the two DNp07 lines used and LLPC2/3_1. **I**, Two exemplary consecutive frames from the activation of DNg79. Left half, T1 reaches to the front of the fly similarly to activation of the other DNs. Right, Subsequently the coxa is moved anteriorly which is a unique movement sequence only observed during DNg79 activation.

**Fig. S2-2:**
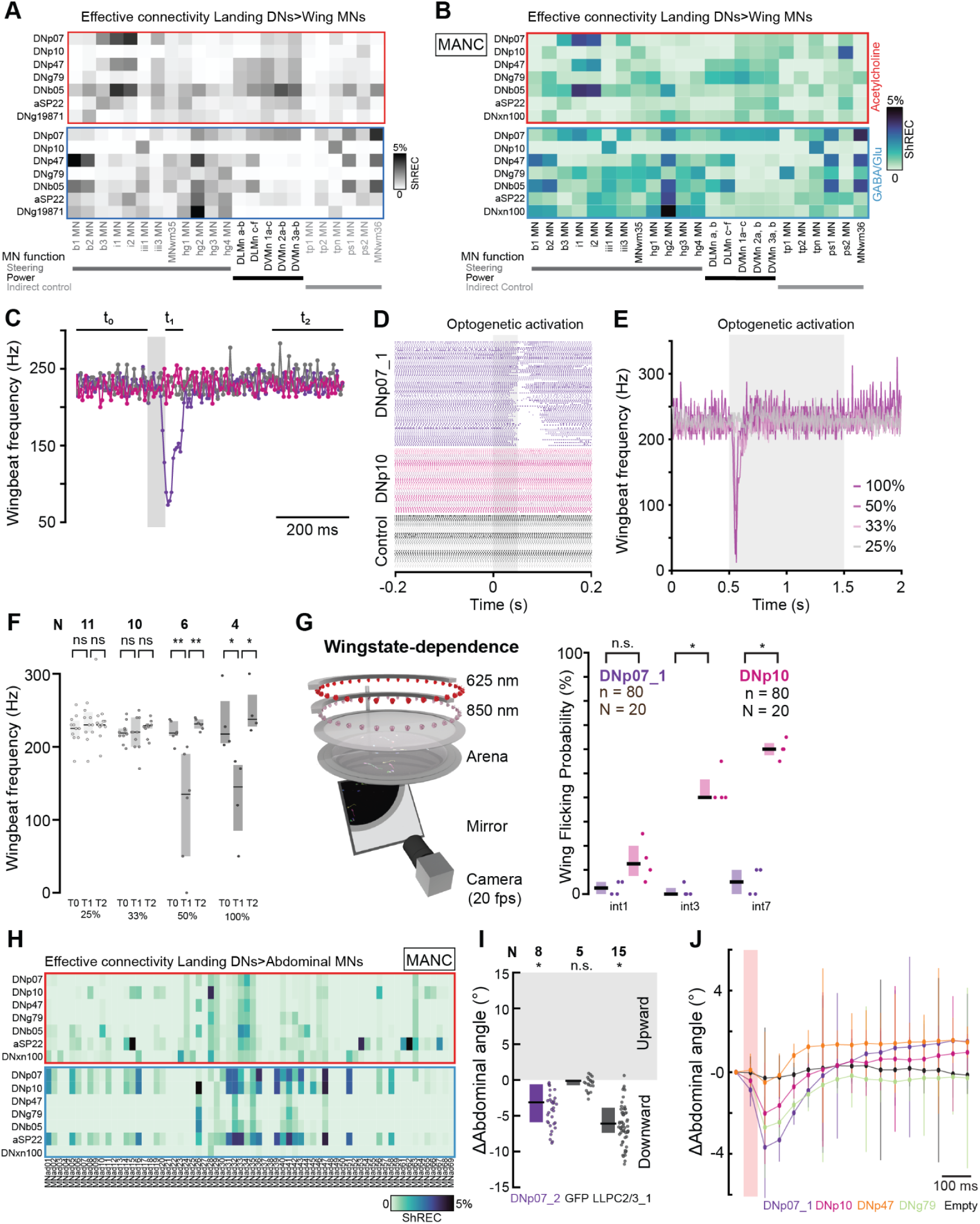
Members of the Landing DN Ensemble exert differential control on wing and abdominal MNs. **A-B**, Heatmap showing effective connectivity (top, direct and disynaptic excitation; bottom, disynaptic inhibition) between each landing DN and MNs driving different wing muscles in the maleCNS (A) and the MANC connectome (B). Heatmaps display values between 0 and 5%, for (A): maximum inhibitory value 8.1% (DNg19871:hg2 MN), maximum excitatory value 4.0% (DNb05:i1 MN); for (B): maximum inhibitory value 10.5% (DNxn100/DNg19871: hg2 MN), maximum excitatory value 3.74% (DNb05: i1) **C**, Mean wingbeat frequency of all flies during optogenetic activation (shaded area) of DNp07, DNP10, and empty controls. Bin size, 5 ms. **D**, Wingbeat events for all flight trials after optogenetic activation (shaded area) of DNp07, DNp10, and control flies (color coded by genotype and all trials of a fly). **E**, Mean wingbeat frequency over time after optogenetic activation (1 s) of DNp07_1 with different LED intensities. **F**, Quantification of (E). **G**, Wing flicking probability in flies walking in an open field during optogenetic stimulation of DNp07 and DNp10 with different intensities. **H**, Heatmaps showing effective connectivity (top, direct and feedforward excitation; bottom, feedforward inhibition) between each landing DN and MNs driving different abdominal muscles in the MANC connectome. Heatmaps display values between 0 and 5%: maximum inhibitory value 7.4% (DNp10:MNad26), maximum excitatory value 6.3% (aSP22:MNad15). **I**, Quantification of abdominal angle changes after optogenetic activation for additional genotypes. **J**, Change of abdominal angle over time plotted as grand mean and standard deviation for each genotype (bin size 50 ms). All p-values calculated via Wilcoxon rank sum tests.

**Fig. S3-1:**
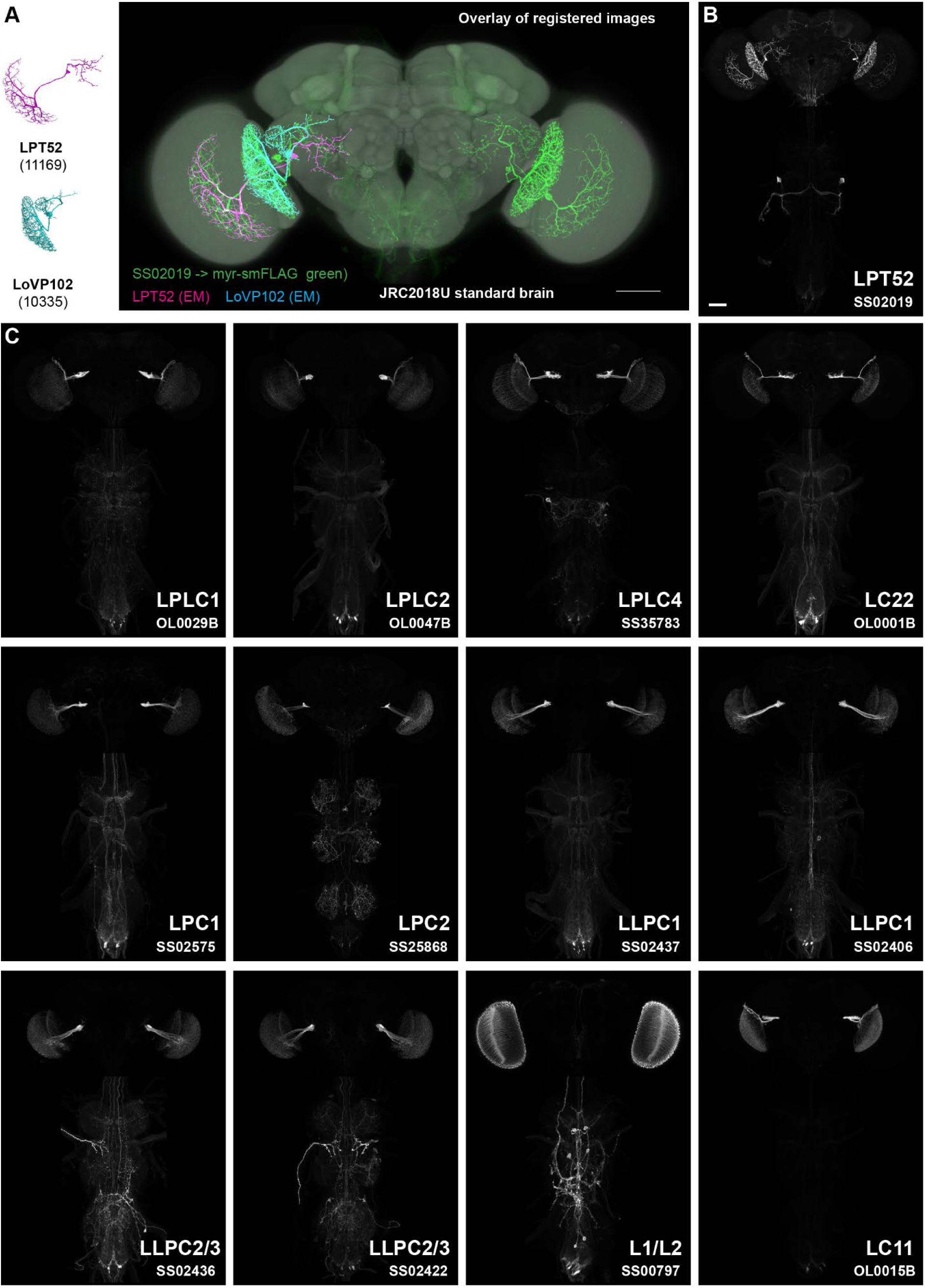
VPN driver lines used for activation and silencing screen. **A-B**, Expression pattern of SS02019. **A**, Overlay of registered images showing the SS02019 expression pattern (myr::smFLAG membrane marker, green), EM reconstructions (also left with male CNS bodyIDs) of LPT52 (magenta) and LoVP102 (cyan) and the JRC2018U template brain. **B**, Composite of maximum intensity projections of registered confocal images of the brain and ventral nerve cord. Genotype as in (A). Brain and VNC are of the same specimen but were imaged separately. Scale bar represents 50 µm. **C**, Images of additional driver lines (myr::smFLAG reporter, imaged and displayed as in (B). Most panels are generated from previously published confocal stacks. Scale bar in (B) also applies to (C). Image stacks are available at www.janelia.org/split-GAL4 (some pending for release).

**Fig. S3-2:**
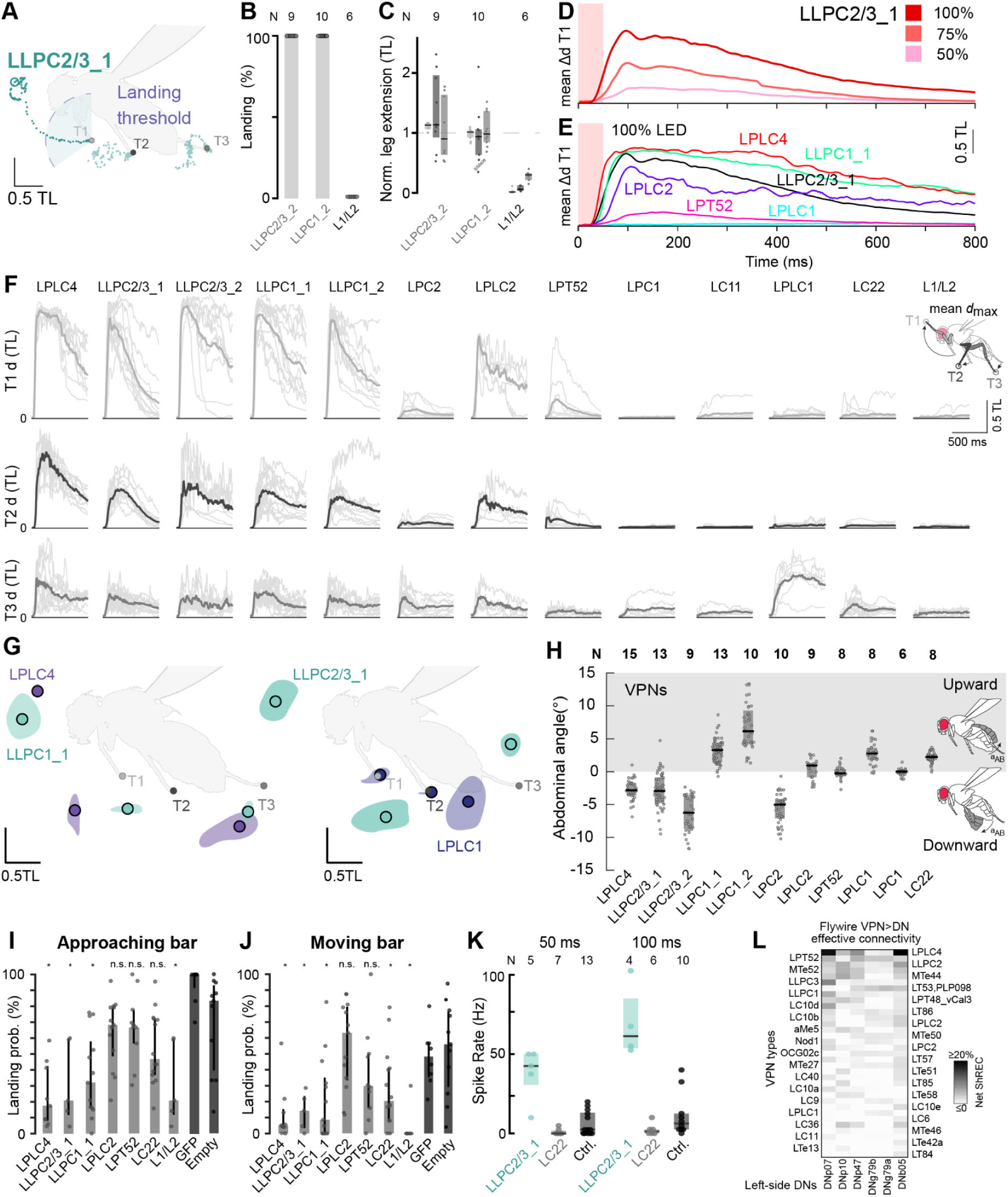
Several VPN populations are necessary and sufficient for landing and drive unique body movements. **A**, Leg trajectories of single flies after optogenetic activation of LLPC2/3_1. Dashed line, landing threshold based on 50% of the mean maximum Euclidean distance to rest from all flies of LLPC2/3_1. Open circle, max. Euclidean distance to rest in that particular fly. **B**, Quantification of landing in additional fly genotypes, quantified as movements greater than dashed line from (Fig. S3-2A). **C**, Mean maximum Euclidean distance of the front, middle, and hindleg, normalized to LLPC2/3_1 responses. Dots, single fly averages. **D**, Change in Euclidean distance of T1 after optogenetic activation (shaded area) of LLPC2/3_1 with different LED intensities. **E**, Change in Euclidean distance of T1 after optogenetic activation (shaded area) of different VPNs with full LED intensity. **F**, Mean Euclidean distance of T1-T3 (grayscale) to resting position of all trials. Bolt line, grand mean of all flies. **G**, Probability density distribution (50% cutoff) of the mean maximum Euclidean distance from rest of all trials and from all files of the respective genotype 400 ms after optogenetic activation. Solid circle, grand mean. **H**, Change in abdomen angle after optogenetic activation of different VPNs. **I-J**, Landing probability of VPN silenced flies using Kir2.1 in response to an approaching or moving bar. **K**, Spike rate of DNp10 during 100 ms of optogenetic activation of LLPC2/3, LC22, and controls. All p-values calculated via Wilcoxon rank sum tests (see Supplemental Table 2). **L**, Net ShREC between VPNs and DNs, as in Fig. 4F, calculated using the FlyWire connectome. Heatmap displays values between 0 and 20%: actual maximum 22% (LPLC4:DNb05), actual minimum −3.2% (LT52:DNp10).

**Fig S3-3.**
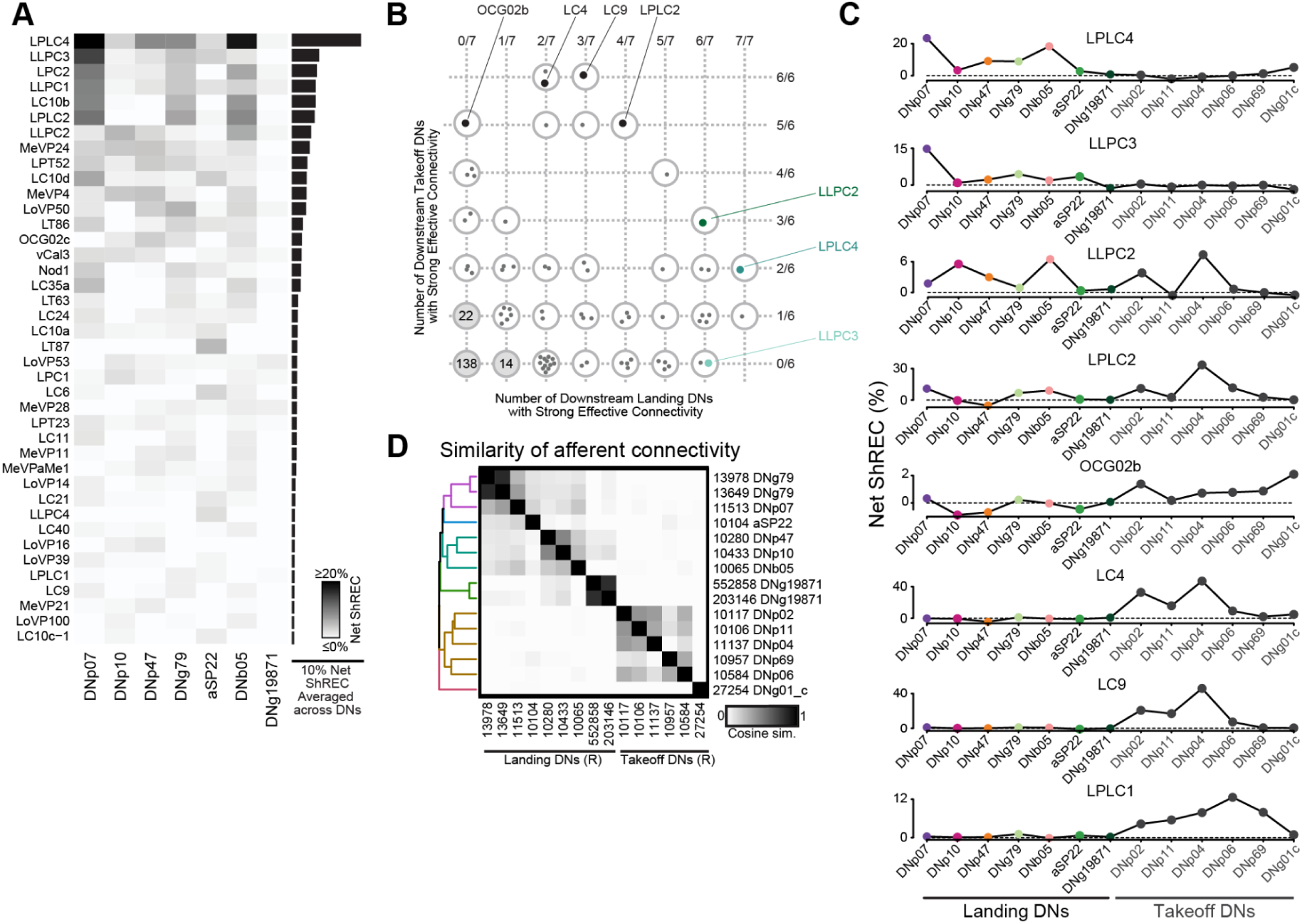
Landing and Takeoff Descending Ensembles receive distinct VPN inputs. **A**, Net ShREC from VPNs to landing DNs. Heatmap is limited to the top 40 best-connected VPNs, as measured by average net ShREC from that VPN to all landing DNs. Heatmap is sorted by the same value, which is plotted in the bar plot (right). Heatmap displays values between 0 and 20%, actual minimum −2.3% (LT82b:DNp11), actual maximum 47.1% (LC4:DNp04). **B**, Scatter plot of the number of VPN types that have a ShREC ≥0.4% with given numbers of landing DN types or takeoff DN types. Each circle contains a point for each VPN types that falls into that category, unless >12 VPN types qualify (in which case the number in that circle reflects the number of VPNs that meet that criteria, e.g., there are 138 VPNs that do not have ShREC ≥0.4% with a single takeoff or landing DN). Example neuron types are highlighted. **C**, ShREC values for eight example VPNs onto each landing and takeoff DN, plotted as in Fig. 3N. **D**, Cosine similarity of direct upstream inputs between landing DNs and takeoff DNs from the right hemisphere, hierarchically clustered using average distance as a linkage method. Colors in dendrogram highlight six DN input clusters.

**Fig. S4-1:**
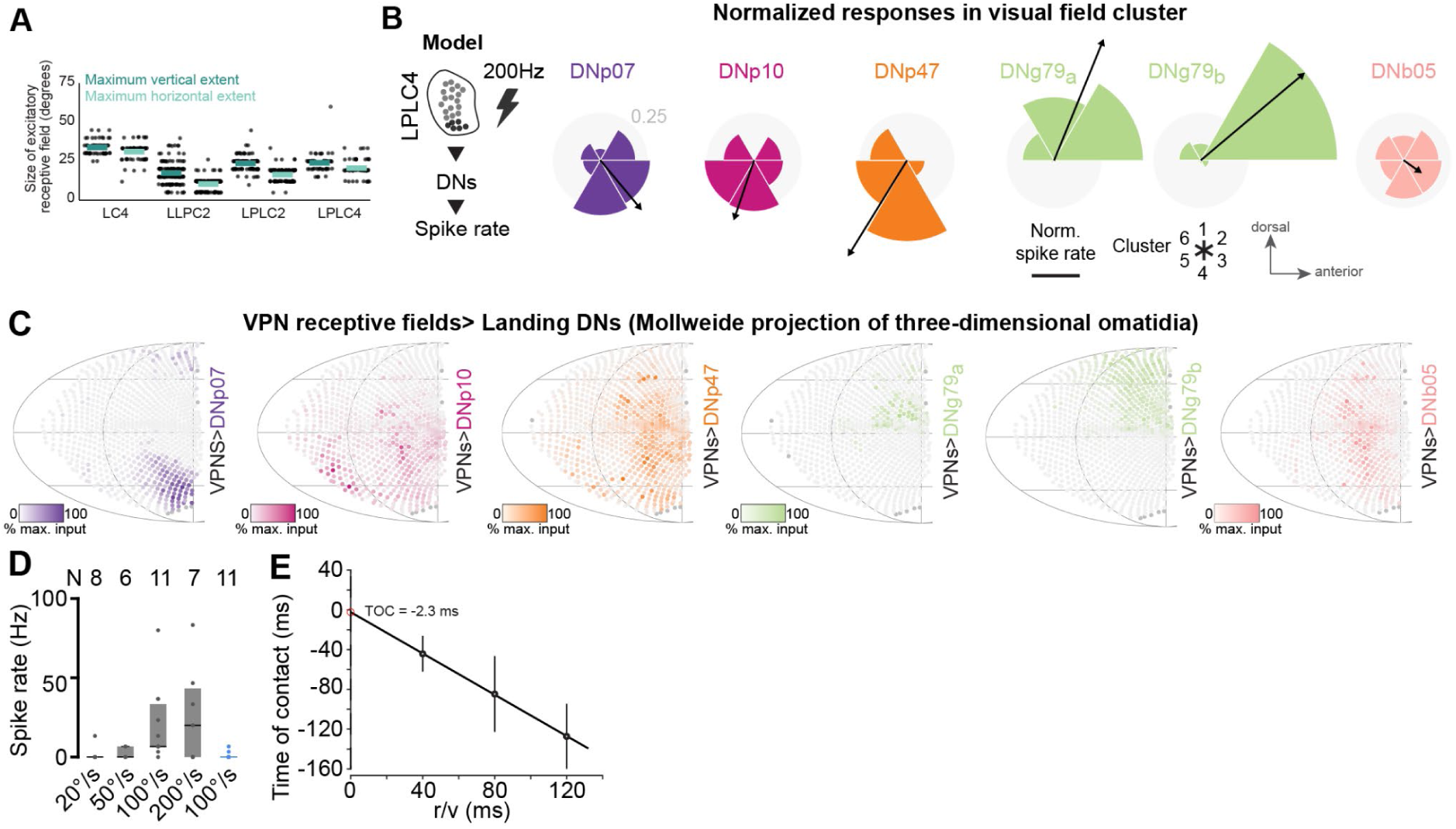
Individual landing DNs have different spatial sensitivities. **A**, Predicted spatial tuning of three VPN types (right hemisphere only). For each VPN, an approximate spatial receptive field was computed. Each lobula column from the right hemisphere was considered to contribute to the neuron’s receptive field if the number of cholinergic inputs to that neuron within the column exceeded the summed GABAergic and glutamatergic input in that column by 0.5% of the neuron’s total synaptic input. The width (height) of the receptive field was measured as the maximum extent of the receptive field within a single horizontal (vertical) line of lobula columns. Width estimates were scaled by √2 to account for the interommatidial distance of the eye’s hexagonal lattice, and inter-columnar intervals in both dimensions were multiplied by 5°, an approximation of how much of a fly’s visual field is represented by a single ommatidium in 1 dimension. Each point reflects the tuning of a single neuron of the indicated type and dimension. Horizontal lines indicate mean. **B**, Connectome-constrained computational model simulating recruitment of landing DNs in response to activating LPLC4 clusters from (Fig. 4B) (at 200 Hz). The resulting spike rate in landing DNs was normalized to the sum of all clusters. For DNg79, responses for both neurons on the left side of the brain are shown. Arrow, directional vector. Grey shaded area and scale bar, 25% of the normalized spike rate of all clusters. **C**, Visual maps of retinotopically tuned input to landing DNs derived from Male CNS connectome. Maps show summed net ShREC of all upstream VPNs onto each right-hemisphere landing DN, weighted by the column-wise cholinergic inputs to those VPNs and normalized to the visual column receiving the most input. Negative values are forced to zero. Data are depicted as a Mollweide projection of the fly’s compound eye, with each point representing a specific ommatidium/viewing direction. **D**, Mean spike rate during upward (magenta) and downward (black) moving bar stimuli with different speeds. Spike rate was calculated as the average of the two bins (bin size 50 ms) with the highest firing rates during stimulation. **E**, Time of contact (TOC) plotted as a function of *r/v* (size/speed) (von Reyn et al., 2014), showing a linear relationship between stimulus expansion rate and neuronal response timing; the y-intercept represents the estimated neuronal delay (red dot).

**Fig. S5-1.**
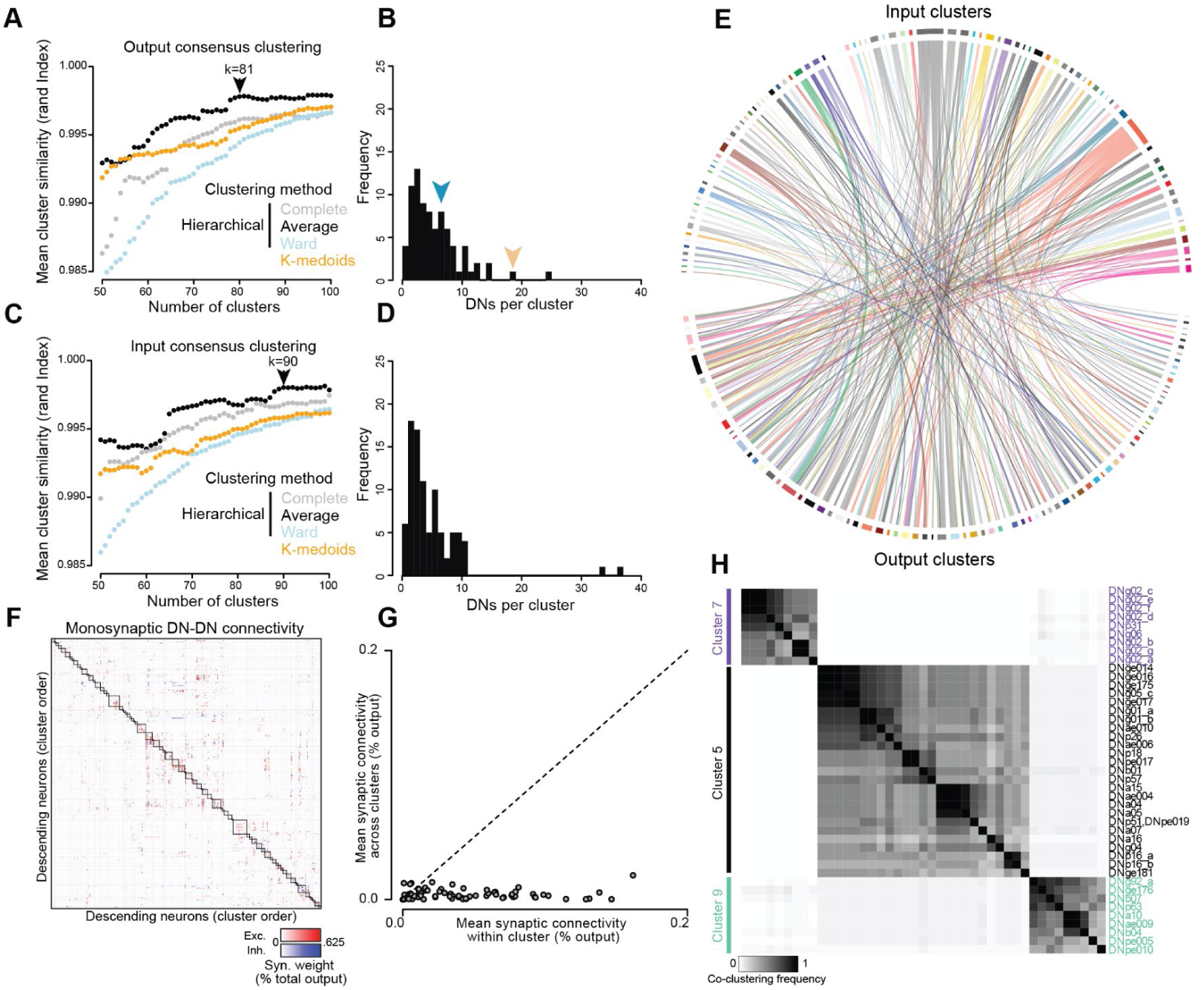
DNs cluster into ensembles based on both input and output connectivity. **A**, Resampling-based validation of clustering methods for assigning clusters based on DN outputs. For each clustering method (colors) and for K values ranging from 50 to 100 (x-axis) we assessed the similarity of cluster assignments when 20% of the neurons were held out (see Methods). Y-axis, average similarity across 50 repetitions. Arrowhead indicates the lowest K value to achieve asymptotic performance. **B**, Histogram of DNs per cluster for output clustering. Arrowheads, DNs per cluster for landing (left) and takeoff (right) clusters. **C**, Clustering performance as in (A) based on DN inputs. **D**, Histogram of DNs per cluster for input clustering. **E**, Chord diagram showing un-thresholded mapping between input clusters and output clusters. **F**, DN to DN synaptic connectivity, normalized according to DN output. DNs are sorted according to cluster identity from Fig. 5B, boxes indicate neurons belonging to the same cluster. Cells are colored (red or blue) according to the neurotransmitter released by the presynaptic DN (acetylcholine or glutamate/GABA, respectively). **G**, For each cluster, plot of mean synaptic connectivity between DNs in that cluster (x-axis) versus mean synaptic connectivity between DNs in that cluster and DNs and other clusters (y-axis). DNs have significantly stronger connectivity with other DNs belonging to their own output cluster than with DNs in other clusters (paired t-test, *p*<0.0001). **H**, Higher-magnification visualization of three non-consecutive output clusters that each contain DN types involved in distinct flight-related actions.

### Supplemental Tables

**Supplemental Table 1:** Reagents and resources.

| REAGENT or RESOURCE | SOURCE | IDENTIFIER |
| --- | --- | --- |
| **Antibodies** |  |  |
|  | N/A | N/A |
| **Chemicals, peptides, and recombinant proteins** |  |  |
| All-trans-retinal | Sigma-Aldrich, St. Louis, MO, USA | R2500 |
| Collagenase | Sigma-Aldrich | #C5138 |
| **Deposited data** |  |  |
| Primary data | This paper | DOI (pending) |
| **Experimental models: Organisms/strains** |  |  |
| LLPC2/3_1: w-; VT029598-p65ADZp in attp40/CyoTb; VT046081-ZpGdbd in attp2 | BDSC #88777 | SS02436 |
| LLPC1_1: w; VT048351-p65ADZ in attp40 VT057342; ZpGdbd in attP2 | BDSC #86816 | SS02437 |
| LC11: w; 22H02-p65ADZp in attp40; 20G06-ZpGdbd in attp2 | BDSC #68362 | OL0015B |
| LC22: w; 64G10-p65ADZp in attp40/Cyo; 35B06-ZpGdbd in attp2 | BDSC #68357 | OL0001B |
| LPC2: w; VT032900-p65ADZp in attp40; VT016114-ZpGDBD in attp2 | BDSC #89097 | SS25868 |
| LPLC1: w; 64G09-p65ADZp in attP40; 37H04-ZpGdbd in attP2/TM6B | BDSC #88344 | OL0029B |
| LPLC2: w; 19G02-p65ADZp in attp40; 12E04-ZpGdbd in attp2/TM6B | BDSC #88726 | OL0047B |
| LLPC1_2: w-; VT048351-p65ADZ in attp40; VT044492-ZpGdbd in attp2/TM6B | BDSC #88962 | SS02406 |
| LPT52: w; VT010661-p65ADZp in attp40/CyOTb; VT064570-ZpGDBD in attp2 | This paper | SS02019 |
| LLPC2/3_2: w-; VT046081-p65ADZp in attp40; VT029598-ZpGdbd in attp2 | BDSC #88964 | SS02422 |
| L1/L2: w-; 48A08-p65ADZp in attP40/CyO; 29G11-ZpGdbd in attP2 | BDSC #86944 | SS00797 |
| LPC1: w-; 81A05-65ADZp in attp40/CyO::Tb-RFP; VT031495-ZpGdbd in attp2/TM6B | BDSC #86818 | SS02575 |
| LPLC4: w-; 93E01-p65ADZp in attp40; VT04980-ZpGdbd in attp2 | BDSC #87849 | SS35783 |
| DNp07_1: w-; P{y[+t7.7] w[+mC]=VT029814-p65.AD}attP40/CyO, P{2xTb[1]-RFP}CyO; P{y[+t7.7] w[+mC]=VT047755-GAL4.DBD}attP2 | BDSC #75934 | SS02276 |
| DNp07_2: w-; P{y[+t7.7] w[+mC]=VT047755-p65.AD}attP40; P{y[+t7.7] w[+mC]=VT003280-GAL4.DBD}attP2 | BDSC #75973 | SS02612 |
| DNp10: w-; P{y[+t7.7] w[+mC]=R94E01-p65.AD}attP40; P{y[+t7.7] w[+mC]=VT031084-GAL4.DBD}attP2 | BDSC #75954 | SS01608 |
| DNp47: w-;; P{y[+t7.7] w[+mC]=GMR70C09-GAL4}attP2 | BDSC #39524 | SS39524 |
| DNb05: w-; P{y[+t7.7] w[+mC]=VT019391-p65.AD}attP40; P{y[+t7.7] w[+mC]=VT028198-GAL4.DBD}attP2 | BDSC #75921 | SS01051 |
| DNg79: w*; 17A04-p65ADZp in attP40; VT037534-ZpGdbd in attP2 | (Zung et al., 2025) | SS81942 |
| DNb04: w-; P{y[+t7.7] w[+mC]=VT032900-p65.AD}attP40; P{y[+t7.7] w[+mC]=VT043145-GAL4.DBD}attP2 | BDSC #75915 | SS01582 |
| Empty-split-Gal4: w-; P(y[+t7.7] w[+mC]=p65.AD.Uw)attP40; P(y[+t7.7] w[+mC]=GAL4.DBD.Uw)attP2 | BDSC #79603 | Empty |
| UAS-GFP: w[*]; P(10XUAS-IVS-GFP)attp2 | BDSC #32201 | GFP10 |
| w; VT031084(JK22C)-LexA, pJFRC 57-13XLexAOp-GFP (su(Hw)attP5); 20XUAS-CsChrimson-mCherry(su(Hw)attP1) | This study | I2B |
| w;; pJFRC49-10XUAS-IVS-eGFPKir2.1 in attP2 | (Pfeiffer et al., 2010) | Kir2.1 |
| DNb05: w-; P{y[+t7.7] w[+mC]=VT004417-p65.AD}attP40; P{y[+t7.7] w[+mC]=VT000765-GAL4.DBD}attP2 | (Zung et al., 2025) | SS94697 |
| DNb05: w-; P{y[+t7.7] w[+mC]=R73C04-p65.AD}attP40; P{y[+t7.7] w[+mC]=VT000765-GAL4.DBD}attP2 | (Zung et al., 2025) | SS94700 |
| DNb05: w-; P{y[+t7.7] w[+mC]=R73C04-p65.AD}attP40; P{y[+t7.7] w[+mC]=R39H12-GAL4.DBD}attP2 | (Namiki et al., 2018) | SS00734 |
| DNb05: w-; P{y[+t7.7] w[+mC]=VT019391-p65.AD}attP40; P{y[+t7.7] w[+mC]=VT000765-GAL4.DBD}attP2 | (Zung et al., 2025) | SS94698 |
| DNb05: w-; P{y[+t7.7] w[+mC]=VT004417-p65.AD}attP40; P{y[+t7.7] w[+mC]=R73C04-GAL4.DBD}attP2 | (Zung et al., 2025) | SS53402 |
| Oligonucleotides |  |  |
|  | This study | N/A |
| Recombinant DNA |  |  |
|  | This study | N/A |
| Software and algorithms |  |  |
| MATLAB 2020b | MathWorks, MA, USA | RRID:SCR_001622 |
| Adobe Illustrator | Adobe, San Jose, CA, USA | RRID:SCR_011323 |
| pCLAMP 11 Software Suite | Molecular Devices, LLC., CA, USA | RRID:SCR_011323 |
| RStudio | Posit, Boston, MA, USA | RRID:SCR_000432 |
| gplots | (Warnes et al., 2005) | SCR_025035 |
| iGraph for R | https://igraph.org | SCR_021238 |
| RCy3 | (Gustavsen et al., 2019) | N/A |
| Cytoscape |  | SCR_003032 |
| maleCNS R package | Jefferis G (2025). malecns: Access to the latest 'Janelia FlyEM' datasets. R package version 0.3.6, https://github.com/flyconnectome/malecns. | N/A |
| maleVNC R package | Jefferis G, Krzeminski D (2025). malevnc: Support for Working with Janelia FlyEM Male VNC Dataset. R package version 0.3.1.9100, https://natverse.org/malevnc/, https://github.com/natverse/malevnc. | N/A |
| Nat.ggplot | Bates A (2025). nat.ggplot: Functions for Creating Neuroanatomy Plots with ggplot2 and Nat. R package version 0.1.0, https://natverse.org/nat.ggplot/. |  |

**Supplemental Table 2:** Statistical reports. P-values for landing probability during DN activation, VPN silencing, and changes in wingbeat frequency during activation of DNp07 and DNp10 were calculated via Wilcoxon rank sum tests. For the DN activations, lines were tested against empty control flies. Leg angles in the last third of the analysis window and for the maximum Euclidean distance to rest within the complete analysis window were analyzed using one-way ANOVA with Tukey–Kramer post hoc correction for multiple pairwise comparisons.

| Landing probability during DN activation |  |  |  |  |  |
| --- | --- | --- | --- | --- | --- |
| Line | N | n | P-value |  |  |
| DNp07_1 | 13 | 52 | 0.000000000828055371971887 |  |  |
| DNp07_2 | 8 | 32 | 0.00000000000193530350803733 |  |  |
| DNp47 | 17 | 68 | 0.0000000711828637791926 |  |  |
| DNp10 | 8 | 32 | 0.000000683665132148772 |  |  |
| DNb05 | 13 | 52 | 0.805285972490294 |  |  |
| DNg79 | 13 | 52 | 0.000000000000000267899152442171 |  |  |
| GFP | 5 | 20 | 0.432783921121599 |  |  |
| Empty | 14 | 56 | 1 |  |  |
| Changes in wingbeat frequency during DN activation |  |  |  |  |  |
| Line | N | n | P-value t<sub>0</sub> vs t<sub>1</sub> | P-value t<sub>1</sub> vs t<sub>2</sub> |  |
| DNp07_1 | 18 | 72 | 0.0006210330514 | 0.0002079786040 |  |
| DNp10 | 11 | 44 | 0.3352275724416 | 0.5941052414022 |  |
| Empty | 9 | 36 | 0.6125051419169 | 0.6176470588235 |  |
| VPN silencing |  |  |  |  |  |
| Line | N | n | P-value, loom | P-value, bar | P-value, Approaching bar |
| LLPC1 | 14 | 56 | 0.014938603 | 0.012343316 | 0.009114974 |
| LPLC2 | 13 | 52 | 0.581882782 | 0.861985364 | 0.296593738 |
| LPLC3 | 9 | 36 | 0.012171501 | 0.005517998 | 0.004356019 |
| LC22 | 14 | 56 | 0.024742191 | 0.03738752 | 0.131827537 |
| LPT52 | 10 | 40 | 0.169290208 | 0.192666696 | 0.596929406 |

| LLPC2/3_1 | 5 | 20 | 0.08974359 | 0.019230769 | 0.047619048 |
| --- | --- | --- | --- | --- | --- |
| Leg angles, maximum Euclidian distance, front leg |  |  |  |  |  |
| Group1 | Group2 | Lower_CI | Mean_Diff | Upper_CI | p_Value |
| DNp07_1 | DNg79 | -21.1494 | -11.7122 | -2.27502 | 0.009524 |
| DNp07_1 | DNp10 | -12.8836 | -2.07196 | 8.739693 | 0.956209 |
| DNp07_1 | DNp47 | -6.53182 | 2.33287 | 11.19756 | 0.896205 |
| DNg79 | DNp10 | -1.17141 | 9.640247 | 20.4519 | 0.096072 |
| DNg79 | DNp47 | 5.180386 | 14.04508 | 22.90977 | 0.000622 |
| DNg79 | DNp47 | -5.91089 | 4.40483 | 14.72055 | 0.668618 |
| Leg angles, maximum Euclidian distance, middle leg |  |  |  |  |  |
| Group1 | Group2 | Lower_CI | Mean_Diff | Upper_CI | p_Value |
| DNp07_1 | DNg79 | 8.198754 | 20.31848 | 32.43821 | 0.000284 |
| DNp07_1 | DNp10 | -29.1572 | -15.2723 | -1.3874 | 0.025953 |
| DNp07_1 | DNp47 | -22.7228 | -11.3383 | 0.046202 | 0.051296 |
| DNg79 | DNp10 | -49.4757 | -35.5908 | -21.7059 | 9.23994650792<br>181e-08 |
| DNg79 | DNp47 | -43.0413 | -31.6568 | -20.2723 | 1.55588210093<br>427e-08 |
| DNp10 | DNp47 | -9.314 | 3.933996 | 17.18199 | 0.858218 |
| Leg angles, last third of time window, front leg |  |  |  |  |  |
| Group1 | Group2 | Lower_CI | Mean_Diff | Upper_CI | p_Value |
| DNp07_1 | DNg79 | -28.2752 | -18.3118 | -8.3484 | 6.91295724868<br>368e-05 |
| DNp07_1 | DNp10 | -10.0452 | 1.36931 | 12.78379 | 0.988572 |
| DNp07_1 | DNp47 | -2.85507 | 6.503897 | 15.86286 | 0.263066 |
| DNg79 | DNp10 | 8.266606 | 19.68109 | 31.09557 | 0.000188 |
| DNg79 | DNp47 | 15.45671 | 24.81567 | 34.17464 | 4.27988158246<br>961e-08 |
| DNp10 | DNp47 | -5.75631 | 5.134587 | 16.02549 | 0.595202 |
| Leg angles, last third of time window, middle leg |  |  |  |  |  |

| Group1 | Group2 | Lower_CI | Mean_Diff | Upper_CI | p_Value |
| --- | --- | --- | --- | --- | --- |
| DNp07_1 | DNg79 | 2.871419 | 15.40504 | 27.93866 | 0.010402 |
| DNp07_1 | DNp10 | -22.1721 | -7.81299 | 6.546071 | 0.47583 |
| DNp07_1 | DNp47 | -19.3246 | -7.55127 | 4.222013 | 0.330931 |
| DNg79 | DNp10 | -37.5771 | -23.218 | -8.85897 | 0.000473 |
| DNg79 | DNp47 | -34.7296 | -22.9563 | -11.183 | 2.53632690900<br>84e-05 |
| DNp10 | DNp47 | -13.4387 | 0.261721 | 13.96214 | 0.999952 |
| Comparisons of inter-cluster DN-DN connectivity |  |  |  |  |  |
| Group1 | Group2 | Lower_CI | Mean_Diff | Upper_CI | p_Value |
| Within cluster | Between cluster | 0.0004492463 | 0.001044054 | 0.0016388610 | 0.0007921 |

**Supplemental Table 3:** Fly genotypes used, sorted by Figures.

| Figure | Panel | Genotype |
| --- | --- | --- |
| Fig. 1 | E | DNp07_1 x CsChrimson:<br>P(y[+t7.7] w[+mC]=20XUAS-IVS-CsChrimson.mVenus)attP18/w-; P(y[+t7.7] w[+mC]=VT029814-p65.AD)attP40/+; P(y[+t7.7] w[+mC]=VT047755-GAL4.DBD)attP2)/+<br>DNp10 x CsChrimson:<br>P(y[+t7.7] w[+mC]=20XUAS-IVS-CsChrimson.mVenus)attP18/w-; P(y[+t7.7] w[+mC]=R94E01-p65.AD)attP40/+; P(y[+t7.7] w[+mC]=VT031084-GAL4.DBD)attP2)/+<br>DNp47 x CsChrimson:<br>P(y[+t7.7] w[+mC]=20XUAS-IVS-CsChrimson.mVenus)attP18/w-;;P(y[+t7.7] w[+mC]=GMR70C09-GAL4)attP2)/+<br>DNg79 x CsChrimson:<br>P(y[+t7.7] w[+mC]=20XUAS-IVS-CsChrimson.mVenus)attP18/w-; 17A04-p65ADZp in attP40/+; VT037534-ZpGdbd in attP2)/+ |
| Fig. 1 | F-G | DNp07_1 x CsChrimson:<br>P(y[+t7.7] w[+mC]=20XUAS-IVS-CsChrimson.mVenus)attP18/-; P(y[+t7.7] w[+mC]=VT029814-p65.AD)attP40/+; P(y[+t7.7] w[+mC]=VT047755-GAL4.DBD)attP2)/+<br>DNp10 x CsChrimson:<br>P(y[+t7.7] w[+mC]=20XUAS-IVS-CsChrimson.mVenus)attP18/w-; P(y[+t7.7] w[+mC]=R94E01-p65.AD)attP40/+; P(y[+t7.7] w[+mC]=VT031084-GAL4.DBD)attP2)/+<br>DNp47 x CsChrimson:<br>P(y[+t7.7] w[+mC]=20XUAS-IVS-CsChrimson.mVenus)attP18/w-;;P(y[+t7.7] w[+mC]=GMR70C09-GAL4)attP2)/+<br>DNg79 x CsChrimson:<br>P(y[+t7.7] w[+mC]=20XUAS-IVS-CsChrimson.mVenus)attP18/w-; 17A04-p65ADZp in attP40/+; VT037534-ZpGdbd in attP2)/+<br>DNb05 x CsChrimson:<br>P(y[+t7.7] w[+mC]=20XUAS-IVS-CsChrimson.mVenus)attP18/w-; P(y[+t7.7] w[+mC]=VT019391-p65.AD)attP40/+; P(y[+t7.7] w[+mC]=VT028198-GAL4.DBD)attP2)/+<br>Empty x CsChrimson:<br>P(y[+t7.7] w[+mC]=20XUAS-IVS-CsChrimson.mVenus)attP18/w-; P(y[+t7.7] w[+mC]=p65.AD.Uw)attP40/+; P(y[+t7.7] w[+mC]=GAL4.DBD.Uw)attP2)/+<br>DNp07_1 x GFP:<br>-/*; P(y[+t7.7] w[+mC]=VT029814-p65.AD)attP40/P(10XUAS-IVS-GFP)attP2)/+;P(y[+t7.7] w[+mC]=VT047755-GAL4.DBD)attP2)/+ |
| Fig. S1-2 | A | SS01598:<br>w; 42B02-p65ADZp in attP40/+; VT015782-ZpGdbd in attP2)/+<br>SS02260:<br>w; VT015782-p65ADZp in attP40/+; VT061933-ZpGdbd in attP2)/+ |
| Fig. 2 | A | DNp07_1 x CsChrimson:<br>P(y[+t7.7] w[+mC]=20XUAS-IVS-CsChrimson.mVenus)attP18/-; P(y[+t7.7] w[+mC]=VT029814-p65.AD)attP40/+; P(y[+t7.7] w[+mC]=VT047755-GAL4.DBD)attP2)/+<br>DNp10 x CsChrimson:<br>P(y[+t7.7] w[+mC]=20XUAS-IVS-CsChrimson.mVenus)attP18/w-; P(y[+t7.7] w[+mC]=R94E01-p65.AD)attP40/+; P(y[+t7.7] w[+mC]=VT031084-GAL4.DBD)attP2)/+<br>DNp47 x CsChrimson:<br>P(y[+t7.7] w[+mC]=20XUAS-IVS-CsChrimson.mVenus)attP18/w-;;P(y[+t7.7] w[+mC]=GMR70C09-GAL4)attP2)/+<br>DNg79 x CsChrimson:<br>P(y[+t7.7] w[+mC]=20XUAS-IVS-CsChrimson.mVenus)attP18/w-; 17A04-p65ADZp in attP40/+; VT037534-ZpGdbd in attP2)/+<br>DNb05 x CsChrimson:<br>P(y[+t7.7] w[+mC]=20XUAS-IVS-CsChrimson.mVenus)attP18/w-; P(y[+t7.7] w[+mC]=VT019391-p65.AD)attP40/+; P(y[+t7.7] w[+mC]=VT028198-GAL4.DBD)attP2)/+ |
| Fig. 2 | B | DNp07_1 x CsChrimson:<br>P(y[+t7.7] w[+mC]=20XUAS-IVS-CsChrimson.mVenus)attP18/-; P{y[+t7.7] w[+mC]=VT029814-p65.AD}attP40/+; P{y[+t7.7] w[+mC]=VT047755-GAL4.DBD}attP2/+<br>DNp10 x CsChrimson:<br>P(y[+t7.7] w[+mC]=20XUAS-IVS-CsChrimson.mVenus)attP18/w-; P{y[+t7.7] w[+mC]=R94E01-p65.AD}attP40/+; P{y[+t7.7] w[+mC]=VT031084-GAL4.DBD}attP2/+<br>DNp47 x CsChrimson:<br>P(y[+t7.7] w[+mC]=20XUAS-IVS-CsChrimson.mVenus)attP18/w-; P{y[+t7.7] w[+mC]=GMR70C09-GAL4}attP2/+<br>DNg79 x CsChrimson:<br>P(y[+t7.7] w[+mC]=20XUAS-IVS-CsChrimson.mVenus)attP18/w-; 17A04-p65ADZp in attP40/+; VT037534-ZpGdbd in attP2/+<br>DNb05 x CsChrimson:<br>P(y[+t7.7] w[+mC]=20XUAS-IVS-CsChrimson.mVenus)attP18/w-; P{y[+t7.7] w[+mC]=VT019391-p65.AD}attP40/+; P{y[+t7.7] w[+mC]=VT028198-GAL4.DBD}attP2/+<br>Empty x CsChrimson:<br>P(y[+t7.7] w[+mC]=20XUAS-IVS-CsChrimson.mVenus)attP18/w-; P{y[+t7.7] w[+mC]=p65.AD.Uw}attP40/+; P{y[+t7.7] w[+mC]=GAL4.DBD.Uw}attP2/+ |
| Fig 2 | G-H | DNp07_1 x CsChrimson:<br>P(y[+t7.7] w[+mC]=20XUAS-IVS-CsChrimson.mVenus)attP18/-; P{y[+t7.7] w[+mC]=VT029814-p65.AD}attP40/+; P{y[+t7.7] w[+mC]=VT047755-GAL4.DBD}attP2/+<br>DNp10 x CsChrimson:<br>P(y[+t7.7] w[+mC]=20XUAS-IVS-CsChrimson.mVenus)attP18/w-; P{y[+t7.7] w[+mC]=R94E01-p65.AD}attP40/+; P{y[+t7.7] w[+mC]=VT031084-GAL4.DBD}attP2/+<br>DNp47 x CsChrimson:<br>P(y[+t7.7] w[+mC]=20XUAS-IVS-CsChrimson.mVenus)attP18/w-; P{y[+t7.7] w[+mC]=GMR70C09-GAL4}attP2/+<br>DNg79 x CsChrimson:<br>P(y[+t7.7] w[+mC]=20XUAS-IVS-CsChrimson.mVenus)attP18/w-; 17A04-p65ADZp in attP40/+; VT037534-ZpGdbd in attP2/+ |
| Fig. 2 | J-K | DNp07_1 x CsChrimson:<br>P(y[+t7.7] w[+mC]=20XUAS-IVS-CsChrimson.mVenus)attP18/-; P{y[+t7.7] w[+mC]=VT029814-p65.AD}attP40/+; P{y[+t7.7] w[+mC]=VT047755-GAL4.DBD}attP2/+<br>DNp10 x CsChrimson:<br>P(y[+t7.7] w[+mC]=20XUAS-IVS-CsChrimson.mVenus)attP18/w-; P{y[+t7.7] w[+mC]=R94E01-p65.AD}attP40/+; P{y[+t7.7] w[+mC]=VT031084-GAL4.DBD}attP2/+<br>Empty x CsChrimson:<br>P(y[+t7.7] w[+mC]=20XUAS-IVS-CsChrimson.mVenus)attP18/w-; P{y[+t7.7] w[+mC]=p65.AD.Uw}attP40/+; P{y[+t7.7] w[+mC]=GAL4.DBD.Uw}attP2/+ |
| Fig. 2 | M | DNp07_1 x CsChrimson:<br>P(y[+t7.7] w[+mC]=20XUAS-IVS-CsChrimson.mVenus)attP18/-; P{y[+t7.7] w[+mC]=VT029814-p65.AD}attP40/+; P{y[+t7.7] w[+mC]=VT047755-GAL4.DBD}attP2/+<br>DNp07_2 x CsChrimson:<br>P(y[+t7.7] w[+mC]=20XUAS-IVS-CsChrimson.mVenus)attP18/w-; P{y[+t7.7] w[+mC]=VT047755-p65.AD}attP40/+; P{y[+t7.7] w[+mC]=VT003280-GAL4.DBD}attP2/+<br>DNp10 x CsChrimson:<br>P(y[+t7.7] w[+mC]=20XUAS-IVS-CsChrimson.mVenus)attP18/w-; P{y[+t7.7] w[+mC]=R94E01-p65.AD}attP40/+; P{y[+t7.7] w[+mC]=VT031084-GAL4.DBD}attP2/+<br>DNp47 x CsChrimson:<br>P(y[+t7.7] w[+mC]=20XUAS-IVS-CsChrimson.mVenus)attP18/w-; P{y[+t7.7] w[+mC]=GMR70C09-GAL4}attP2/+<br>DNg79 x CsChrimson:<br>P(y[+t7.7] w[+mC]=20XUAS-IVS-CsChrimson.mVenus)attP18/w-; 17A04-p65ADZp in attP40/+; VT037534-ZpGdbd in attP2/+ |
|  |  | DNb05 x CsChrimson:<br>P(y[+t7.7] w[+mC]=20XUAS-IVS-CsChrimson.mVenus)attP18/w-; P(y[+t7.7] w[+mC]=VT019391-p65.AD)attP40/+; P(y[+t7.7] w[+mC]=VT028198-GAL4.DBD)attP2/+<br>Empty x CsChrimson:<br>P(y[+t7.7] w[+mC]=20XUAS-IVS-CsChrimson.mVenus)attP18/w-; P(y[+t7.7] w[+mC]=p65.AD.Uw)attP40/+; P(y[+t7.7] w[+mC]=GAL4.DBD.Uw)attP2/+<br>LLPC2/3 1 x CsChrimson:<br>P(y[+t7.7] w[+mC]=20XUAS-IVS-CsChrimson.mVenus)attP18/w-; VT029598-p65ADZp in attP40/+; VT046081-ZpGdbd in attP2/+<br>DNp07_1 x GFP:<br>-/*; P(y[+t7.7] w[+mC]=VT029814-p65.AD)attP40/P(10XUAS-IVS-GFP)attP2/+; P(y[+t7.7] w[+mC]=VT047755-GAL4.DBD)attP2)/+ |
| Fig. S2-1 | A | DNp07_2 x CsChrimson:<br>P(y[+t7.7] w[+mC]=20XUAS-IVS-CsChrimson.mVenus)attP18/w-; P(y[+t7.7] w[+mC]=VT047755-p65.AD)attP40/+; P(y[+t7.7] w[+mC]=VT003280-GAL4.DBD)attP2/+<br>Empty x CsChrimson:<br>P(y[+t7.7] w[+mC]=20XUAS-IVS-CsChrimson.mVenus)attP18/w-; P(y[+t7.7] w[+mC]=p65.AD.Uw)attP40/+; P(y[+t7.7] w[+mC]=GAL4.DBD.Uw)attP2/+<br>LLPC2/3 1 x CsChrimson:<br>P(y[+t7.7] w[+mC]=20XUAS-IVS-CsChrimson.mVenus)attP18/w-; VT029598-p65ADZp in attP40/+; VT046081-ZpGdbd in attP2/+ |
| Fig. S2-1 | B | DNp07_1 x CsChrimson:<br>P(y[+t7.7] w[+mC]=20XUAS-IVS-CsChrimson.mVenus)attP18/-; P(y[+t7.7] w[+mC]=VT029814-p65.AD)attP40/+; P(y[+t7.7] w[+mC]=VT047755-GAL4.DBD)attP2)/+<br>DNp07_2 x CsChrimson:<br>P(y[+t7.7] w[+mC]=20XUAS-IVS-CsChrimson.mVenus)attP18/w-; P(y[+t7.7] w[+mC]=VT047755-p65.AD)attP40/+; P(y[+t7.7] w[+mC]=VT003280-GAL4.DBD)attP2/+<br>DNp10 x CsChrimson:<br>P(y[+t7.7] w[+mC]=20XUAS-IVS-CsChrimson.mVenus)attP18/w-; P(y[+t7.7] w[+mC]=R94E01-p65.AD)attP40/+; P(y[+t7.7] w[+mC]=VT031084-GAL4.DBD)attP2/+<br>DNp47 x CsChrimson:<br>P(y[+t7.7] w[+mC]=20XUAS-IVS-CsChrimson.mVenus)attP18/w-; P(y[+t7.7] w[+mC]=GMR70C09-GAL4)attP2/+<br>DNp79 x CsChrimson:<br>P(y[+t7.7] w[+mC]=20XUAS-IVS-CsChrimson.mVenus)attP18/w-; 17A04-p65ADZp in attP40/+; VT037534-ZpGdbd in attP2/+<br>DNb05 x CsChrimson:<br>P(y[+t7.7] w[+mC]=20XUAS-IVS-CsChrimson.mVenus)attP18/w-; P(y[+t7.7] w[+mC]=VT019391-p65.AD)attP40/+; P(y[+t7.7] w[+mC]=VT028198-GAL4.DBD)attP2/+<br>Empty x CsChrimson:<br>P(y[+t7.7] w[+mC]=20XUAS-IVS-CsChrimson.mVenus)attP18/w-; P(y[+t7.7] w[+mC]=p65.AD.Uw)attP40/+; P(y[+t7.7] w[+mC]=GAL4.DBD.Uw)attP2/+<br>LLPC2/3 1 x CsChrimson:<br>P(y[+t7.7] w[+mC]=20XUAS-IVS-CsChrimson.mVenus)attP18/w-; VT029598-p65ADZp in attP40/+; VT046081-ZpGdbd in attP2/+<br>DNp07_1 x GFP:<br>-/*; P(y[+t7.7] w[+mC]=VT029814-p65.AD)attP40/P(10XUAS-IVS-GFP)attP2/+; P(y[+t7.7] w[+mC]=VT047755-GAL4.DBD)attP2)/+ |
| Fig. S2-1 | C | DNp07_2 x CsChrimson:<br>P(y[+t7.7] w[+mC]=20XUAS-IVS-CsChrimson.mVenus)attP18/w-; P(y[+t7.7] w[+mC]=VT047755-p65.AD)attP40/+; P(y[+t7.7] w[+mC]=VT003280-GAL4.DBD)attP2/+<br>Empty x CsChrimson:<br>P(y[+t7.7] w[+mC]=20XUAS-IVS-CsChrimson.mVenus)attP18/w-; P(y[+t7.7] w[+mC]=p65.AD.Uw)attP40/+; P(y[+t7.7] w[+mC]=GAL4.DBD.Uw)attP2/+<br>LLPC2/3 1 x CsChrimson:<br>P(y[+t7.7] w[+mC]=20XUAS-IVS-CsChrimson.mVenus)attP18/w-; VT029598-p65ADZp in attP40/+; VT046081-ZpGdbd in attP2/+ |
| Fig. S2-1 | G-H | DNp07_1 x CsChrimson:<br>P(y[+t7.7] w[+mC]=20XUAS-IVS-CsChrimson.mVenus)attP18/-; P{y[+t7.7] w[+mC]=VT029814-p65.AD}attP40/+; P{y[+t7.7] w[+mC]=VT047755-GAL4.DBD}attP2)/+<br>DNp07_2 x CsChrimson:<br>P(y[+t7.7] w[+mC]=20XUAS-IVS-CsChrimson.mVenus)attP18/w-; P{y[+t7.7] w[+mC]=VT047755-p65.AD}attP40/+; P{y[+t7.7] w[+mC]=VT003280-GAL4.DBD}attP2)/+<br>LLPC2/3 1 x CsChrimson:<br>P(y[+t7.7] w[+mC]=20XUAS-IVS-CsChrimson.mVenus)attP18/w-; VT029598-p65ADZp in attP40/+; VT046081-ZpGdbd in attP2)/+ |
| Fig. S2-1 | I | DNg79 x CsChrimson:<br>P(y[+t7.7] w[+mC]=20XUAS-IVS-CsChrimson.mVenus)attP18/w-; 17A04-p65ADZp in attP40/+; VT037534-ZpGdbd in attP2)/+ |
| Fig. S2-2 | C-G | DNp07_1 x CsChrimson:<br>P(y[+t7.7] w[+mC]=20XUAS-IVS-CsChrimson.mVenus)attP18/-; P{y[+t7.7] w[+mC]=VT029814-p65.AD}attP40/+; P{y[+t7.7] w[+mC]=VT047755-GAL4.DBD}attP2)/+<br>DNp10 x CsChrimson:<br>P(y[+t7.7] w[+mC]=20XUAS-IVS-CsChrimson.mVenus)attP18/w-; P{y[+t7.7] w[+mC]=R94E01-p65.AD}attP40/+; P{y[+t7.7] w[+mC]=VT031084-GAL4.DBD}attP2)/+<br>Empty x CsChrimson:<br>P(y[+t7.7] w[+mC]=20XUAS-IVS-CsChrimson.mVenus)attP18/w-; P{y[+t7.7] w[+mC]=p65.AD.Uw}attP40/+; P{y[+t7.7] w[+mC]=GAL4.DBD.Uw}attP2)/+ |
| Fig. S2-2 | I-J | DNp07_1 x CsChrimson:<br>P(y[+t7.7] w[+mC]=20XUAS-IVS-CsChrimson.mVenus)attP18/-; P{y[+t7.7] w[+mC]=VT029814-p65.AD}attP40/+; P{y[+t7.7] w[+mC]=VT047755-GAL4.DBD}attP2)/+<br>DNp07_2 x CsChrimson:<br>P(y[+t7.7] w[+mC]=20XUAS-IVS-CsChrimson.mVenus)attP18/w-; P{y[+t7.7] w[+mC]=VT047755-p65.AD}attP40/+; P{y[+t7.7] w[+mC]=VT003280-GAL4.DBD}attP2)/+<br>DNp10 x CsChrimson:<br>P(y[+t7.7] w[+mC]=20XUAS-IVS-CsChrimson.mVenus)attP18/w-; P{y[+t7.7] w[+mC]=R94E01-p65.AD}attP40/+; P{y[+t7.7] w[+mC]=VT031084-GAL4.DBD}attP2)/+<br>DNp47 x CsChrimson:<br>P(y[+t7.7] w[+mC]=20XUAS-IVS-CsChrimson.mVenus)attP18/w-; P{y[+t7.7] w[+mC]=GMR70C09-GAL4}attP2)/+<br>DNg79 x CsChrimson:<br>P(y[+t7.7] w[+mC]=20XUAS-IVS-CsChrimson.mVenus)attP18/w-; 17A04-p65ADZp in attP40/+; VT037534-ZpGdbd in attP2)/+<br>DNb05 x CsChrimson:<br>P(y[+t7.7] w[+mC]=20XUAS-IVS-CsChrimson.mVenus)attP18/w-; P{y[+t7.7] w[+mC]=VT019391-p65.AD}attP40/+; P{y[+t7.7] w[+mC]=VT028198-GAL4.DBD}attP2)/+<br>Empty x CsChrimson:<br>P(y[+t7.7] w[+mC]=20XUAS-IVS-CsChrimson.mVenus)attP18/w-; P{y[+t7.7] w[+mC]=p65.AD.Uw}attP40/+; P{y[+t7.7] w[+mC]=GAL4.DBD.Uw}attP2)/+<br>LLPC2/3 1 x CsChrimson:<br>P(y[+t7.7] w[+mC]=20XUAS-IVS-CsChrimson.mVenus)attP18/w-; VT029598-p65ADZp in attP40/+; VT046081-ZpGdbd in attP2)/+<br>DNp07_1 x GFP:<br>-/*; P{y[+t7.7] w[+mC]=VT029814-p65.AD}attP40/P(10XUAS-IVS-GFP)attP2)/+; P{y[+t7.7] w[+mC]=VT047755-GAL4.DBD}attP2)/+ |
| Fig. 3 | D | DNp10: w; P{y[+t7.7] w[+mC]=R94E01-p65.AD}attP40/+; P{y[+t7.7] w[+mC]=VT031084-GAL4.DBD}attP2/5XUAS-IVS-myr::smFLAG in VK00005, pJFRC51-3XUAS-IVS-Syt::smHA in su(Hw)attP1<br>OL0029B:<br>w; 64G09-p65ADZp in attP40/+; 37H04-ZpGdbd in attP2/5XUAS-IVS-myr::smFLAG in VK00005, pJFRC51-3XUAS-IVS-Syt::smHA in su(Hw)attP1<br>OL0047B: |
|  |  | w; 19G02-p65ADZp in attP40/+; 12E04-ZpGdbd in attP2/5XUAS-IVS-myr::smFLAG in VK00005, pJFRC51-3XUAS-IVS-Syt::smHA in su(Hw)attP1 SS35783:<br>w-; 93E01-p65ADZp in attP40/+; VT04980-ZpGdbd in attP2/5XUAS-IVS-myr::smFLAG in VK00005, pJFRC51-3XUAS-IVS-Syt::smHA in su(Hw)attP1 SS01608:<br>w; VT010661-p65ADZp in attP40/+; VT064570-ZpGDBD in attP2/5XUAS-IVS-myr::smFLAG in VK00005, pJFRC51-3XUAS-IVS-Syt::smHA in su(Hw)attP1 |
| Fig. 3 | H-I | LLPC2/3 1 x CsChrimson:<br>P(y[+t7.7] w[+mC]=20XUAS-IVS-CsChrimson.mVenus)attP18/w-; VT029598-p65ADZp in attP40/+; VT046081-ZpGdbd in attP2/+<br>LLPC1_1 x CsChrimson:<br>P(y[+t7.7] w[+mC]=20XUAS-IVS-CsChrimson.mVenus)attP18/w-; VT048351-p65ADZ in attP40/+; VT057342; ZpGdbd in attP2/+<br>LC11 x CsChrimson:<br>P(y[+t7.7] w[+mC]=20XUAS-IVS-CsChrimson.mVenus)attP18/w; 22H02-p65ADZp in attP40/+; 20G06-ZpGdbd in attP2/+<br>LC22 x CsChrimson:<br>P(y[+t7.7] w[+mC]=20XUAS-IVS-CsChrimson.mVenus)attP18/w; 64G10-p65ADZp in attP40/+; 35B06-ZpGdbd in attP2/+<br>LPC2 x CsChrimson:<br>P(y[+t7.7] w[+mC]=20XUAS-IVS-CsChrimson.mVenus)attP18/w; VT032900-p65ADZp in attP40/+; VT016114-ZpGDBD in attP2/+<br>LPLC1 x CsChrimson:<br>P(y[+t7.7] w[+mC]=20XUAS-IVS-CsChrimson.mVenus)attP18/w; 64G09-p65ADZp in attP40/+; 37H04-ZpGdbd in attP2/+<br>LPLC2 x CsChrimson:<br>P(y[+t7.7] w[+mC]=20XUAS-IVS-CsChrimson.mVenus)attP18/w; 19G02-p65ADZp in attP40/+; 12E04-ZpGdbd in attP2/+<br>LPT52 x CsChrimson:<br>P(y[+t7.7] w[+mC]=20XUAS-IVS-CsChrimson.mVenus)attP18/w; VT010661-p65ADZp in attP40/+; VT064570-ZpGDBD in attP2/+<br>LPC1 x CsChrimson:<br>P(y[+t7.7] w[+mC]=20XUAS-IVS-CsChrimson.mVenus)attP18/w-; 81A05-65ADZp in attP40/+; VT031495-ZpGdbd in attP2/+<br>LPLC4 x CsChrimson:<br>P(y[+t7.7] w[+mC]=20XUAS-IVS-CsChrimson.mVenus)attP18/w-; 93E01-p65ADZp in attP40/+; VT04980-ZpGdbd in attP2/+ |
| Fig. 3 | J | LLPC2/3 1 x Kir2.1:<br>w-; VT029598-p65ADZp in attP40/+; VT046081-ZpGdbd in attP2/pJFRC49-10XUAS-IVS-eGFPKir2.1 (attP2)<br>LLPC1_1 x Kir2.1:<br>w-; VT048351-p65ADZ in attP40/+; VT057342; ZpGdbd in attP2/pJFRC49-10XUAS-IVS-eGFPKir2.1 (attP2)<br>LC22 x Kir 2.1:<br>w; 64G10-p65ADZp in attP40/+; 35B06-ZpGdbd in attP2/pJFRC49-10XUAS-IVS-eGFPKir2.1 (attP2)<br>LPLC2 x Kir2.1:<br>w; 19G02-p65ADZp in attP40/+; 12E04-ZpGdbd in attP2/pJFRC49-10XUAS-IVS-eGFPKir2.1 (attP2)<br>LPT52 x Kir2.1:<br>w; VT010661-p65ADZp in attP40/+; VT064570-ZpGDBD in attP2/pJFRC49-10XUAS-IVS-eGFPKir2.1 (attP2)<br>L1/L2 x Kir2.1:<br>w-; 48A08-p65ADZp in attP40/+; 29G11-ZpGdbd in attP2/pJFRC49-10XUAS-IVS-eGFPKir2.1 (attP2)<br>LPLC4 x Kir2.1:<br>w-; 93E01-p65ADZp in attP40/+; VT04980-ZpGdbd in attP2/pJFRC49-10XUAS-IVS-eGFPKir2.1 (attP2)<br>DNp10 x GFP10:<br>w-/w*; P(y[+t7.7] w[+mC]=VT029814-p65.AD)attP40/+; P(10XUAS-IVS-GFP)attP2/P(y[+t7.7] w[+mC]=VT047755-GAL4.DBD)attP2<br>Empty x Kir2.1:<br>w-/w; P(y[+t7.7] w[+mC]=p65.AD.Uw)attP40/+; P(y[+t7.7] w[+mC]=GAL4.DBD.Uw)attP2/ pJFRC49-10XUAS-IVS-eGFPKir2.1 in attP2 |
| Fig. 3 | K-I | I2B x LLPC2/3 1:<br>w-/w; VT029598-p65ADZp in attp40/VT031084(JK22C)-LexA, pJFRC 57-13XLexAOp-GFP (su(Hw)attP5); VT046081-ZpGdbd in attp2/20XUAS-CsChrimson-mCherry(su(Hw)attP1)<br>I2B x LC22:<br>w; 64G10-p65ADZp in attp40/VT031084(JK22C)-LexA, pJFRC 57-13XLexAOp-GFP (su(Hw)attP5); 35B06-ZpGdbd in attp2/20XUAS-CsChrimson-mCherry(su(Hw)attP1)<br>Empty x CsChrimson:<br>P(y[+t7.7] w[+mC]=20XUAS-IVS-CsChrimson.mVenus)attP18/w-; P(y[+t7.7] w[+mC]=p65.AD.Uw)attP40/+; P(y[+t7.7] w[+mC]=GAL4.DBD.Uw)attP2/+ |
| Fig. S3 | B-E | DNp07_1 x CsChrimson:<br>P(y[+t7.7] w[+mC]=20XUAS-IVS-CsChrimson.mVenus)attP18/-; P(y[+t7.7] w[+mC]=VT029814-p65.AD)attP40/+; P(y[+t7.7] w[+mC]=VT047755-GAL4.DBD)attP2)/+<br>DNp10 x CsChrimson:<br>P(y[+t7.7] w[+mC]=20XUAS-IVS-CsChrimson.mVenus)attP18/w-; P(y[+t7.7] w[+mC]=R94E01-p65.AD)attP40/+; P(y[+t7.7] w[+mC]=VT031084-GAL4.DBD)attP2)/+<br>Empty x CsChrimson:<br>P(y[+t7.7] w[+mC]=20XUAS-IVS-CsChrimson.mVenus)attP18/w-; P(y[+t7.7] w[+mC]=p65.AD.Uw)attP40/+; P(y[+t7.7] w[+mC]=GAL4.DBD.Uw)attP2)/+ |
| Fig. S3-1 | A-B | w; VT010661-p65ADZp in attp40/+; VT064570-ZpGDBD in attp2/5XUAS-IVS-myr::smFLAG in VK00005, pJFRC51-3XUAS-IVS-Syt::smHA in su(Hw)attP1 |
| Fig. S3-1 | C | OL0029B:<br>w; 64G09-p65ADZp in attP40/+; 37H04-ZpGdbd in attP2/5XUAS-IVS-myr::smFLAG in VK00005, pJFRC51-3XUAS-IVS-Syt::smHA in su(Hw)attP1<br>OL0047B:<br>w; 19G02-p65ADZp in attp40/+; 12E04-ZpGdbd in attp2/5XUAS-IVS-myr::smFLAG in VK00005, pJFRC51-3XUAS-IVS-Syt::smHA in su(Hw)attP1<br>SS35783:<br>w-; 93E01-p65ADZp in attp40/+; VT04980-ZpGdbd in attp2/5XUAS-IVS-myr::smFLAG in VK00005, pJFRC51-3XUAS-IVS-Syt::smHA in su(Hw)attP1<br>OL0001B:<br>w; 64G10-p65ADZp in attp40/+; 35B06-ZpGdbd in attp2/5XUAS-IVS-myr::smFLAG in VK00005, pJFRC51-3XUAS-IVS-Syt::smHA in su(Hw)attP1<br>SS02575:<br>w-; 81A05-p65ADZp in attp40/+; VT031495-ZpGdbd in attp2/5XUAS-IVS-myr::smFLAG in VK00005, pJFRC51-3XUAS-IVS-Syt::smHA in su(Hw)attP1<br>SS25868:<br>w; VT032900-p65ADZp in attp40/+; VT016114-ZpGDBD in attp2/5XUAS-IVS-myr::smFLAG in VK00005, pJFRC51-3XUAS-IVS-Syt::smHA in su(Hw)attP1<br>SS02437:<br>w-; VT048351-p65ADZ in attp40/+; VT057342; ZpGdbd in attP2/5XUAS-IVS-myr::smFLAG in VK00005, pJFRC51-3XUAS-IVS-Syt::smHA in su(Hw)attP1<br>SS02406:<br>w-; VT048351-p65ADZ in attp40/+; VT044492-ZpGdbd in attp2/5XUAS-IVS-myr::smFLAG in VK00005, pJFRC51-3XUAS-IVS-Syt::smHA in su(Hw)attP1<br>SS02436:<br>w-; VT029598-p65ADZp in attp40/+; VT046081-ZpGdbd in attp2/5XUAS-IVS-myr::smFLAG in VK00005, pJFRC51-3XUAS-IVS-Syt::smHA in su(Hw)attP1<br>SS02422:<br>w-; VT046081-p65ADZp in attp40/+; VT029598-ZpGdbd in attp2/5XUAS-IVS-myr::smFLAG in VK00005, pJFRC51-3XUAS-IVS-Syt::smHA in su(Hw)attP1<br>SS00797:<br>w-; 48A08-p65ADZp in attP40/+; 29G11-ZpGdbd in attP2/5XUAS-IVS-myr::smFLAG in VK00005, pJFRC51-3XUAS-IVS-Syt::smHA in su(Hw)attP1<br>OL0015B:<br>w; 22H02-p65ADZp in attp40/+; 20G06-ZpGdbd in attp2/5XUAS-IVS-myr::smFLAG in VK00005, pJFRC51-3XUAS-IVS-Syt::smHA in su(Hw)attP1 |
| Fig. S3-2 | A | LLPC2/3 1 x CsChrimson:<br>P(y[+t7.7] w[+mC]=20XUAS-IVS-CsChrimson.mVenus)attP18/w-; VT029598-p65ADZp in attp40/+; VT046081-ZpGdbd in attp2/+ |
| Fig. S3-2 | B-C | LLPC2/3 2 x CsChrimson:<br>P(y[+t7.7] w[+mC]=20XUAS-IVS-CsChrimson.mVenus)attP18/w-; VT046081-p65ADZp in attp40/+; VT029598-ZpGdbd in attp2/+<br>LLPC1_2 x CsChrimson:<br>P(y[+t7.7] w[+mC]=20XUAS-IVS-CsChrimson.mVenus)attP18/w-; VT048351-p65ADZ in attp40/+; VT044492-ZpGdbd in attp2/+<br>L1/L2 x CsChrimson:<br>P(y[+t7.7] w[+mC]=20XUAS-IVS-CsChrimson.mVenus)attP18/w-; 48A08-p65ADZp in attp40/+; 29G11-ZpGdbd in attp2/+ |
| Fig. S3-2 | D | LLPC2/3 1 x CsChrimson:<br>P(y[+t7.7] w[+mC]=20XUAS-IVS-CsChrimson.mVenus)attP18/w-; VT029598-p65ADZp in attp40/+; VT046081-ZpGdbd in attp2/+ |
| Fig. S3-2 | E | LLPC2/3 1 x CsChrimson:<br>P(y[+t7.7] w[+mC]=20XUAS-IVS-CsChrimson.mVenus)attP18/w-; VT029598-p65ADZp in attp40/+; VT046081-ZpGdbd in attp2/+<br>LLPC1_1 x CsChrimson:<br>P(y[+t7.7] w[+mC]=20XUAS-IVS-CsChrimson.mVenus)attP18/w-; VT048351-p65ADZ in attp40/+; VT057342; ZpGdbd in attp2/+<br>LPLC1 x CsChrimson:<br>P(y[+t7.7] w[+mC]=20XUAS-IVS-CsChrimson.mVenus)attP18/w; 64G09-p65ADZp in attp40/+; 37H04-ZpGdbd in attp2/+<br>LPLC2 x CsChrimson:<br>P(y[+t7.7] w[+mC]=20XUAS-IVS-CsChrimson.mVenus)attP18/w; 19G02-p65ADZp in attp40/+; 12E04-ZpGdbd in attp2/+<br>LPT52 x CsChrimson:<br>P(y[+t7.7] w[+mC]=20XUAS-IVS-CsChrimson.mVenus)attP18/w; VT010661-p65ADZp in attp40/+; VT064570-ZpGDBD in attp2/+<br>LPLC4 x CsChrimson:<br>P(y[+t7.7] w[+mC]=20XUAS-IVS-CsChrimson.mVenus)attP18/w-; 93E01-p65ADZp in attp40/+; VT04980-ZpGdbd in attp2/+ |
| Fig. S3-2 | F | LLPC2/3 1 x CsChrimson:<br>P(y[+t7.7] w[+mC]=20XUAS-IVS-CsChrimson.mVenus)attP18/w-; VT029598-p65ADZp in attp40/+; VT046081-ZpGdbd in attp2/+<br>LLPC1_1 x CsChrimson:<br>P(y[+t7.7] w[+mC]=20XUAS-IVS-CsChrimson.mVenus)attP18/w-; VT048351-p65ADZ in attp40/+; VT057342; ZpGdbd in attp2/+<br>LC11 x CsChrimson:<br>P(y[+t7.7] w[+mC]=20XUAS-IVS-CsChrimson.mVenus)attP18/w; 22H02-p65ADZp in attp40/+; 20G06-ZpGdbd in attp2/+<br>LC22 x CsChrimson:<br>P(y[+t7.7] w[+mC]=20XUAS-IVS-CsChrimson.mVenus)attP18/w; 64G10-p65ADZp in attp40/+; 35B06-ZpGdbd in attp2/+<br>LPC2 x CsChrimson:<br>P(y[+t7.7] w[+mC]=20XUAS-IVS-CsChrimson.mVenus)attP18/w; VT032900-p65ADZp in attp40/+; VT016114-ZpGDBD in attp2/+<br>LPLC1 x CsChrimson:<br>P(y[+t7.7] w[+mC]=20XUAS-IVS-CsChrimson.mVenus)attP18/w; 64G09-p65ADZp in attp40/+; 37H04-ZpGdbd in attp2/+<br>LPLC2 x CsChrimson:<br>P(y[+t7.7] w[+mC]=20XUAS-IVS-CsChrimson.mVenus)attP18/w; 19G02-p65ADZp in attp40/+; 12E04-ZpGdbd in attp2/+<br>LPT52 x CsChrimson:<br>P(y[+t7.7] w[+mC]=20XUAS-IVS-CsChrimson.mVenus)attP18/w; VT010661-p65ADZp in attp40/+; VT064570-ZpGDBD in attp2/+<br>LPC1 x CsChrimson:<br>P(y[+t7.7] w[+mC]=20XUAS-IVS-CsChrimson.mVenus)attP18/w-; 81A05-p65ADZp in attp40/+; VT031495-ZpGdbd in attp2/+<br>LPLC4 x CsChrimson:<br>P(y[+t7.7] w[+mC]=20XUAS-IVS-CsChrimson.mVenus)attP18/w-; 93E01-p65ADZp in attp40/+; VT04980-ZpGdbd in attp2/+ |
|  |  | LLPC2/3 2 x CsChrimson:<br>P(y[+t7.7] w[+mC]=20XUAS-IVS-CsChrimson.mVenus)attP18/w-; VT046081-p65ADZp in attP40/+; VT029598-ZpGdbd in attP2/+<br>LLPC1_2 x CsChrimson:<br>P(y[+t7.7] w[+mC]=20XUAS-IVS-CsChrimson.mVenus)attP18/w-; VT048351-p65ADZ in attP40/+; VT044492-ZpGdbd in attP2/+<br>L1/L2 x CsChrimson:<br>P(y[+t7.7] w[+mC]=20XUAS-IVS-CsChrimson.mVenus)attP18/w-; 48A08-p65ADZp in attP40/+; 29G11-ZpGdbd in attP2/+ |
| Fig. S3-2 | G | LPLC4 x CsChrimson:<br>P(y[+t7.7] w[+mC]=20XUAS-IVS-CsChrimson.mVenus)attP18/w-; 93E01-p65ADZp in attP40/+; VT04980-ZpGdbd in attP2/+<br>LLPC2/3 1 x CsChrimson:<br>P(y[+t7.7] w[+mC]=20XUAS-IVS-CsChrimson.mVenus)attP18/w-; VT029598-p65ADZp in attP40/+; VT046081-ZpGdbd in attP2/+<br>LLPC1_1 x CsChrimson:<br>P(y[+t7.7] w[+mC]=20XUAS-IVS-CsChrimson.mVenus)attP18/w-; VT048351-p65ADZ in attP40/+; VT057342; ZpGdbd in attP2/+<br>LPLC1 x CsChrimson:<br>P(y[+t7.7] w[+mC]=20XUAS-IVS-CsChrimson.mVenus)attP18/w; 64G09-p65ADZp in attP40/+; 37H04-ZpGdbd in attP2/+ |
| Fig. S3-2 | H | LLPC2/3 1 x CsChrimson:<br>P(y[+t7.7] w[+mC]=20XUAS-IVS-CsChrimson.mVenus)attP18/w-; VT029598-p65ADZp in attP40/+; VT046081-ZpGdbd in attP2/+<br>LLPC2/3 2 x CsChrimson:<br>P(y[+t7.7] w[+mC]=20XUAS-IVS-CsChrimson.mVenus)attP18/w-; VT046081-p65ADZp in attP40/+; VT029598-ZpGdbd in attP2/+<br>LLPC1_1 x CsChrimson:<br>P(y[+t7.7] w[+mC]=20XUAS-IVS-CsChrimson.mVenus)attP18/w-; VT048351-p65ADZ in attP40/+; VT057342; ZpGdbd in attP2/+<br>LLPC1_2 x CsChrimson:<br>P(y[+t7.7] w[+mC]=20XUAS-IVS-CsChrimson.mVenus)attP18/w-; VT048351-p65ADZ in attP40/+; VT044492-ZpGdbd in attP2/+<br>LPLC4 x CsChrimson:<br>P(y[+t7.7] w[+mC]=20XUAS-IVS-CsChrimson.mVenus)attP18/w-; 93E01-p65ADZp in attP40/+; VT04980-ZpGdbd in attP2/+<br>LPLC2 x CsChrimson:<br>P(y[+t7.7] w[+mC]=20XUAS-IVS-CsChrimson.mVenus)attP18/w; 19G02-p65ADZp in attP40/+; 12E04-ZpGdbd in attP2/+<br>LPLC1 x CsChrimson:<br>P(y[+t7.7] w[+mC]=20XUAS-IVS-CsChrimson.mVenus)attP18/w; 64G09-p65ADZp in attP40/+; 37H04-ZpGdbd in attP2/+<br>LPT52 x CsChrimson:<br>P(y[+t7.7] w[+mC]=20XUAS-IVS-CsChrimson.mVenus)attP18/w; VT010661-p65ADZp in attP40/+; VT064570-ZpGDBD in attP2/+<br>LPC2 x CsChrimson:<br>P(y[+t7.7] w[+mC]=20XUAS-IVS-CsChrimson.mVenus)attP18/w; VT032900-p65ADZp in attP40/+; VT016114-ZpGDBD in attP2/+<br>LPC1 x CsChrimson:<br>P(y[+t7.7] w[+mC]=20XUAS-IVS-CsChrimson.mVenus)attP18/w-; 81A05-p65ADZp in attP40/+; VT031495-ZpGdbd in attP2/+ |
| Fig. S3-2 | I-J | LLPC2/3 1 x Kir2.1:<br>w-; VT029598-p65ADZp in attP40/+; VT046081-ZpGdbd in attP2/pJFRC49-10XUAS-IVS-eGFPKir2.1 (attP2)<br>LLPC1_1 x Kir2.1:<br>w-; VT048351-p65ADZ in attP40/+; VT057342; ZpGdbd in attP2/pJFRC49-10XUAS-IVS-eGFPKir2.1 (attP2)<br>LC22 x Kir 2.1:<br>w; 64G10-p65ADZp in attP40/+; 35B06-ZpGdbd in attP2/pJFRC49-10XUAS-IVS-eGFPKir2.1 (attP2)<br>LPLC2 x Kir2.1:<br>w; 19G02-p65ADZp in attP40/+; 12E04-ZpGdbd in attP2/pJFRC49-10XUAS-IVS-eGFPKir2.1 (attP2) |
|  |  | LPT52 x Kir2.1:<br>w; VT010661-p65ADZp in attP40/+; VT064570-ZpGDBD in attP2/pJFRC49-10XUAS-IVS-eGFPKir2.1 (attP2)<br>L1/L2 x Kir2.1:<br>w-; 48A08-p65ADZp in attP40/+; 29G11-ZpGdbd in attP2/pJFRC49-10XUAS-IVS-eGFPKir2.1 (attP2)<br>LPLC4 x Kir2.1:<br>w-; 93E01-p65ADZp in attP40/+; VT04980-ZpGdbd in attP2/pJFRC49-10XUAS-IVS-eGFPKir2.1 (attP2)<br>DNp10 x GFP10:<br>w-/w*; P{y[+t7.7] w[+mC]=VT029814-p65.AD}attP40/+; P(10XUAS-IVS-GFP)attP2/P{y[+t7.7] w[+mC]=VT047755-GAL4.DBD}attP2<br>Empty x Kir2.1:<br>w-/w; P{y[+t7.7] w[+mC]=p65.AD.Uw}attP40/+; P{y[+t7.7] w[+mC]=GAL4.DBD.Uw}attP2/ pJFRC49-10XUAS-IVS-eGFPKir2.1 in attP2 |
| Fig. S3-2 | K | I2B x LLPC2/3 1:<br>w-/w; VT029598-p65ADZp in attP40/VT031084(JK22C)-LexA, pJFRC 57-13XLexAOp-GFP (su(Hw)attP5); VT046081-ZpGdbd in attP2/20XUAS-CsChrimson-mCherry(su(Hw)attP1)<br>I2B x LC22:<br>w; 64G10-p65ADZp in attP40/VT031084(JK22C)-LexA, pJFRC 57-13XLexAOp-GFP (su(Hw)attP5); 35B06-ZpGdbd in attP2/20XUAS-CsChrimson-mCherry(su(Hw)attP1)<br>Empty x CsChrimson:<br>P{y[+t7.7] w[+mC]=20XUAS-IVS-CsChrimson.mVenus}attP18/w-; P{y[+t7.7] w[+mC]=p65.AD.Uw}attP40/+; P{y[+t7.7] w[+mC]=GAL4.DBD.Uw}attP2/+ |
| Fig. 4 | C-D | DNp07_1 x CsChrimson:<br>P{y[+t7.7] w[+mC]=20XUAS-IVS-CsChrimson.mVenus}attP18/-; P{y[+t7.7] w[+mC]=VT029814-p65.AD}attP40/+; P{y[+t7.7] w[+mC]=VT047755-GAL4.DBD}attP2/+<br>DNp10 x CsChrimson:<br>P{y[+t7.7] w[+mC]=20XUAS-IVS-CsChrimson.mVenus}attP18/w-; P{y[+t7.7] w[+mC]=R94E01-p65.AD}attP40/+; P{y[+t7.7] w[+mC]=VT031084-GAL4.DBD}attP2/+<br>DNp47 x CsChrimson:<br>P{y[+t7.7] w[+mC]=20XUAS-IVS-CsChrimson.mVenus}attP18/w-; P{y[+t7.7] w[+mC]=GMR70C09-GAL4}attP2/+<br>DNp79 x CsChrimson:<br>P{y[+t7.7] w[+mC]=20XUAS-IVS-CsChrimson.mVenus}attP18/w-; 17A04-p65ADZp in attP40/+; VT037534-ZpGdbd in attP2/+<br>DNb05 x CsChrimson:<br>P{y[+t7.7] w[+mC]=20XUAS-IVS-CsChrimson.mVenus}attP18/w-; P{y[+t7.7] w[+mC]=VT019391-p65.AD}attP40/+; P{y[+t7.7] w[+mC]=VT028198-GAL4.DBD}attP2/+<br>Empty x CsChrimson:<br>P{y[+t7.7] w[+mC]=20XUAS-IVS-CsChrimson.mVenus}attP18/w-; P{y[+t7.7] w[+mC]=p65.AD.Uw}attP40/+; P{y[+t7.7] w[+mC]=GAL4.DBD.Uw}attP2/+<br>GFP x CsChrimson:<br>P{y[+t7.7] w[+mC]=20XUAS-IVS-CsChrimson.mVenus}attP18/*; P(10XUAS-IVS-GFP)attP2/+ |
| Fig. 4 | F-K | DNp10 x GFP10:<br>w-/w*; P{y[+t7.7] w[+mC]=VT029814-p65.AD}attP40/P(10XUAS-IVS-GFP)attP2;<br>P{y[+t7.7] w[+mC]=VT047755-GAL4.DBD}attP2/+ |
| Fig. S4-1 | D-E | DNp10 x GFP10:<br>w-/w*; P{y[+t7.7] w[+mC]=VT029814-p65.AD}attP40/P(10XUAS-IVS-GFP)attP2;<br>P{y[+t7.7] w[+mC]=VT047755-GAL4.DBD}attP2/+ |

**Supplemental Table 4:**
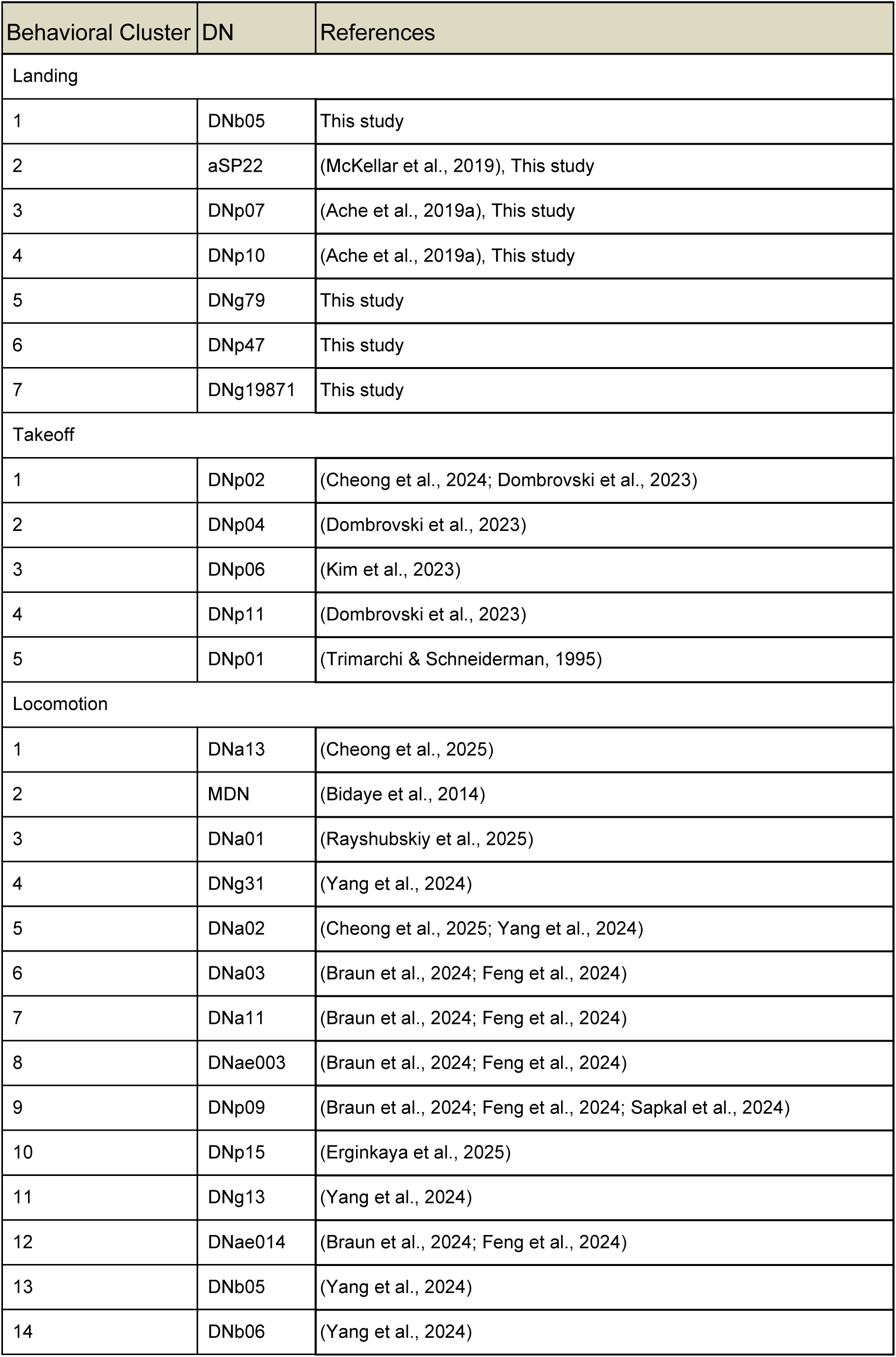
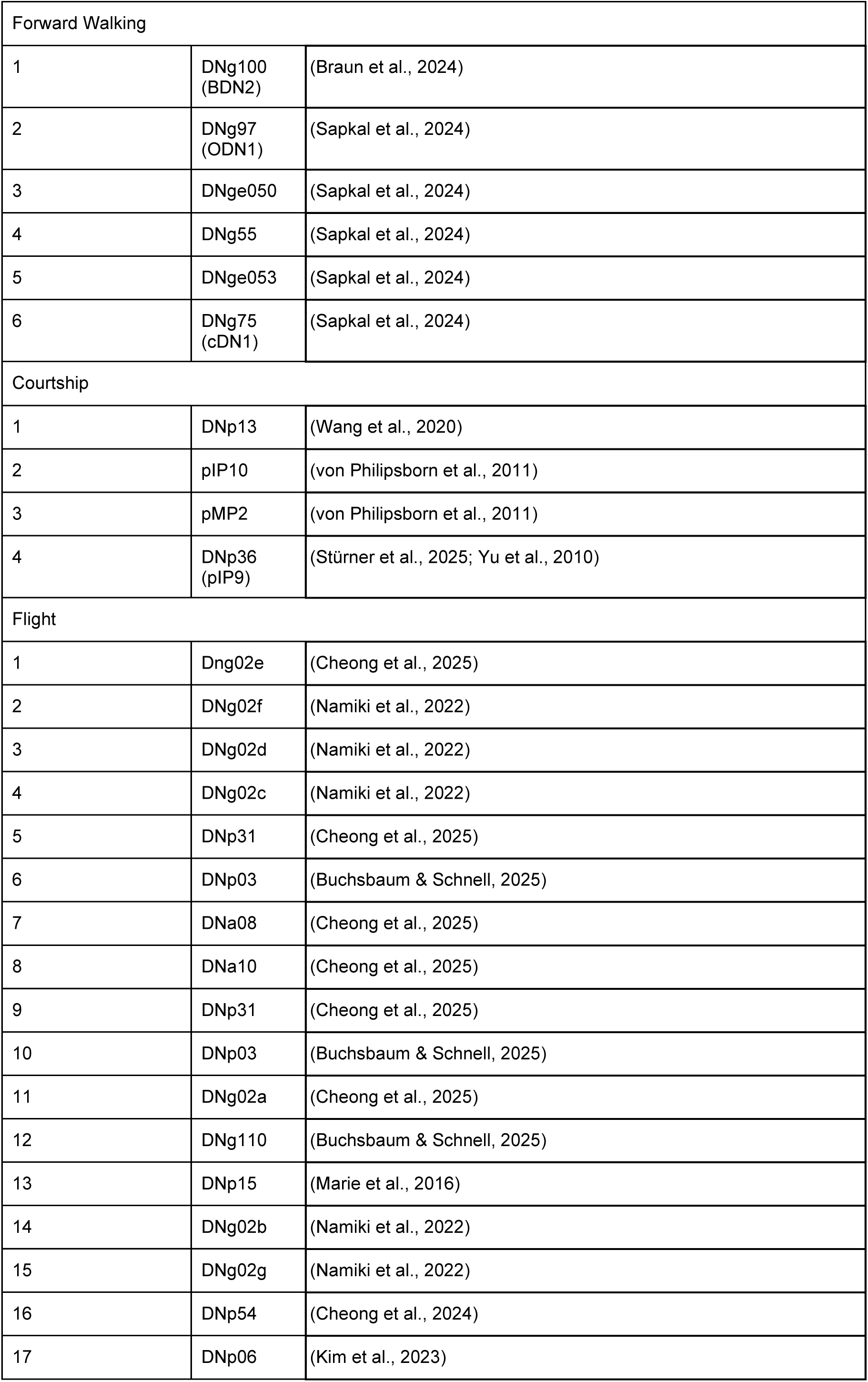
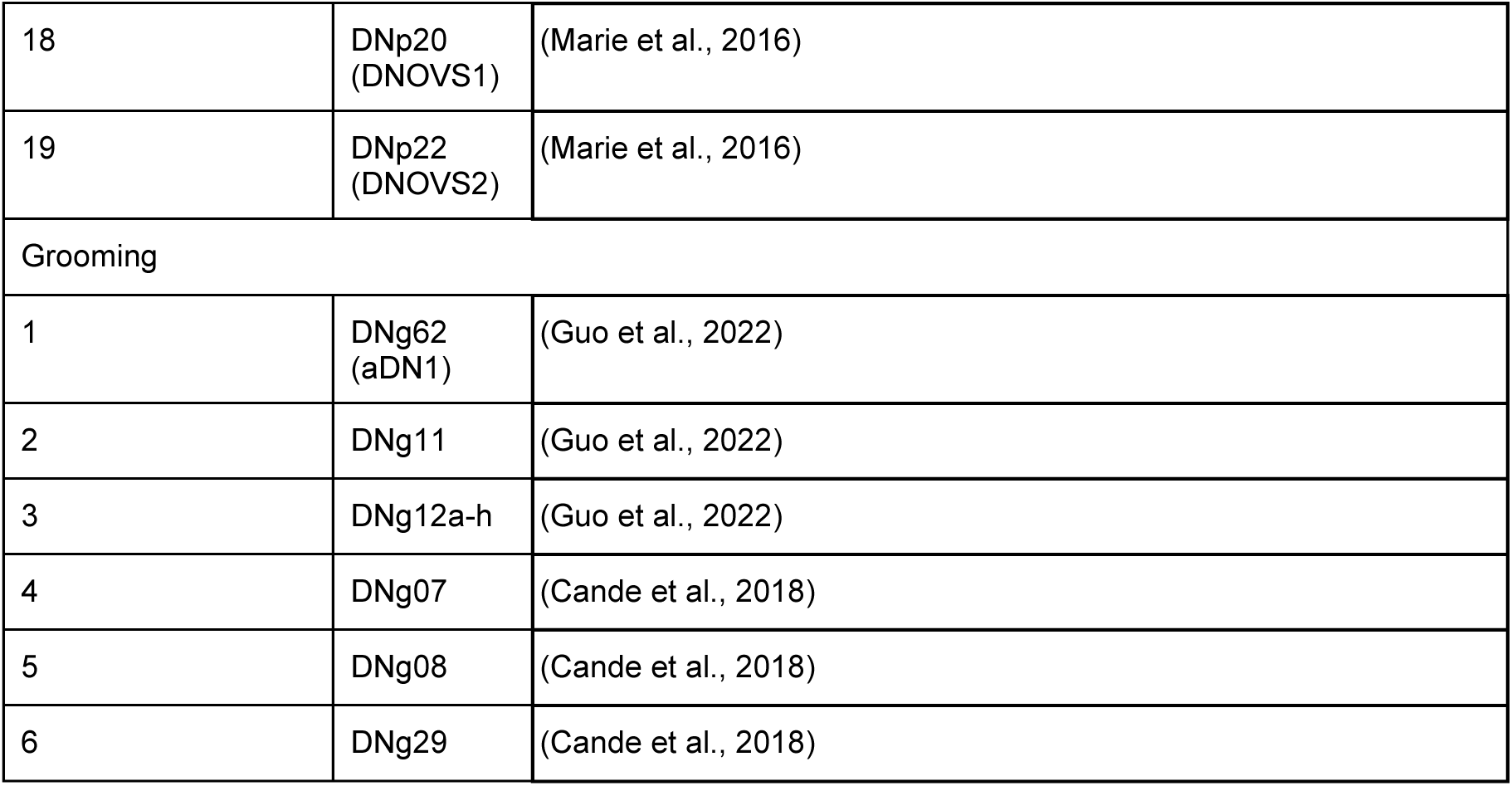
References for behaviorally functional units and are known to actuate similar behaviors (sorted into the same cluster in Fig. 5B).

| Behavioral Cluster | DN | References |
| --- | --- | --- |
| Landing |  |  |
| 1 | DNb05 | This study |
| 2 | aSP22 | (McKellar et al., 2019), This study |
| 3 | DNp07 | (Ache et al., 2019a), This study |
| 4 | DNp10 | (Ache et al., 2019a), This study |
| 5 | DNg79 | This study |
| 6 | DNp47 | This study |
| 7 | DNg19871 | This study |
| Takeoff |  |  |
| 1 | DNp02 | (Cheong et al., 2024; Dombrovski et al., 2023) |
| 2 | DNp04 | (Dombrovski et al., 2023) |
| 3 | DNp06 | (Kim et al., 2023) |
| 4 | DNp11 | (Dombrovski et al., 2023) |
| 5 | DNp01 | (Trimarchi & Schneiderman, 1995) |
| Locomotion |  |  |
| 1 | DNa13 | (Cheong et al., 2025) |
| 2 | MDN | (Bidaye et al., 2014) |
| 3 | DNa01 | (Rayshubskiy et al., 2025) |
| 4 | DNg31 | (Yang et al., 2024) |
| 5 | DNa02 | (Cheong et al., 2025; Yang et al., 2024) |
| 6 | DNa03 | (Braun et al., 2024; Feng et al., 2024) |
| 7 | DNa11 | (Braun et al., 2024; Feng et al., 2024) |
| 8 | DNae003 | (Braun et al., 2024; Feng et al., 2024) |
| 9 | DNp09 | (Braun et al., 2024; Feng et al., 2024; Sapkal et al., 2024) |
| 10 | DNp15 | (Erginkaya et al., 2025) |
| 11 | DNg13 | (Yang et al., 2024) |
| 12 | DNae014 | (Braun et al., 2024; Feng et al., 2024) |
| 13 | DNb05 | (Yang et al., 2024) |
| 14 | DNb06 | (Yang et al., 2024) |
| Forward Walking |  |  |
| 1 | DNg100<br>(BDN2) | (Braun et al., 2024) |
| 2 | DNg97<br>(ODN1) | (Sapkal et al., 2024) |
| 3 | DNge050 | (Sapkal et al., 2024) |
| 4 | DNg55 | (Sapkal et al., 2024) |
| 5 | DNge053 | (Sapkal et al., 2024) |
| 6 | DNg75<br>(cDN1) | (Sapkal et al., 2024) |
| Courtship |  |  |
| 1 | DNp13 | (Wang et al., 2020) |
| 2 | pIP10 | (von Philipsborn et al., 2011) |
| 3 | pMP2 | (von Philipsborn et al., 2011) |
| 4 | DNp36<br>(pIP9) | (Stürner et al., 2025; Yu et al., 2010) |
| Flight |  |  |
| 1 | Dng02e | (Cheong et al., 2025) |
| 2 | DNg02f | (Namiki et al., 2022) |
| 3 | DNg02d | (Namiki et al., 2022) |
| 4 | DNg02c | (Namiki et al., 2022) |
| 5 | DNp31 | (Cheong et al., 2025) |
| 6 | DNp03 | (Buchsbaum & Schnell, 2025) |
| 7 | DNa08 | (Cheong et al., 2025) |
| 8 | DNa10 | (Cheong et al., 2025) |
| 9 | DNp31 | (Cheong et al., 2025) |
| 10 | DNp03 | (Buchsbaum & Schnell, 2025) |
| 11 | DNg02a | (Cheong et al., 2025) |
| 12 | DNg110 | (Buchsbaum & Schnell, 2025) |
| 13 | DNp15 | (Marie et al., 2016) |
| 14 | DNg02b | (Namiki et al., 2022) |
| 15 | DNg02g | (Namiki et al., 2022) |
| 16 | DNp54 | (Cheong et al., 2024) |
| 17 | DNp06 | (Kim et al., 2023) |
| 18 | DNp20<br>(DNOVS1) | (Marie et al., 2016) |
| 19 | DNp22<br>(DNOVS2) | (Marie et al., 2016) |
| Grooming |  |  |
| 1 | DNg62<br>(aDN1) | (Guo et al., 2022) |
| 2 | DNg11 | (Guo et al., 2022) |
| 3 | DNg12a-h | (Guo et al., 2022) |
| 4 | DNg07 | (Cande et al., 2018) |
| 5 | DNg08 | (Cande et al., 2018) |
| 6 | DNg29 | (Cande et al., 2018) |

**Supplemental Table 5.** Identities of neurons described in this paper.

| Neuron name | Module membership | Male CNS IDs | FlyWire IDs |
| --- | --- | --- | --- |
| **DNg19871 (DNg06)** | Landing DN Ensemble | 217430<br>552858<br>19871<br>203146<br>29310 | DNg06b/c<br>720575940635222640,<br>720575940656144289,<br>720575940610348458,<br>720575940626088835,<br>720575940623491987,<br>720575940627021349 |
| **IN800073** | Core Takeoff Motor Assembly | 800147<br>800073 | N/A |
| **IN800383** | Core Takeoff Motor Assembly | 903429<br>800383 | N/A |
| **IN800898** | Core Takeoff Motor Assembly | 904246<br>800898 | N/A |
| **IN802944** | Core Takeoff Motor Assembly | 802944<br>904036 | N/A |
| **IN802751** | Core Takeoff Motor Assembly | 802751<br>802843 | N/A |
| **IN803304** | N/A | 803967<br>803864<br>803304 | N/A |
| **IN801053** | N/A | 801162<br>801053 | N/A |
| **DNg79a** | Landing DN Ensemble | 13978 |  |
| **DNg79b** | Landing DN Ensemble | 13649 |  |

## Supplemental Videos

**Video S1: Optogenetic activation of DNp07_1 during flight.** Overimposed tracked movements of the front-, middle-, and hind-leg tarsal tips, antennal basis, pronotum on the thorax, and abdominal tip. Red box, time of stimulation.

**Video S2: Optogenetic activation of DNp07_2 during flight.** Overimposed tracked movements of the front-, middle-, and hind-leg tarsal tips, antennal basis, pronotum on the thorax, and abdominal tip. Red box, time of stimulation.

**Video S3: Optogenetic activation of DNp10 during flight.** Overimposed tracked movements of the front-, middle-, and hind-leg tarsal tips, antennal basis, pronotum on the thorax, and abdominal tip. Red box, time of stimulation.

**Video S4: Optogenetic activation of DNp47 during flight.** Overimposed tracked movements of the front-, middle-, and hind-leg tarsal tips, antennal basis, pronotum on the thorax, and abdominal tip. Red box, time of stimulation.

**Video S5: Optogenetic activation of DNg79 during flight.** Overimposed tracked movements of the front-, middle-, and hind-leg tarsal tips, antennal basis, pronotum on the thorax, and abdominal tip. Red box, time of stimulation.

**Video S6: Optogenetic activation of DNb05 during flight.** Overimposed tracked movements of the front-, middle-, and hind-leg tarsal tips, antennal basis, pronotum on the thorax, and abdominal tip. Red box, time of stimulation.

**Video S7: Optogenetic activation of empty control flies during flight.** Overimposed tracked movements of the front-, middle-, and hind-leg tarsal tips, antennal basis, pronotum on the thorax, and abdominal tip. Red box, time of stimulation.

**Video S8: Optogenetic activation of GFP control flies during flight.** Overimposed tracked movements of the front-, middle-, and hind-leg tarsal tips, antennal basis, pronotum on the thorax, and abdominal tip. Red box, time of stimulation.

**Video S9: Optogenetic activation of DNp07_1 and DNp10 during walking.** Yellow box, closeup from one animal of DNp10 during wing flicking. White box, time of stimulation.

**Video S10: Optogenetic activation of LLPC2/3_1 during flight.** Overimposed tracked movements of the front-, middle-, and hind-leg tarsal tips, antennal basis, pronotum on the thorax, and abdominal tip. Red box, time of stimulation.

**Video S11: Optogenetic activation of LPLC4 during flight.** Overimposed tracked movements of the front-, middle-, and hind-leg tarsal tips, antennal basis, pronotum on the thorax, and abdominal tip. Red box, time of stimulation.

**Video S12: Optogenetic activation of LPLC2 during flight.** Overimposed tracked movements of the front-, middle-, and hind-leg tarsal tips, antennal basis, pronotum on the thorax, and abdominal tip. Red box, time of stimulation.

**Video S13: Optogenetic activation of LPLC1 during flight.** Overimposed tracked movements of the front-, middle-, and hind-leg tarsal tips, antennal basis, pronotum on the thorax, and abdominal tip. Red box, time of stimulation.

**Video S14: Optogenetic activation of LPT52 during flight.** Overimposed tracked movements of the front-, middle-, and hind-leg tarsal tips, antennal basis, pronotum on the thorax, and abdominal tip. Red box, time of stimulation.

**Video S15: Optogenetic activation of LPC2 during flight.** Overimposed tracked movements of the front-, middle-, and hind-leg tarsal tips, antennal basis, pronotum on the thorax, and abdominal tip. Red box, time of stimulation.

**Video S16: Optogenetic activation of LPC1 during flight.** Overimposed tracked movements of the front-, middle-, and hind-leg tarsal tips, antennal basis, pronotum on the thorax, and abdominal tip. Red box, time of stimulation.

**Video S17: Optogenetic activation of LC11 during flight.** Overimposed tracked movements of the front-, middle-, and hind-leg tarsal tips, antennal basis, pronotum on the thorax, and abdominal tip. Red box, time of stimulation.

**Video S18: Optogenetic activation of LC22 during flight.** Overimposed tracked movements of the front-, middle-, and hind-leg tarsal tips, antennal basis, pronotum on the thorax, and abdominal tip. Red box, time of stimulation.

**Video S19: Optogenetic activation of LLPC1_1 during flight.** Overimposed tracked movements of the front-, middle-, and hind-leg tarsal tips, antennal basis, pronotum on the thorax, and abdominal tip. Red box, time of stimulation.

**Video S20: Optogenetic activation of LLPC1_2 during flight.** Overimposed tracked movements of the front-, middle-, and hind-leg tarsal tips, antennal basis, pronotum on the thorax, and abdominal tip. Red box, time of stimulation.

**Video S21: Optogenetic activation of L1/L2 during flight.** Overimposed tracked movements of the front-, middle-, and hind-leg tarsal tips, antennal basis, pronotum on the thorax, and abdominal tip. Red box, time of stimulation.

